# A Nanoscale Jitterbug Transformer from DNA

**DOI:** 10.1101/2025.10.21.683790

**Authors:** Seongmin Seo, Alexander A. Swett, Mallikarjuna Reddy Kesama, Anirudh S. Madhvacharyula, Ruixin Li, Yancheng Du, Markus Eder, Friedrich C. Simmel, Jong Hyun Choi

## Abstract

Many viruses have intricate polyhedral shells capable of symmetric transformations in response to external stimuli to initiate payload release. Such deployable auxetic nanostructures are not available in the synthetic realm. Here we present a nanoscale Jitterbug transformer using DNA origami that can reconfigure its structure upon chemical and optical signals while maintaining a Poisson’s ratio of −1. By leveraging molecular dynamics simulations, we design the Jitterbug DNA to form a compact octahedron by storing elastic energy and spontaneously transition into an expanded cuboctahedron by releasing it. DNA transformers are explored like viruses that can create nanopores on lipid membranes and regulate payload release into vesicles. This work integrates programmable DNA self-assembly with free-energy-guided mechanical design, providing a pathway toward adaptive nanomaterials with potential in synthetic organelles and stimuli-responsive nanodevices.

## INTRODUCTION

Deployable auxetic materials can change geometry, switching from compact to expanded configurations, and vice versa, in response to external stimuli^1-3^. They exhibit unique mechanical behaviors with negative Poisson’s ratios (*v*)^4^ which are advantageous for applications in aerospace engineering^5^, soft robotics^6^, and biomedical devices^7^. Especially, materials with *v* = −1 show isotropic deformation, preserving geometric similarity and enabling force exertion and predictable changes in dimensions with the ability to stow into compact volumes.

Similarly in nature, viruses such as cowpea chlorotic mottle virus^8, 9^, brome mosaic virus^10^, and bacteriophage HK97^11^ undergo reversible capsid contraction-expansion in response to environmental cues including pH and ionic strength. These transformations scale pore size and permeability of lipid membranes while retaining symmetry, allowing for selective molecular exchange. Despite their considerable utility, the development of synthetic 3D deployable auxetics at the nanoscale has been hindered by the difficulties of constructing precise geometries and actuating at such small length scales.

Buckminster Fuller’s Jitterbug transformer^12^ is an ideal example of a deployable transformation. It transforms between octahedral and cuboctahedral configurations via a single kinematically constrained pathway^13^. Its three-fold symmetric reconfiguration results in a five-fold increase in volume^14^. These characteristics make the Jitterbug transformer a simple yet compelling model for construction at the nanoscale.

DNA origami enables nanometer-scale assembly with exceptional precision and programmability^15, 16^. This powerful approach has been used to construct a variety of systems such as nanoscale turbines^17^, ratchet motors^18^ and vesicle pincers^19^ rendering it as a powerful tool to construct stimuli-responsive reconfigurable devices. Static DNA polyhedra^20^ like icosahedron and dodecahedron were also constructed by proper strand routing and crossover placement using design tools like Cadnano^21^, Scadnano^22^ or automated workflows like Athena^23^ and Perdix^24^. However, deployable auxetics must be designed to ensure mechanical responses with symmetry. Proper design choices need to be made to bias their equilibrium configurations toward the deployed states based on desired functionalities. Their mechanical behavior may be tuned further by engineering structural components^25^, defects^26^, and chemical adducts^27, 28^.

Here we show a nanoscale, deployable Jitterbug transformer made of wireframe DNA origami via free-energy-guided mechanical design. Leveraging molecular dynamics (MD)^29^ and virtual-move Monte Carlo (VMMC)^30^ simulations, we tune its free energy landscape and mechanical response such that transformations are driven by storing and releasing elastic energy. Reversible reconfiguration is demonstrated in response to external stimuli like toehold-mediated strand displacement (TMSD)^31, 32^ and photocleavage. Furthermore, we harness the transformation capability into a bioinspired force actuator and nanoscale platform that regulates membrane permeability in synthetic vesicles, creating nanopores for controlled delivery of payloads.

## RESULTS

### Free-energy-guided mechanical design

A Jitterbug transforms between octahedral and cuboctahedral configurations by closing and opening six squares uniformly (Figs. 1a, S1, and S2). The opening distance of the squares is thus the key parameter that determines the configuration. Each of the eight triangular faces of the initial octahedron undergoes radial displacement and axial rotation about its surface normal, resulting in screw-like motion while retaining overall symmetry. We initially designed a Jitterbug transformer using wireframe DNA with 6-helix bundles (6HB) on a honeycomb lattice to ensure edge rigidity during reconfiguration. Each edge is approximately 28 nm long, and the overall structure size is ∼40 nm (Fig. 1b-c). Figure 1c shows MD structures of the two final states computed using coarse-grained oxDNA simulations^29^ (see SI S3.4 for computational details).

**Figure 1.**
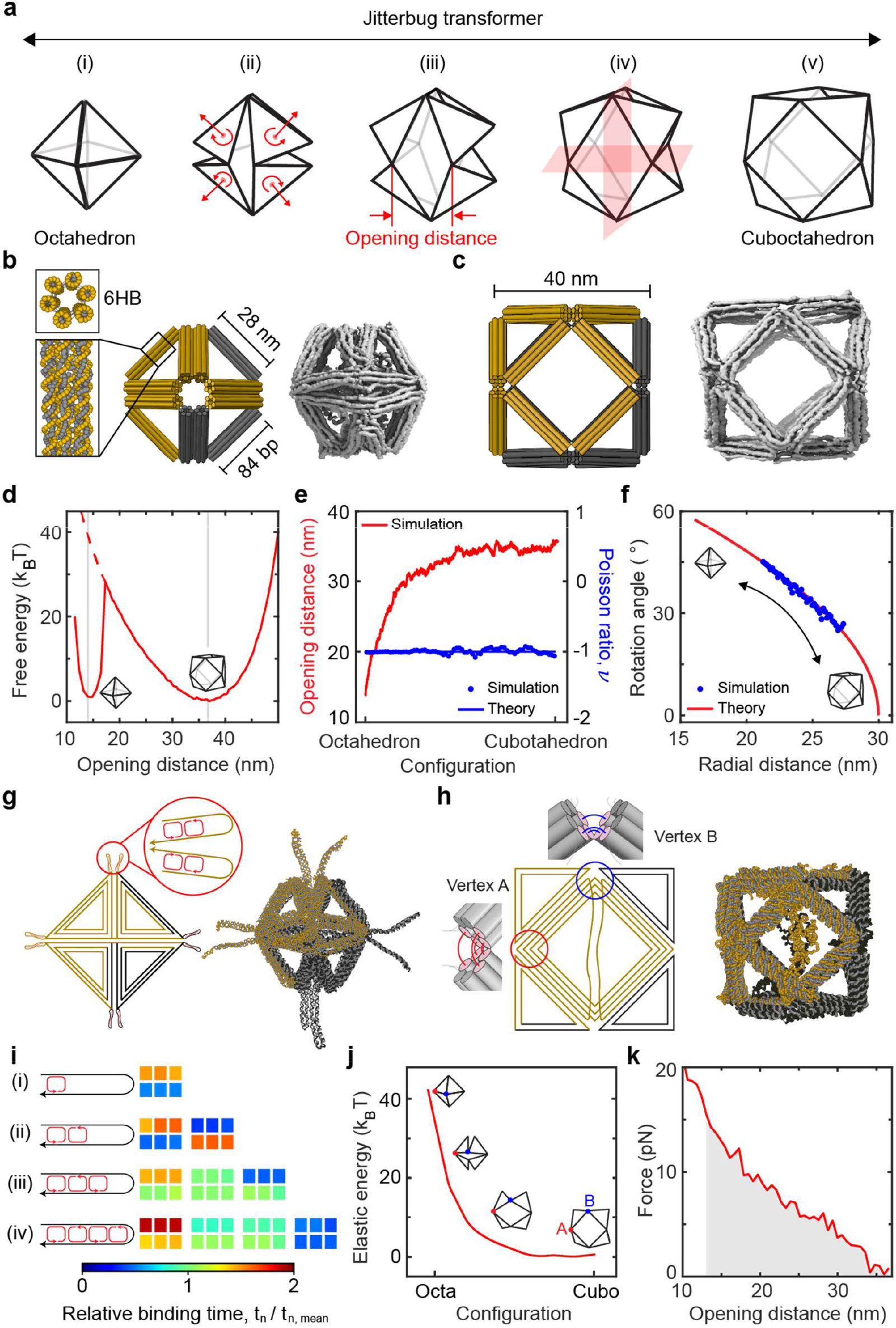
Free-energy-guided mechanical design of a DNA transformer. **a**, A Jitterbug switching between compact octahedral and expanded cuboctahedral configurations. During the transformation, six square faces open or close uniformly, while eight triangles displace radially and rotate about their normal axes. **b** and **c**, Design scheme of a DNA Jitterbug made of 28-nm-long edges consisting of 6HB on a honeycomb lattice. Static MD structures in **b** octahedral and **c** cuboctahedral states. **d**, Free energy landscape with the opening distance as reaction coordinate. Cuboctahedral profile (including the dashed line) from umbrella sampling simulations reveals an energy gap of ∼40 k_B_T per square between closed and open conformations marked by vertical lines in light gray color. VMMC-simulated energy profile (left) of an octahedron which may be stabilized by DNA base-pairs. **e**, Opening distance during a spontaneous transition from octahedral to cuboctahedral states, after releasing the stabilization energy. The Poisson ratio maintains the theoretical value of −1, confirming isotropic deformation. **f**, Transformation kinematics analysis between radial displacement and rotation angle. Portions of the initial and final transformation phases are inaccessible due to the finite edge thickness. **g**, Octahedral schematic and MD model stabilized by short jack edges. **h**, Front view and MD conformation of cuboctahedron with unparied scaffold segments in the square face. Vertices A and B are four and three 10-nt ssDNA connections, respectively. **i**, Relative binding kinetics from DNAfold simulations for (i) one, (ii) two, (iii) three, and (iv) four staples per jack edge. **j**, Elastic energy distribution during reconfiguration, with the majority of strain energy (∼40 k_B_T) localized at vertex A regions. **k**, Distribution of force required to maintain instantaneous conformation. Integration of all forces (gray shade) yields ∼42 k_B_T, consistent with free energy predictions.

The deployability of the Jitterbug was designed by tuning two main components of its free energy, (i) inherent structural response and (ii) chemical stabilization, which enable reconfigurations into the open and closed states, respectively. The structural free energy was estimated using umbrella sampling simulations^33, 34^, with the average opening distance of the six squares as the reaction coordinate. As shown in Fig. 1d, the structure has a minimum at around 37 nm revealing a natural bias towards the open state. A closer look at the dotted curve indicates that the difference in energy between the open and closed states (marked by light gray lines) is approximately 40 k_B_T per square. This is the amount of energy needed to stabilize the octahedral configuration, which may be provided via DNA base-pairing. The minimum number of DNA base-pairs (bp) required for a stable octahedral state was calculated using NUPACK^35^. The calculation suggests that at least 20 bp are necessary. The free energy of the closed state is a summation of structural and stabilization energies. The latter was characterized as a function of opening distance using VMMC simulations^30^ in oxDNA. Stabilization via base-pairing enables a minimum in the closed state (the left well in Fig. 1d), negating the effect of increased strain in the structure. Both wells are nearly equal in depth since we choose the theoretical minimum number of base-pairs required for comparison. Upon removal of stabilization (i.e., with sufficient activation energy to overcome the barrier), the structure is expected to shift to its natural open state at the free energy minimum.

We tested the structural transformation numerically by removing base-pairs, which indeed triggered a spontaneous transition from octahedral to cuboctahedral configurations, with the opening distance increasing from approximately 13 to 37 nm (Fig. 1e). Poisson’s ratio, averaged across three orthogonal planes (Figs. 1a(iv) and S7), is approximately −1 during the process, agreeing with the theoretical estimates (see SI S2.1, S3.2-3.3). We also analyzed the rotation of the triangular faces for their radial displacement from the origin. The rotation angle vs. the radial distance from the MD simulation closely matches theoretical predictions^13^ (Fig. 1f). However, portions of the initial and final transformation phases are inaccessible due to the finite edge thickness of the DNA construct.

The requisite energy for stabilizing the octahedral configuration can be supplied via adjustable DNA ‘jack’ mechanisms, conceptually analogous to mechanical jacks that regulate structure through controlled length modulation^25, 36^. We included jack edges in our design by placing them diagonally in an alternating fashion across the six square faces. This arrangement is expected to enable switching of the Jitterbug between two configurations by inserting a set of staples for short and long jacks. A key aspect here is that since the cuboctahedron is designed as the free energy minimum, the configuration should be achieved regardless of jack staples. Our scaffold routing between 6HB edges in the square faces has two topological configurations: vertices A and B having four and three 10-nt connections, respectively (Fig. 1h). During reconfiguration vertex A undergoes conformational changes while vertex B remains nearly unchanged.

Mesoscopic simulations using a simplified jack design reconciled theoretical energy estimates with experimental feasibility. We estimated the normalized binding times between scaffold and jack staples for varying numbers of staples using DNAfold simulations^37^ (Fig. 1i). As more staples were added, the staples closest to the middle of the scaffold bound first while those located at the ends of the scaffold bound later due to the high entropic cost of bringing the scaffold segments together. Since this binding pattern correlates with experimental folding yields, we implemented four short jack staples to reliably fold each scaffold segment and thus stabilize the overall octahedral configuration.

In our Jitterbug design, joints at vertices A and B behave differently but together contribute to the overall response. Using a simple model consisting of a square unit with triangles, we simulated both connection types with repulsion planes in the oxDNA platform to enforce symmetric transformation constraints (Figs. 1j and S10). The estimated difference in free energy between the open and closed squares is approximately 40 k_B_T. This is the same as the cumulative estimate from Fig. 1d, indicating that the structural response is a summation of contributions from individual joints. That is, each square can open and close independently of the others. Notably, most of this elastic energy is localized at vertex A regions, providing insight into the mechanical stress distribution within the structure.

Lastly, we calculated the force required to maintain different configurations during the octahedron-to-cuboctahedron transition as a function of opening distance to estimate the mechanical work required throughout the transformation. For example, about 15 pN is needed to stabilize the octahedral state, whereas the cuboctahedral state requires negligible force due to its equilibrium configuration, as shown in Fig. 1k. Integration of the force profile over the distance yields ∼42 k_B_T (shaded area). This value demonstrates excellent agreement with stored elastic energy and free energy calculations, validating our free-energy-guided mechanical design.

### Jitterbug construction

The size of the Jitterbug requires using two scaffolds in a single origami structure: M13mp18 (7249-nt) and p9072 (termed 9k scaffold)^38^ colored gold and black, respectively, in Figs. 1 and 2. The origami design consists of 7k and 9k portions that are connected by linker staples (Fig. 2a). The structures were synthesized using one-pot assembly, where all staple strands and both scaffolds were mixed to form the initial cuboctahedra without jack staples (Fig. 2b(i)). Staple sequences were optimized computationally to minimize sequence overlap and reduce unintended hybridization (SI S3.1).

**Figure 2.**
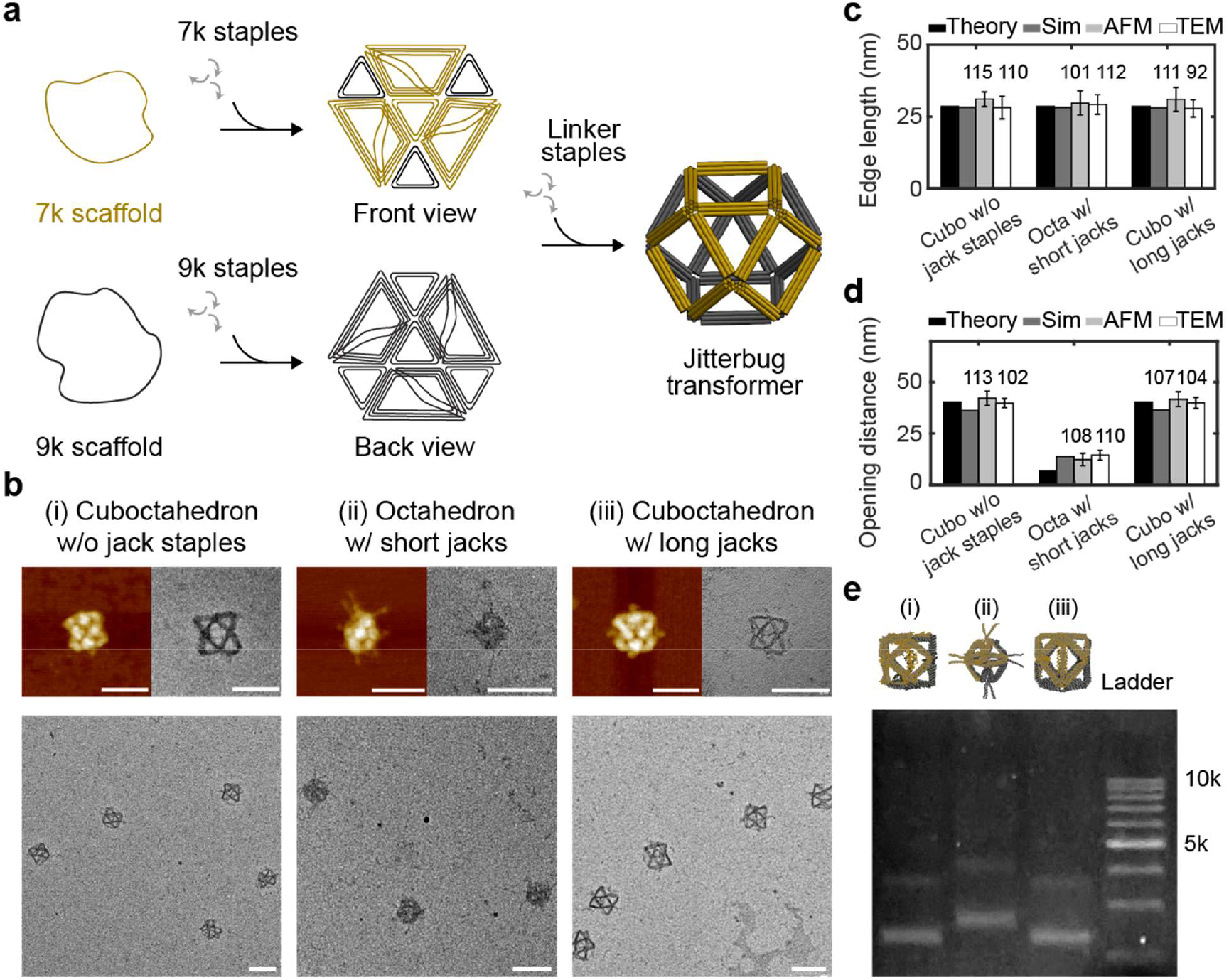
Construction and characterization of a DNA Jitterbug transformer. **a**, Design schematic of a DNA Jitterbug transformer assembled using two scaffolds: M13mp18 (7k scaffold, gold) and p9072 (9k scaffold, black) connected via linker staples. **b**, Three distinct configurations characterized with AFM and TEM imaging. (i) initial cuboctahedra without jack staples, (ii) octahedra with short jacks, and (iii) cuboctahedra with long jacks. Scale bars: 100 nm. **c** and **d**, Comparison of **c** edge lengths and **d** opening distances from theoretical models, MD simulations, AFM, and TEM images. **e**, Agarose gel electrophoresis analysis of the three configurations.

Octahedral and cuboctahedral configurations were built by introducing distinct types of jack staples to the initial cuboctahedron (SI Fig. S4). Short jacks closed the squares diagonally, contracting the structure into an octahedron. Long jacks kept the squares open, preserving the cuboctahedral conformation. AFM and TEM images in Fig. 2b confirm these configurations. Compact octahedra are clearly observed with several jacks hanging loose, as in the MD structure. Cuboctahedra with and without jacks appear nearly identical, distinguished solely by the inclusion of long jacks in the square faces, confirming that the cuboctahedral state is thermodynamically the most stable as indicated by the free energy profile.

Consistency across different configurations was examined by quantitative comparison of edge lengths and opening distances. The edge lengths measured from MD, AFM, and TEM images are about 28 nm, agreeing with the theoretical design (Fig. 2c). Opening distances are also consistent although they are slightly larger than the designed value in the octahedral state (Fig. 2d). Gel electrophoresis in Fig. 2e shows that both cuboctahedra migrate faster in the gel than the octahedral Jitterbug^39^.

### Jitterbug transformation

We next demonstrated the reversible Jitterbug transformation via adjustable jack edges using TMSD and jack staples. Figure 3a-b illustrates our transformation strategy, beginning with (i) cuboctahedron without jack staples, where two unpaired scaffold segments are pre-positioned diagonally across each of the six squares to serve as binding sites for jack staples in subsequent steps. Staples for short jacks (red) contract the squares, transforming the structure into (ii) an octahedron. The introduction of ‘releasers’ remove short jack staples via TMSD, returning to (i) a cuboctahedron. Likewise, long jack staples (blue) form extended jacks (iii), preserving the cuboctahedral geometry. TMSD using appropriate releaser strands can revert to (i) the cuboctahedron, as demonstrated in Fig. 3c.

**Figure 3.**
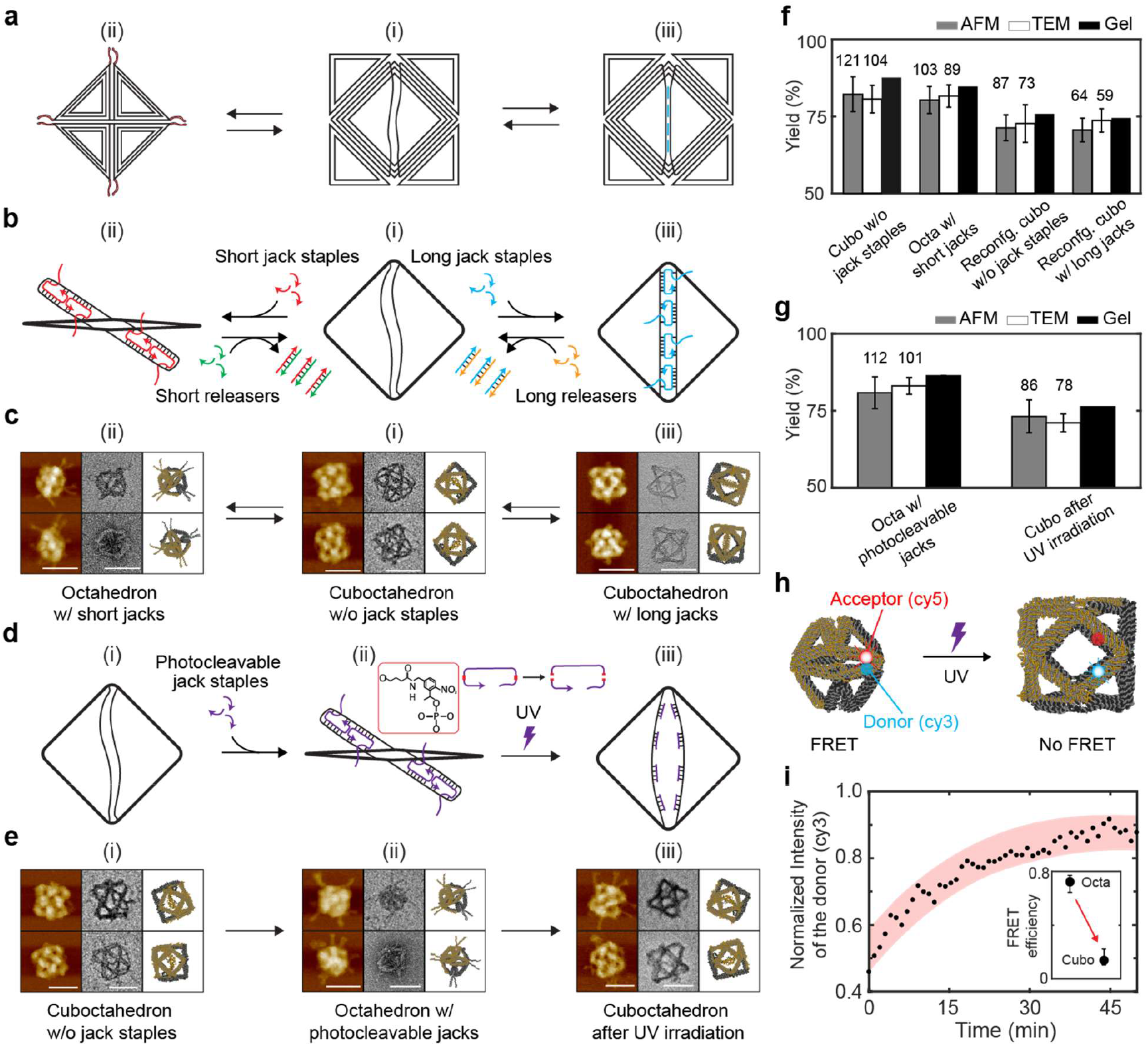
Reconfiguration via elastic energy storage and release. **a** and **b**, Jack-mediated transformation mechanism. Starting from (i) initial cuboctahedron with unpaired scaffold segments across square openings, two pathways are possible: (ii) an octahedron is formed by shutting the opening with short jack staples (red), while long jacks (blue) maintain the square opening preserving (iii) cuboctahedral geometry. Reversible reconfiguration is initiated by TMSD that removes jack staples with releaser strands. Subsequent introduction of another set of jack staples completes reconfiguration, enabling transitions between all three configurations. **c**, AFM, TEM, and MD results showing the three structural states achieved through controlled jack edge modulation. Scale bars: 100 nm. **d**, UV-responsive design. Photocleavable jack staples (purple) are incorporated into the initial cuboctahedron, with red indicating UV-triggered cleavage sites. **e**, AFM, TEM, and MD data showing transformation from (i) initial cuboctahedron to (ii) octahedron using photocleavable short jacks and eventually to (iii) cuboctahedron after UV exposure. **f** and **g**, Experimental yields of three configurations using AFM, TEM, and agarose gel electrophoresis. The numbers of DNA structures used for yield estimation are included in each panel. **h** and **i**, FRET measurement of reconfiguration kinetics, showing a complete transformation over 30 minutes.

To achieve convenient reconfiguration without requiring additional annealing steps (with releasers) and generating chemical waste, we engineered jack staples with photocleavable moieties^27^ such that long-wavelength UV (UVA) light will trigger spontaneous transformation from octahedral to cuboctahedral configurations (Fig. 3d and SI S1.4). Figure 3e confirms this approach, validating elastic energy storage via DNA base-pairing and the release by external light irradiation.

Figures 3f-g and S29 present experimental yields of the Jitterbug structures. The initial cuboctahedra, octahedra, and cuboctahedra with extended jacks all show approximately 83% yields, while reconfigured structures are slightly lower (also see SI S8). The yields calculated from gel electrophoresis were slightly higher since partially defective structures migrated similar to intact origami. We also measured reconfiguration kinetics upon UV irradiation using Förster resonance energy transfer (FRET). Cy3 (donor) and Cy5 (acceptor) dyes are placed on adjacent edges in a square (Fig. 3h) such that transformation will change the FRET signals. In the octahedral configuration, the proximity between the donor and the acceptor quenches the donor emission. During the transition into the cuboctahedron, the donor signal restores about 90% (the FRET efficiency decreases from nearly 80 to 20%) as shown in Figs. 3i and S30, suggesting that UV irradiation over 30 minutes is sufficient for complete reconfiguration.

### Light-induced nanopores on synthetic vesicles

Next, we explored the utility of the DNA transformer as a nanoscale actuator to regulate permeability of lipid membranes. The Jitterbug generates elastic energy that exceeds the minimum energy required to form nanopores in lipid membranes (∼30 k_B_T)^40, 41^. DNA reconfiguration produces an external force against lateral tension on the membrane, opening a nanopore until the force equilibrium is reached. To achieve this, we anchored the DNA nanostructures onto the membrane and induced their transformation via UV irradiation.

The experimental workflow consists of four steps (Fig. 4a). (i) Giant vesicles decorated with biotin are immobilized on a biotinylated BSA-functionalized channel surface via biotin-streptavidin chemistry. (ii) The DNA nanostructures are labeled with Cy5 dyes and cholesterol groups for fluorescence tracking and stable membrane anchoring, respectively (Fig. 4b). Octahedral structures are bound to the membrane. (iii) Upon UV exposure, photocleavable jack staples are cleaved, transforming the Jitterbug and creating transient nanopores. This reconfiguration allows dextran-rhodamine (DR) in the surroundings to diffuse into the vesicle through the pores. (iv) With UV light off, the nanopores are resealed through elastic relaxation of the membrane^41, 42^, retaining fluorescent DR molecules inside the vesicle.

**Figure 4.**
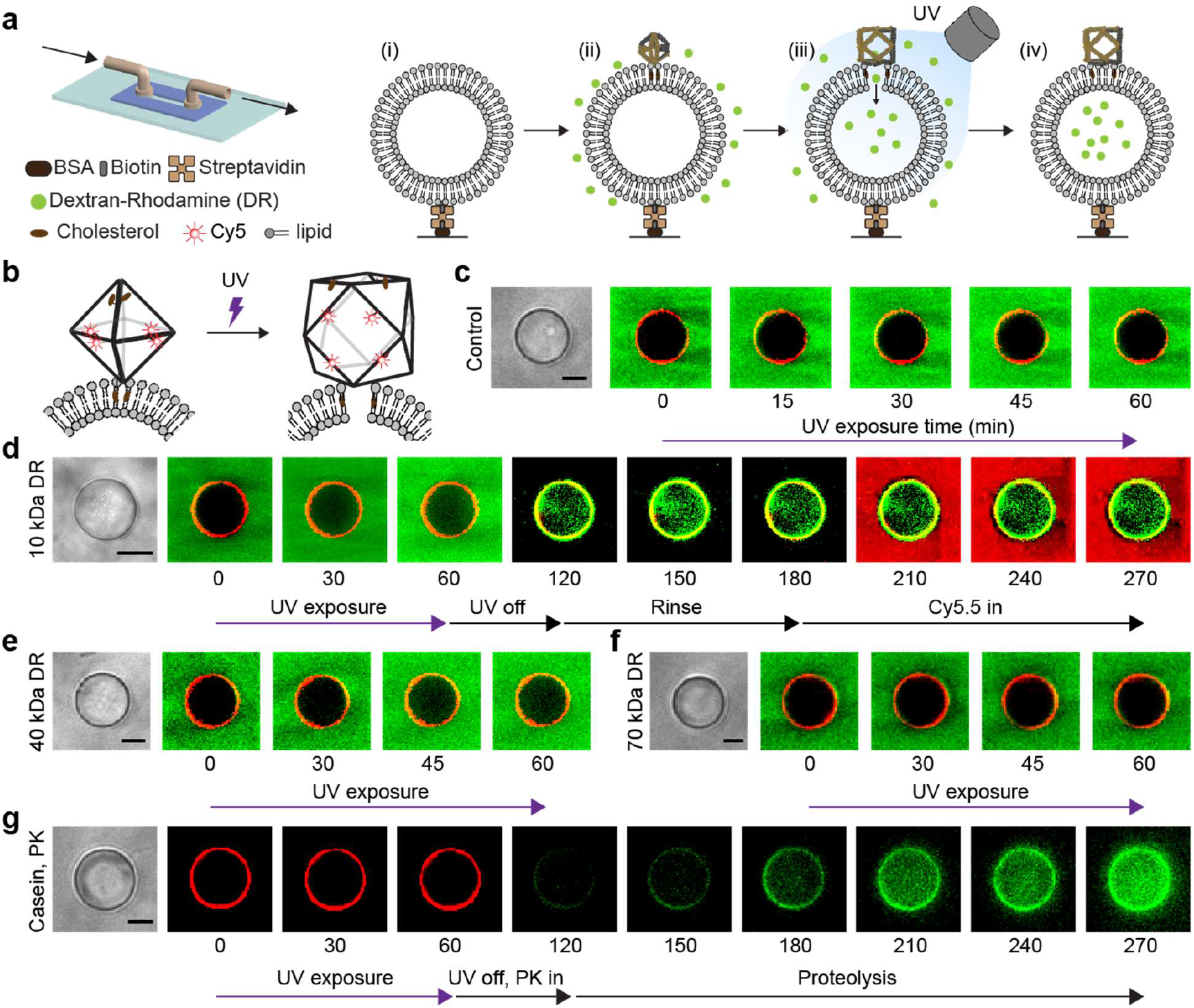
UV-triggered nanopore formation and molecular transport in synthetic vesicles. **a**, Schematics of a microfluidic imaging chamber and experimental workflow. (i) Immobilization of a biotin-functionalized vesicle on a BSA-decorated surface via biotin-streptavidin binding. (ii) Cy5-labeled Jitterbug binding to lipid membrane and introduction of dextran-rhodamine (DR). (iii) UV-triggered DNA transformation creating transient nanopores through which DR enters the vesicle. (iv) Membrane resealing after UV light is turned off, prohibiting molecular transport. **b**, The DNA transformer with photocleavable jacks and cholesterol moieties undergoes a UV-triggered octahedral-to-cuboctahedral transition, creating transient membrane pores for controlled molecular passage. **c**, Control experiment using non-photocleavable DNA. Despite UV irradiation, the vesicle with Cy5-labeled transformers (red) shows no DR influx (green), confirming that nanopore formation requires Jitterbug transformation. **d**, During 60-minute UV exposure, 10 kDa DR (green) enters the vesicle through nanopores created by Jitterbug transformation. Post-UV recovery shows complete pore closure with no DR leakage or exclusion of externally added Cy5.5 dyes (red), confirming a resealing of lipid membrane. **e** and **f**, Size-selective molecular transport: 40 kDa DR penetrates the vesicle with UV exposure, while 70 kDa DR is excluded. **g**, Jitterbug (red) transformation allows PK to enter the vesicle preloaded with casein-Bodipy substrate. Enzymatic proteolysis activates Bodipy fluorescence (green), confirming PK entry and its function. Scale bars in brightfield images: 10 μm. See Figs. S32-S38 for additional data and analysis.

UV-triggered nanopores were studied in a series of experiments using several molecular probes. Cy5 fluorescence along the vesicle membrane shows the presence of DNA Jitterbug (red, Figs. 4c-g). Control experiments using non-photocleavable staples show that no DR entered the vesicle during UV irradiation, verifying that nanopores were not formed (Figs. 4c and S32). In contrast, photoactivatable transformers can create nanopores under UV light. During 60 minutes of UV exposure, the green fluorescence signal increases inside the vesicle, providing direct evidence of 10 kDa DR entry (Figs. 4d, S33, and S37). Notably, the membrane restored after the UV lamp was turned off; 60 minutes post-irradiation, the absence of DR leakage and exclusion of externally added Cy5.5 confirm a complete pore closure. To characterize nanopore size selectivity, we evaluated the diffusion of larger probes (40 and 70 kDa DR). While 40 kDa DR signal was observed inside the vesicle, 70 kDa DR was excluded, indicating size-dependent permeability (Figs. 4e-f, S34-S36). From this observation, we estimate the pore diameter to be approximately 11-15 nm^43^.

We also examined enzymatic assays inside vesicles preloaded with casein-Bodipy (CB) substrate. This fluorogenic substrate remains quenched until proteolytic cleavage activates green fluorescence of the Bodipy dye (Figs. 4g and S38). Initially, CB encapsulated in the vesicle shows little to no emission signals, while Cy5 fluorescence confirms the Jitterbug conjugation on the membrane. UV activation creates nanopores via Jitterbug transformation that allows external proteinase K (PK) to enter the vesicle and digest the substrate, gradually restoring the fluorescence of the quenched Bodipy. The observation indicates molecular transport through UV-induced nanopores and enzymatic activities. This UV-responsive system combined with membrane dynamics presents a versatile platform for externally regulating membrane permeability.

### Programmable release of encapsulated payloads into vesicles

Leveraging the deployability of our DNA Jitterbug, we investigated its feasibility as a programmable nanosystem that delivers payload enzymes to initiate proteolysis inside synthetic vesicles. The Jitterbug maintains a consistent overall size of ∼40 nm, while increasing its internal volume drastically during deployment. We functionalized 5-nm-diameter gold nanoparticles (AuNPs) with photocleavable oligonucleotides for dual purposes: encapsulation via base-pairing^44^ and UV-triggered release from the Jitterbug. Either PK or trypsin (TRP) were used as enzyme payloads which were conjugated onto AuNPs (Figs. 5a-b). We encapsulated Cy5.5-labeled Au-enzyme complexes within initial cuboctahedra which were subsequently converted into octahedra using a mixture of photocleavable and regular jack staples. Upon UV irradiation, only one square of the DNA transformer interfacing the vesicle opened, enabling directional payload release into the cavity while preventing unintended release into the free solution. Opening all six squares would make the cargo delivery less efficient (Fig. S43).

**Figure 5.**
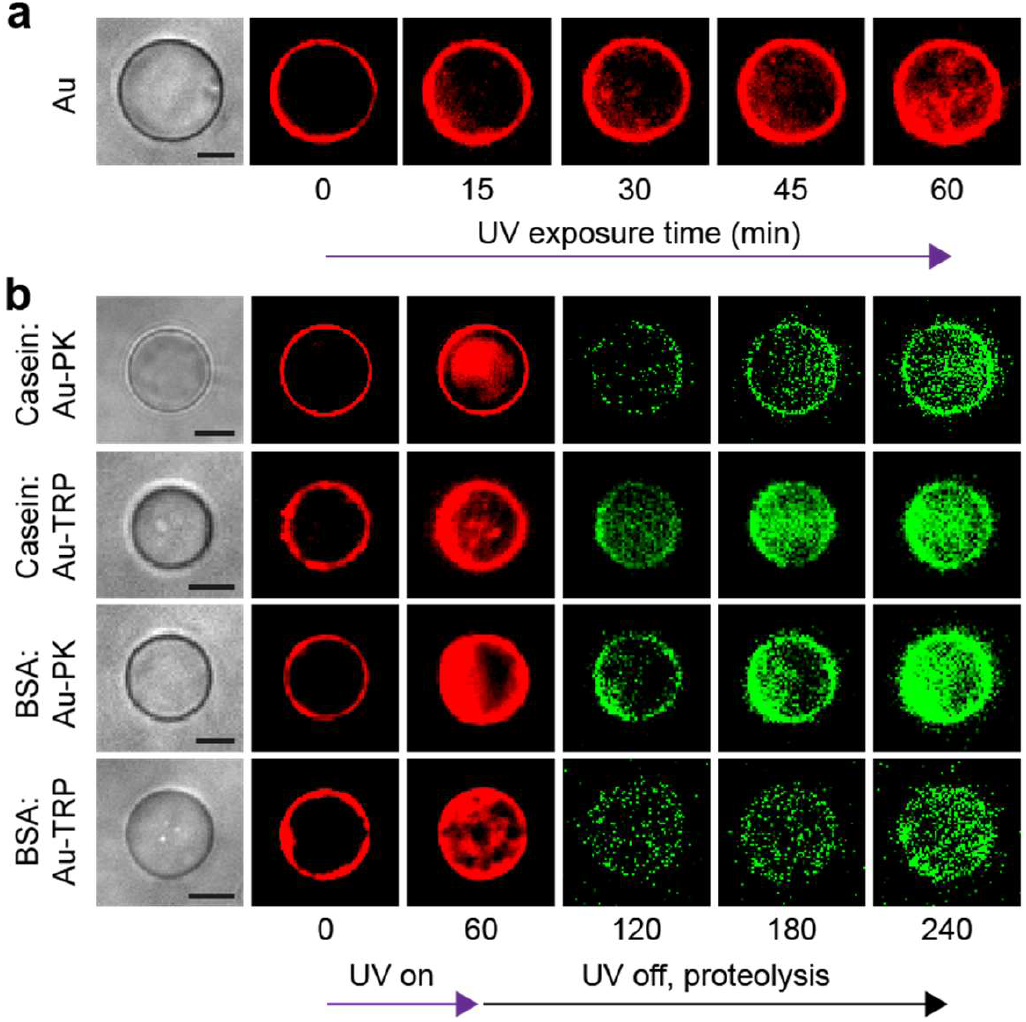
Programmable delivery of enzyme payload via Jitterbug transformation. **a**, UV-activated entry of Cy5.5-labeled AuNPs (red) into the vesicle cavity. Initially, Cy5-labeled DNA transformers (red) encapsulating AuNPs bind to vesicle surfaces. During 60-minutes UV exposure, AuNPs (red) enter vesicles through nanopores created by Jitterbug transformation. **b**, Proteolysis with several enzyme-substrate pairs (enzymes: PK and TRP, self-quenched (green) fluorescent substrates: casein-Bodipy or BSA-TMR). Red signals at 60 minutes indicate the entry of Au-enzyme complexes via UV-triggered DNA transformation. Following UV exposure, enzymatic activity increases green fluorescence intensity (Bodipy and TMR) of substrates inside vesicles over time, confirming enzyme (PK and TRP) entry and function. Scale bars: 10 μm.

We first tested the entry of the AuNPs by UV activation. Cy5 fluorescence along the vesicle boundary confirms the conjugation of the Jitterbug (containing Cy5.5-labeled AuNPs) on the membrane (Fig. 5a). We observed a gradual increase of red fluorescence in the cavity during UV exposure, indicating successful transport of AuNPs into the vesicle through nanopores created during the DNA transformation. We then investigated various enzyme-substrate combinations (Figs. 5b and S39-S42). We employed Bodipy-conjugated casein and TMR-tagged BSA as self-quenched fluorescent substrates pre-loaded within vesicles, while trypsin and PK were conjugated to AuNPs. Following 60 minutes of UV exposure, DNA transformation enabled the entry of Au-enzyme complexes into vesicles and subsequent enzymatic proteolysis. Green fluorescence gradually increased in the vesicle, indicating increased digestion of the substrates. Proteolysis within the vesicle microenvironment was influenced by several factors including substrate distribution, enzyme loading, and enzyme-substrate affinity. These results demonstrate that the DNA Jitterbug can mediate enzymatic reactions within vesicles, which may be further developed for synthetic organelles^45, 46^.

## DISCUSSION

This work introduced free-energy-informed design strategies for nanoscale metamaterials capable of deployable auxetic transformation. Transition between two topological states was achieved by stabilizing the compact octahedral configuration via insertion of DNA base-pairs into jack edges. Expanded cuboctahedra were designed as the global minimum in free energy landscape, which formed naturally when the elastic energy was released or even when long jacks were implemented. Our framework opens new avenues for designing increasingly complex and adaptive DNA materials by integrating mechanical energy, structural responses, and transformation mechanisms with computational models. For example, this design approach may be extended to engineer transformable multistable nanosystems with three or more topological states that can respond selectively to distinct external stimuli.

Beyond reconfiguration, the DNA transformer serves as a nanoscale actuator capable of generating mechanical energy that exceeds that of natural biomolecular machines. Jitterbug DNA can store and release up to approximately 240 k_B_T of elastic energy (or nearly 100 pN), while motor protein kinesin generates ∼10 k_B_T per step (about 5 pN over 8 nm)^47^. This demonstrates the potential of DNA metamaterials as nanomachines, bridging biomolecular energy scales with engineering precision.

Inspired by viral capsid dynamics, the DNA transformer was exploited to create transient nanopores in synthetic vesicles, enabling a programmable release of payload enzymes into the cavity. The modularity of DNA origami allows for variable payloads (and substrates) and control over biochemical processes, making it useful for applications such as proteolysis, signal transduction, or reaction cascades. This ability could be extended to treat bacterial biofilms^48^. To render this platform applicable to living systems, several mechanisms must be improved in the future, including high-affinity binding to the membrane, faster reconfiguration kinetics, and response to alternative stimuli like pH.

## ACKNOWLEDGMENTS

The authors thank Drs. Dietz, Honemann, and Pinner at Technical University of Munich for providing the p9072 scaffold. This work was financially supported by the U.S. Department of Energy, Office of Science, Basic Energy Sciences under award no. DE-SC0020673. J.H.C. acknowledges a Friedrich Wilhelm Bessel Research Award from the Alexander von Humboldt Foundation.

## AUTHOR CONTRIBUTIONS

R.L., F.C.S., and J.H.C. conceived the idea; S.S., A.A.S., M.R.K., Y.D., and M.D. performed the experiments; A.S.M. and R.L. conducted the numerical simulations; S.S., A.A.S., M.R.K., A.S.M., F.C.S., and J.H.C. analyzed the data. All authors participated in writing the manuscript.

## DATA AVAILABILITY

The datasets generated during and/or analyzed during the current study are available from the corresponding author on reasonable request.

## References

1. Lakes, R., Foam structures with a negative Poisson’s ratio. Science 1987, 235, 1038–1040.

2. Evans, K. E.; Alderson, A., Auxetic materials: functional materials and structures from lateral thinking! Advanced materials 2000, 12, 617–628.

3. Berger, J.; Wadley, H.; McMeeking, R., Mechanical metamaterials at the theoretical limit of isotropic elastic stiffness. Nature 2017, 543, 533–537.

4. Chan, N.; Evans, K., Indentation resilience of conventional and auxetic foams. Journal of Cellular Plastics 1998, 34, 231–260.

5. Alderson, A.; Alderson, K., Auxetic materials. Proceedings of the Institution of Mechanical Engineers, Part G: Journal of Aerospace Engineering 2007, 221, 565–575.

6. Rafsanjani, A.; Bertoldi, K.; Studart, A. R., Programming soft robots with flexible mechanical metamaterials. Science Robotics 2019, 4, eaav7874.

7. Veerabagu, U.; Palza, H.; Quero, F., Auxetic polymer-based mechanical metamaterials for biomedical applications. ACS Biomaterials Science and Engineering 2022, 8, 2798–2824.

8. Speir, J. A.; Munshi, S.; Wang, G.; Baker, T. S.; Johnson, J. E., Structures of the native and swollen forms of cowpea chlorotic mottle virus determined by X-ray crystallography and cryo-electron microscopy. Structure 1995, 3, 63–78.

9. Tama, F.; Brooks III, C. L., The mechanism and pathway of pH induced swelling in cowpea chlorotic mottle virus. Journal of Molecular Biology 2002, 318, 733–747.

10. Lucas, R. W.; Larson, S. B.; McPherson, A., The crystallographic structure of brome mosaic virus. Journal of Molecular Biology 2002, 317, 95–108.

11. Hendrix, R. W.; Johnson, J. E., Bacteriophage HK97 capsid assembly and maturation. Advances in Experimental Medicine and Biology 2012, 726, 351–363.

12. Fuller, R. B., Synergetics: explorations in the geometry of thinking. Macmillan: 1975.

13. Verheyen, H. F., The complete set of Jitterbug transformers and the analysis of their motion. Computers Mathematics with Applications 1989, 17, 203–250.

14. Gray, R. W., The Jitterbug motion. http://www.rwlllY.Projects.comlrbfnotes/toc.html: 2002.

15. Zhang, F.; Jiang, S.; Wu, S.; Li, Y.; Mao, C.; Liu, Y.; Yan, H., Complex wireframe DNA origami nanostructures with multi-arm junction vertices. Nature Nanotechnology 2015, 10, 779–784.

16. Rothemund, P. W., Folding DNA to create nanoscale shapes and patterns. Nature 2006, 440, 297–302.

17. Shi, X.; Pumm, A.-K.; Maffeo, C.; Kohler, F.; Feigl, E.; Zhao, W.; Verschueren, D.; Golestanian, R.; Aksimentiev, A.; Dietz, H.; Dekker, C., A DNA turbine powered by a transmembrane potential across a nanopore. Nature Nanotechnology 2024, 19, 338–344.

18. Pumm, A.-K.; Engelen, W.; Kopperger, E.; Isensee, J.; Vogt, M.; Kozina, V.; Kube, M.; Honemann, M. N.; Bertosin, E.; Langecker, M.; Golestanian, R.; Simmel, F. C.; Dietz, H., A DNA origami rotary ratchet motor. Nature 2022, 607, 492–498.

19. Zhan, P.; Yang, J.; Ding, L.; Jing, X.; Hipp, K.; Nussberger, S.; Yan, H.; Liu, N., 3D DNA origami pincers that multitask on giant unilamellar vesicles. Science Advances 2024, 10, eadn8903.

20. Iinuma, R.; Ke, Y.; Jungmann, R.; Schlichthaerle, T.; Woehrstein, J. B.; Yin, P., Polyhedra self-assembled from DNA tripods and characterized with 3D DNA-PAINT. Science 2014, 344, 65–69.

21. Douglas, S. M.; Marblestone, A. H.; Teerapittayanon, S.; Vazquez, A.; Church, G. M.; Shih, W. M., Rapid prototyping of 3D DNA-origami shapes with caDNAno. Nucleic Acids Research 2009, 37, 5001–5006.

22. Doty, D.; Lee, B. L.; Stérin, T., Scadnano: A browser-based, easily scriptable tool for designing DNA nanostructures. In DNA 2020: Proceedings of the 26th International Meeting on DNA Computing and Molecular Programming 2020, Vol. 174, 9:1–9:17.

23. Jun, H.; Wang, X.; Parsons, M. F.; Bricker, W. P.; John, T.; Li, S.; Jackson, S.; Chiu, W.; Bathe, M., Rapid prototyping of arbitrary 2D and 3D wireframe DNA origami. Nucleic Acids Research 2021, 49, 10265–10274.

24. Jun, H.; Zhang, F.; Shepherd, T.; Ratanalert, S.; Qi, X.; Yan, H.; Bathe, M., Autonomously designed free-form 2D DNA origami. Science Advances 2019, 5, eaav0655.

25. Li, R.; Chen, H.; Choi, J. H., Auxetic two-dimensional nanostructures from DNA. Angewandte Chemie International Edition 2021, 60, 7165–7173.

26. Madhvacharyula, A. S.; Li, R.; Swett, A. A.; Du, Y.; Seo, S.; Simmel, F. C.; Choi, J. H., Realizing mechanical frustration at the nanoscale using DNA origami. Nature Communications 2025, 16, 5164.

27. Li, R.; Chen, H.; Lee, H.; Choi, J. H., Conformational control of DNA origami by DNA oligomers, intercalators and UV Light. Methods and Protocols 2021, 4, 38.

28. Lee, C.; Kim, Y.-J.; Kim, K. S.; Lee, J. Y.; Kim, D.-N., Modulating the chemomechanical response of structured DNA assemblies through binding molecules. Nucleic Acids Research 2021, 49, 12591–12599.

29. Rovigatti, L.; Šulc, P.; Reguly, I. Z.; Romano, F., A comparison between parallelization approaches in molecular dynamics simulations on GPUs. Journal of Computational Chemistry 2015, 36, 1–8.

30. Torrie, G. M.; Valleau, J. P., Nonphysical sampling distributions in Monte Carlo freeenergy estimation: Umbrella sampling. Journal of Computational Physics 1977, 23, 187–199.

31. Zhang, D. Y.; Seelig, G., Dynamic DNA nanotechnology using strand-displacement reactions. Nature Chemistry 2011, 3, 103–113.

32. Benson, E.; Marzo, R. C.; Bath, J.; Turberfield, A. J., A DNA molecular printer capable of programmable positioning and patterning in two dimensions. Science Robotics 2022, 7, eabn5459.

33. Sample, M.; Liu, H.; Diep, T.; Matthies, M.; Sulc, P., Hairygami: Analysis of DNA nanostructures’ conformational change driven by functionalizable overhangs. ACS Nano 2024, 18, 30004–30016.

34. Poppleton, E.; Romero, R.; Mallya, A.; Rovigatti, L.; Šulc, P., OxDNA.org: a public webserver for coarse-grained simulations of DNA and RNA nanostructures. Nucleic Acids Research 2021, 49, W491–W498.

35. Zadeh, J. N.; Steenberg, C. D.; Bois, J. S.; Wolfe, B. R.; Pierce, M. B.; Khan, A. R.; Dirks, R. M.; Pierce, N. A., NUPACK: Analysis and design of nucleic acid systems. Journal of Computational Chemistry 2011, 32, 170–173.

36. Li, R.; Chen, H.; Choi, J. H., Topological assembly of a deployable Hoberman flight ring from DNA. Small 2021, 17, 2007069.

37. DeLuca, M.; Duke, D.; Ye, T.; Poirier, M.; Ke, Y.; Castro, C.; Arya, G., Mechanism of DNA origami folding elucidated by mesoscopic simulations. Nature Communications 2024, 15, 3015.

38. Engelhardt, F. A.; Praetorius, F.; Wachauf, C. H.; Brüggenthies, G.; Kohler, F.; Kick, B.; Kadletz, K. L.; Pham, P. N.; Behler, K. L.; Gerling, T.; Dietz, H., Custom-size, functional, and durable DNA origami with design-specific scaffolds. ACS Nano 2019, 13, 5015–5027.

39. He, Y.; Ye, T.; Su, M.; Zhang, C.; Ribbe, A. E.; Jiang, W.; Mao, C., Hierarchical self-assembly of DNA into symmetric supramolecular polyhedra. Nature 2008, 452, 198–201.

40. Evans, E.; Heinrich, V.; Ludwig, F.; Rawicz, W., Dynamic tension spectroscopy and strength of biomembranes. Biophysical Journal 2003, 85, 2342–2350.

41. Akimov, S. A.; Volynsky, P. E.; Galimzyanov, T. R.; Kuzmin, P. I.; Pavlov, K. V.; Batishchev, O. V., Pore formation in lipid membrane II: Energy landscape under external stress. Scientific Reports 2017, 7, 12509.

42. Bennett, W. D.; Sapay, N.; Tieleman, D. P., Atomistic simulations of pore formation and closure in lipid bilayers. Biophysical journal 2014, 106, 210–219.

43. Fan, S.; Wang, S.; Ding, L.; Speck, T.; Yan, H.; Nussberger, S.; Liu, N., Morphology remodelling and membrane channel formation in synthetic cells via reconfigurable DNA nanorafts. Nature Materials 2025, 24, 278–286.

44. Li, Y.; Liu, Z.; Yu, G.; Jiang, W.; Mao, C., Self-assembly of molecule-like nanoparticle clusters directed by DNA nanocages. Journal of the American Chemical Society 2015, 137, 4320–4323.

45. Hindley, J. W.; Elani, Y.; McGilvery, C. M.; Ali, S.; Bevan, C. L.; Law, R. V.; Ces, O., Light-triggered enzymatic reactions in nested vesicle reactors. Nature Communications 2018, 9, 1093.

46. Zhan, P.; Jahnke, K.; Liu, N.; Göpfrich, K., Functional DNA-based cytoskeletons for synthetic cells. Nature Chemistry 2022, 14, 958–963.

47. Hwang, W.; Karplus, M., Structural basis for power stroke vs. Brownian ratchet mechanisms of motor proteins. Proceedings of the National Academy of Sciences 2019, 116, 19777–19785.

48. Wang, S.; Zhao, Y.; Breslawec, A. P.; Liang, T.; Deng, Z.; Kuperman, L. L.; Yu, Q., Strategy to combat biofilms: a focus on biofilm dispersal enzymes. npj Biofilms and Microbiomes 2023, 9, 63.

