## Supplementary Information for "A Nanoscale Jitterbug Transformer from DNA"

**Table of Contents**

S1. Materials and Methods
S2. Design
S3. Computational Design and Analysis
S4. Additional AFM Images
S5. Additional TEM Images
S6. Reconfiguration Induced by External Stimuli
S7. DLS Measurements
S8. Gel Electrophoresis and Structural Yield Estimation
S9. FRET Measurements
S10. Absorbance Spectra of AuNPs
S11. Control Experiments for Nanopore Formation on Synthetic Vesicles
S12. Nanopore Formation and Recovery on Giant Vesicles
S13. Enzyme Transport into Vesicles
S14. DNA Sequences
S15. Supplementary References

### S1. Materials and Methods

#### S1.1 Materials

All DNA oligomers were obtained from Integrated DNA Technologies. The M13mp18 (7k scaffold) was purchased from Bayou Biolabs, while the p9072 (9k scaffold)[1] was provided by Prof. H. Dietz, Dr. M. Honemann, and M. Pinner from the Technical University of Munich. All other chemicals were acquired from Sigma-Aldrich. Filtration components included 100 kDa Pall Nanosep centrifugal filters and Amicon Ultra 50 kDa centrifugal filter units, both supplied by Fisher Scientific. DNA gel extraction was performed using Freeze 'N Squeeze spin columns, and agarose powder was obtained from Bio-Rad Laboratories. The DNA ladder (Quick-Load Purple 2-Log DNA Ladder, 0.1–10 kb) was provided by New England Biolabs. Citrate-stabilized gold nanoparticles (AuNPs with an average diameter of 5 nm), enzymes, and fluorophore-conjugated substrates were obtained from Thermo Fisher Scientific. Lipids including 1,2-dimyristoyl-sn-glycero-3-phosphocholine (DMPC) and 1,2-dioleoyl-sn-glycero-3-phosphoethanolamine-N-biotinyl were purchased from Avanti Polar Lipids. Biotinylated bovine serum albumin (BSA) and streptavidin were obtained from Fisher Scientific and Prospec, respectively. Microscopy substrates consisted of quartz slides from Ted Pella and glass coverslips from Fisher Scientific. Components for microfluidic channels included adhesive sheets from Adhesive Research, CapTite bonded-port connectors, one-piece fittings from Lab Smith, and Tygon microbore tubing from Cole-Parmer.

#### S1.2 Buffer conditions

- TAEM12 buffer (DNA origami assembly): 1×TAE (40 mM Tris-acetate, 1 mM EDTA (ethylenediaminetetraacetic acid) disodium salt, 20 mM acetic acid, pH 8.0) supplemented with 12 mM magnesium acetate.
- TAEM6 buffer (DNA origami purification and agarose gel electrophoresis): 1×TAE supplemented with 6 mM magnesium acetate.
- TAEMC buffer (storage of enzyme molecules): 1×TAE supplemented with 6 mM magnesium acetate and 10 mM calcium acetate.
- TAES buffer (substrate storage): 1×TAE with 2 mM sodium azide.
- PBS buffer (AuNP storage): 137 mM NaCl, 2.7 mM KCl, 10 mM $Na_2HPO_4$, and 1.8 mM $KH_2PO_4$ at pH 7.5.

#### S1.3 DNA origami assembly

DNA nanostructures were assembled by combining 8 nM 7k-scaffold strands and 10 nM 9k-scaffold strands with a 4× excess of staple oligonucleotides for the regular edges and a 6× excess of linker staples to connect the 7k scaffold segment with the 9k scaffold segment. The mixture was subjected to controlled thermal annealing according to the following protocol:

- Initial denaturation at 65 °C for 15 minutes
- Gradual cooling from 60 °C to 40 °C at a rate of - 0.1 °C every 18 minutes
- Final cooling from 40 °C to 4 °C

The complete annealing process required approximately 2.5 days. The initial state shown in Fig. 2c(i) was prepared using this thermal cycling protocol. The octahedral and cuboctahedral structures presented in Fig. 2c(ii)-(iii) were generated by incorporating short jack staples and long jack staples, respectively. Following purification of the initial DNA nanostructures using a 100 kDa centrifugal filter, jack staples were added, and the mixture was incubated at 50 °C for 12 hours. DNA nanostructures designed for UV-triggered reconfiguration were synthesized using the same protocol but incorporated photocleavable jack staples.

#### S1.4 Reconfiguration induced by chemical stimuli

Structural transformation was performed following a three-step protocol: (1) synthesis of the initial configuration (cuboctahedra without jack staples), (2) annealing with a set of jack staples for an intended conformation, and (3) toehold-mediated strand displacement (TMSD) using complementary releaser strands. The initial cuboctahedral structure without jack staples served as the starting point for reconfiguration into either octahedron with short jacks or cuboctahedron with long jacks. After synthesis of the initial conformation, jack staples were added at a 4× molar excess. The mixture was incubated at 50°C for 12 hours, followed by gradual cooling to 4 °C. To reverse Jitterbug transformation, complementary releaser strands were introduced to displace the jack staples through TMSD, and the reaction mixture was subjected to identical thermal conditions (50°C for 12 hours, followed by cooling to 4°C).

Between each step, excess staples were removed by centrifugal filtration. Specifically, 55 µL of DNA solution was diluted with 400 µL TAEM6 buffer and centrifuged at 5,000 rpm for 3 minutes using 100 kDa Pall Nanosep centrifugal filters. This procedure was repeated three times in total, and the purified sample was resuspended in fresh TAEM6 buffer.

UV-responsive structures were prepared by incorporating photocleavable jack staples in step (2). Similar to regular short jack staples, the photocleavable jack staples enabled switching from cuboctahedral to octahedral configurations. Subsequent UV irradiation using a UV lamp (UVGL-25, UVP, Upland, CA) cleaved the jack staples, inducing structural reconfiguration back to the cuboctahedral form. We used long-wavelength UV light (320–400 nm) to minimize potential damage to the stability of the DNA nanostructures.

**S1.5 Instrumentation**

Thermal cycling

All annealing processes and temperature-controlled incubations were performed using a thermal cycler (S1000, Bio-Rad Laboratories).

Fluorescence measurements

Steady-state and time-resolved fluorescence spectra were recorded using a Fluorolog-3 fluorometer (HORIBA JOBIN YVON) equipped with an FL-1039 single-housing unit and a 450 W Xenon lamp.

Giant unilamellar vesicles (GUVs) with fluorophore-conjugated DNA nanostructures were imaged using a custom-built inverted fluorescence microscope (Zeiss Axio Observer D1). Excitation was provided by diode lasers operating at 561 and 658 nm (Laserglow Technologies). Fluorescence emission was collected through a 63× oil-immersion objective (Zeiss) and detected using an Andor iXon3 electron-multiplying charge-coupled device (EMCCD) camera. Images were acquired with full fields of view (135 µm × 135 µm) under both bright-field and total internal reflection fluorescence (TIRF) illumination. To minimize photobleaching, GUV imaging was performed using minimal excitation power with 200 ms exposure times. Image analysis and fluorescence intensity quantification were carried out using ImageJ software.

UV-vis spectroscopy

Absorption spectra were recorded using an Agilent Cary 6000i UV-Vis-NIR spectrophotometer. Sample volumes of ~100 µL were loaded into microcuvettes, and spectra were acquired over the wavelength range from 200 to 800 nm at 20°C.

Dynamic light scattering (DLS)

Hydrodynamic diameters of DNA nanostructures, AuNPs, and DNA-AuNP conjugates were determined using a Zetasizer Nano ZS (Malvern Panalytical, Malvern). All buffers were filtered immediately prior to use. Samples were prepared at concentrations ranging from 0.5 to 2 nM and analyzed in microcuvettes. For each sample, three independent measurements were performed, and the final size distribution was determined by averaging across all measurements.

**S1.6 Imaging**

AFM imaging

AFM imaging was performed in air using PeakForce Tapping mode on a Bruker Dimension Icon equipped with SCANASYST-AIR probes. DNA origami samples were diluted to 0.5–1 nM prior to surface deposition. For sample preparation, 2 µL DNA solution was mixed with 8 µL TAEM6 buffer and deposited on freshly cleaved mica for 5 minutes. To enhance DNA adhesion to the mica surface, 20 µL buffer containing 2.5 mM $NiCl_2$ was added and incubated for an additional 2 minutes. The sample was then gently dried with compressed air and rinsed with 80 µL deionized water. Each mica substrate was scanned at multiple locations using scan sizes of 5 × 5 µm, 2 × 2 µm, 1 × 1 µm, and 500 × 500 nm. AFM images were processed using Bruker Nanoscope analysis software and ImageJ. The yield of DNA structures was estimated from 5 × 5 µm and 2 × 2 µm overview scans by quantifying the ratio of correctly formed structures to misfolded conformations.

TEM imaging

TEM imaging was carried out using a Tecnai G2 T20 microscope operated at 200 kV. For sample preparation, formvar-coated copper grids (Electron Microscopy Sciences) were glow-discharged for 30 seconds using an EasiGlow (Pelco) to render them hydrophilic. A 2 µL aliquot of DNA origami solution was applied to the grid and incubated for 60 seconds before removal with filter paper. The grid was subsequently rinsed by pipetting deionized water until overflow, followed by excess water removal. Finally, 2 µL of 1% uranyl acetate was applied for 20 seconds and removed similarly. Structural yields were calculated as with AFM scans, and edge lengths were measured from the TEM micrographs using ImageJ.

#### S1.7 Agarose gel electrophoresis

Samples were analyzed using 1.0 – 1.5 % agarose gels in TAEM6 buffer for 90 minutes at 75 V. Gels were stained with 0.5 µg/mL ethidium bromide. and visualized using a UV lamp (Spectroline TE-312S, Spectronics). Target bands were excised with a razor blade and transferred to Freeze 'N Squeeze tubes. Gel fragments were mechanically disrupted by repeated compression, frozen at −20°C for 10 minutes, and centrifuged at 13,000× g for 3 minutes to extract DNA. The resulting 16-bit TIFF images were analyzed in ImageJ. Cross-sectional intensity profiles were generated for each lane by averaging grayscale values within defined regions of interest. Peak intensities of monomer bands and higher-order assemblies were quantified to determine sample yields.

#### S1.8 DNA-coated with AuNPs

DNA-AuNP conjugation was performed using a freezing method developed by a previous report[2]. Thiolated DNA oligonucleotides were used without prior reduction treatments such as dithiothreitol (DTT) or Tris-(2-Carboxyethyl) phosphine (TCEP) reduction. In a typical experiment, 5 nm citrate-stabilized AuNPs were mixed with the thiolated DNA strands at a 1:100 molar ratio in PBS buffer. The mixture was placed in a freezer (-20 °C) for 3 hours, then thawed at room temperature. After thawing, excess DNA was removed by three consecutive washing cycles using Amicon Ultra 50 kDa centrifugal filters. The purified DNA-functionalized AuNPs were added to DNA origami structures at 5× molar excess relative to the available binding sites. The mixture was incubated overnight at room temperature to facilitate hybridization between DNA-coated nanoparticles and complementary single-stranded extensions on the DNA nanostructures.

#### S1.9 Enzyme conjugation to AuNPs

Negatively charged citrate-capped AuNPs were used to promote electrostatic adsorption of enzymes, including proteinase K (PK) and trypsin (TRP). This interaction reduced the surface potential through electrostatic shielding and increased hydrodynamic diameter. AuNPs and enzymes were mixed at a 1:100 molar ratio in PBS buffer, sealed in PCR tubes, and incubated at room temperature for 24 hours with constant agitation. After incubation, enzyme-functionalized nanoparticles were purified three using Amicon Ultra 50 kDa centrifugal filters (5,000 × g, 15 minutes). The purified conjugates were resuspended in TAEMC buffer for subsequent hybridization with DNA nanostructures.

For dual-functionalized nanoparticles containing both DNA strands and enzymes, AuNPs were first conjugated with DNA using the freezing method described above. After purification to remove excess DNA, the DNA-coated AuNPs were functionalized with enzymes following the identical incubation and purification protocol.

**S1.10 Microfluidic channel assembly and surface passivation**

Microfluidic channels were constructed by sealing a thin glass coverslip to a quartz slide using medical-grade acrylic double-sided adhesive tape. Channel geometry was defined by slots cut into the adhesive layer. The coverslip was treated with piranha solution ($H_2SO_4$ [3 %]: $H_2O_2$ [1 %]) to generate a hydrophilic surface suitable for biotinylated BSA binding. Inlet and outlet ports were created by drilling the quartz slide and securing CapTite bonded port connectors with epoxy. Fluid flow was controlled using CapTite fittings connected to Tygon microbore tubing and driven by syringe-applied pressure. The total channel volume was approximately 20 μL.

Surface passivation was performed to minimize nonspecific binding of vesicles, DNA nanostructures, and fluorescent molecules. The procedure involved sequential incubations: (1) 100 μL of biotinylated BSA (~0.5 mg/mL) for 10 minutes, (2) 100 μL of 0.2% (v/v) Tween 20 in PBS for 10 minutes to repair surface defects, followed by (3) rinsing with 100 μL PBS buffer (pH 7.5) and (4) final treatment with 100 μL streptavidin solution (10 μg/mL) for 10 minutes to complete biotin-streptavidin surface functionalization. Giant vesicles were introduced and immobilized via biotin-streptavidin interactions between their biotinylated lipids and the surface-bound streptavidin.

**S1.11 Synthesis of giant vesicles**

Pristine giant vesicles were synthesized using the inverted emulsion method[3]. DMPC and biotinylated lipids were combined at a 1,000:1 molar ratio in chloroform within a glass vial. The solvent was removed under vacuum for 30 minutes, and the dried lipid film was resuspended in 600 μL mineral oil under parafilm seal. The oil-lipid mixture was sonicated at 50°C for approximately 3 hours to achieve uniform dispersion.

Aqueous droplets were formed by adding 20 μL deionized water followed by 30 seconds of vortexing, producing a cloudy emulsion with lipid-coated droplets. A total of 600 μL emulsion was carefully layered over 300 μL TAEM buffer to create a hydrophilic interface. The assembly was centrifuged at 8,000× g for 15 minutes, during which monolayer-coated droplets traversed the oil-water interface to form bilayered giant vesicles that settled in the aqueous buffer phase. The mineral oil supernatant was discarded, and vesicles were collected from the buffer phase.

To prepare giant vesicles incorporating Casein-BODIPY (CB) or BSA-TMR conjugates, 20 μL of 10 μM substrate solution replaced deionized water using the identical procedure. Synthesized vesicles were introduced into microfluidic channels and incubated for 45 minutes to establish biotin-streptavidin binding.

### S2. Design

#### S2.1 A Jitterbug transformer

The term 'Jitterbug' was introduced by R. Buckminster Fuller to describe a dynamic geometric transformation of the vector equilibrium stick model[4]. The vector equilibrium is defined as the arrangement of 12 vectors radiating from a central origin to the vertices of a cuboctahedron (Fig. S1a), where each vector forms a 60° angle with its four nearest neighbors, and opposite vectors are collinear. In this configuration, the polyhedron's edge lengths are equivalent to the radial distances from the center to each vertex.

The Jitterbug structure consists of eight identical equilateral triangles connected at shared vertices. Its transformation is a continuous, symmetric motion that switches between a cuboctahedron, an Archimedean solid with eight triangles and six squares (Fig. S1b), and a regular octahedron, a Platonic solid with eight identical triangles[5] (Fig. S1c-e).

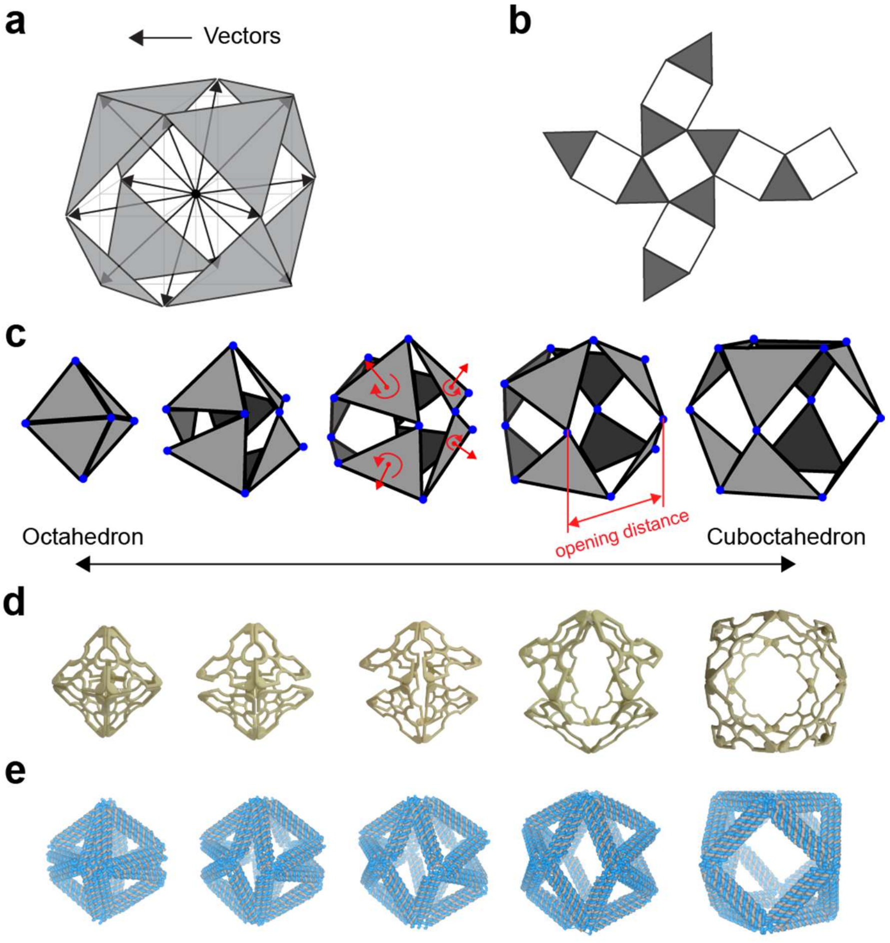


**Figure S1. Diagrams and transformation mechanism of the Jitterbug. a,** Vector equilibrium configuration showing 12 vectors (black arrows) radiating from a central point. Connecting the vector endpoints forms a cuboctahedron, where all edge lengths are equal to the vector lengths. **b,** Planar layout diagram of the Jitterbug transformer geometry. **c,** Transformation pathway from octahedral to cuboctahedral configuration. Red arrows indicate the coupled rotational and radial translational motions of individual triangular faces. **d,** A 3D-printed model demonstrating the Jitterbug transformer mechanism. **e,** An oxDNA model of a DNA Jitterbug transformer. Each edge is made of a 6HB DNA framework. The octahedron-to-cuboctahedron transformation results in approximately fivefold volumetric expansion.

During the transformation, each triangle undergoes a radial displacement combined with a rotation around its 3-fold symmetry axis, resulting in a screw-like motion. Pairs of triangles move symmetrically along and around each of the four 3-fold axes of the octahedron[6]. Since the triangles retain constant size throughout the motion, their vertices trace elliptical paths, as illustrated by the red ellipses in Fig. S2a. Symmetry constraints further confine each vertex to a plane, causing opposite vertices to follow mirrored elliptical segments (Fig. S2b).

The combination of translation and rotation ensures that structural symmetry is preserved throughout the transformation. As each vertex rotates approximately 60° and square faces emerge, the octahedron transforms into a cuboctahedron, expanding the internal volume by a factor of five. To describe the transformation in detail, we present the following formula adopted from previous reports[5, 6]. As shown in Fig. S2e, the trajectory of a vertex follows a segment of an ellipse, which can be described in the YZ plane as:

$$\frac{Y^2}{b^2} + \frac{Z^2}{a^2} = 1 \qquad \text{(Eq. S1)}$$

Here, $a$ is a semi-major axis equal to the edge length $e$ of the octahedron or cuboctahedron, representing the distance from the center to a vertex in the cuboctahedral configuration. Since the ellipse lies on the surface of a circumscribing cylinder, the semi-minor axis $b$ corresponds to the radius, which is the distance from the center of a triangular face to its vertex:

$$a = e\ , b = \frac{a}{\sqrt{3}} \qquad \text{(Eq. S2)}$$

The two focal points of the ellipse are located at (0, $\pm$0.8164·e). To establish the relationship between rotation angle ϕ (along the M-axis) and its radial displacement ($d$) of a triangle from the center (Fig. S2d), the relationship of rotation angle θ (along the X-axis) with the ellipse geometry is derived:

$$3Y^2 + Z^2 = 1 \qquad \text{(Eq. S3)}$$
$$r^2 = Y^2 + Z^2 \qquad \text{(Eq. S4)}$$
$$r \cdot \cos(\theta) = Z \qquad \text{(Eq. S5)}$$

Substituting Eq. S5 into Eq. S4, the coordinates $Y$ and $Z$ can be expressed in terms of θ:

$$Z = \frac{e \cdot cos(\theta)}{\sqrt{3-2co\ ^2(\theta)}}, Y = \frac{e \cdot sin(\theta)}{\sqrt{3-2co\ ^2(\theta)}} \qquad \text{(Eq. S6)}$$

Considering the projection of the trajectory onto the NY plane, which is tilted with respect to the original YZ plane, the ellipse can be expressed as:

$$Y^2 = \frac{1}{3}e^2 - N^2 \qquad \text{(Eq. S7)}$$

Then, the rotation angle ϕ of the triangle around the M-axis can be related to the elliptical trajectory by can be expressed as:

$$\tan(\phi) = Y/N \qquad \text{(Eq. S8)}$$

The relationship between ϕ and the in-plane rotation angle $\theta$ can be derived from Eq. S7:

$$tan^2(\phi) = \frac{\frac{e^2}{3} - N^2}{N^2} = \frac{9-6co\ ^2(\theta)}{3cos^2(\theta)} - 1 \qquad \text{(Eq. S9)}$$
$$\tan(\phi) = \sqrt{3}\tan(\theta) \qquad \text{(Eq. S10)}$$

Using Eq. S10, the vertical displacement of a vertex along the Z-direction can be expressed using ϕ:

$$Z = e \cdot cos(\phi) \quad \text{(Eq. S11)}$$

Accordingly, the radial displacement $d$ of a triangle along the M-axis from the center is:

$$d = \sqrt{\frac{2}{3}} e \cdot cos(\phi) \quad \text{(Eq. S12)}$$

The range of ϕ is $-60° \leq \phi \leq 60°$, where $\phi = 0°$ corresponds to the cuboctahedral configuration and $\phi = \pm 60°$ corresponds to the octahedral configuration[6].

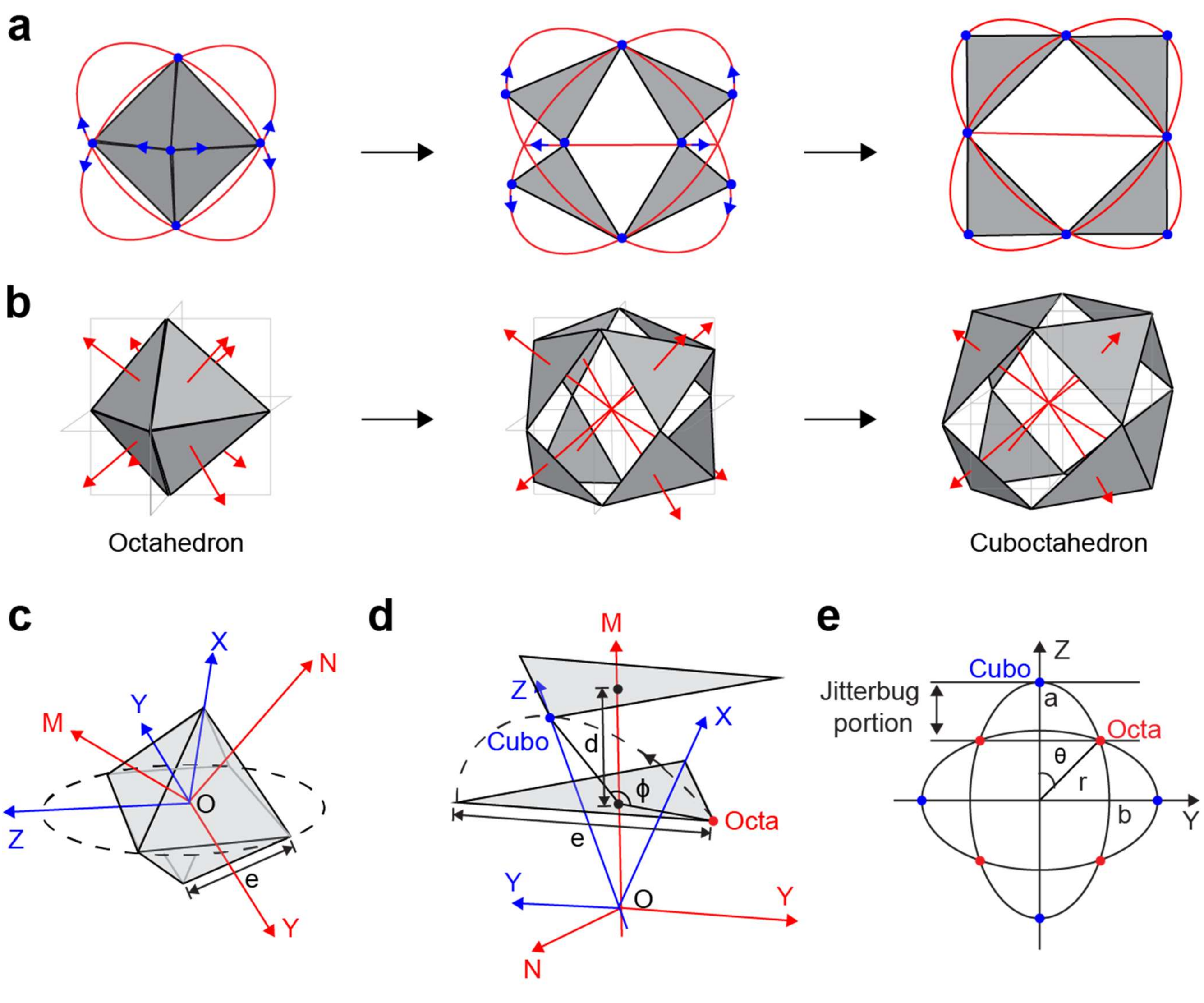


**Figure S2. Kinematic analysis of the Jitterbug transformation. a,** Vertex trajectories during the transformation. Blue arrows indicate the directional motion of individual vertices, while red ellipses represent the elliptical paths traced by each vertex throughout the transformation. **b,** Symmetric deployment of the Jitterbug transformer. Red arrows denote the normal vectors of triangular faces. Each vertex is constrained to planar motion that is mirror-symmetric with respect to its diametrically opposite vertex, ensuring geometric symmetry during the transformation. **c,** Coordinate systems for octahedral geometry. The XYZ and WYV coordinates establish the spatial framework for describing vertex motion. The V and W axes align with triangular face normal vectors, while the X and Z axes represent a unit vector and the normal to the elliptical trajectory plane, respectively. The Y axis serves as the common reference between both coordinates. **d,** Individual vertex trajectory analysis during transformation. As the structure transitions from octahedral to cuboctahedral configuration, each vertex follows a prescribed elliptical path (dashed curve). The triangular face undergoes rotation through angle $\phi$ while its centroid undergoes translational displacement by distance along the V axis. **e,** Vertex configuration as a function of transformation angle. The transformation parameter $\phi$, representing vertex rotation about the X axis, ranges from +60° to −60°. The transformer adopts a cuboctahedral configuration at $\phi$ = 0° and octahedral configurations at the angular extrema $\phi$ = ±60°. The figures were adopted from a previous report[6].

#### S2.2 Design of a Jitterbug transformer using wireframe DNA origami

To design the Jitterbug transformer in a wireframe DNA origami format, the structure was divided into two parts—a front section and a back section—as shown in Fig. S3. Two scaffolds were used: the 7k scaffold (gold in Fig. S3b) for the front and the 9k scaffold (black) for the back. Each edge of the triangles was constructed from 24 six-helix bundle (6HB) edges arranged on a hexagonal lattice to ensure rigidity, resulting in each edge measuring 84 nucleotides in length (approximately 28 nm long). The triangles were numbered from 1 to 8 with their edges sequentially labeled. For example, the three edges of triangle 1 were designated as 1.1, 1.2, and 1.3. As discussed in the main text, actuating the reconfigurable struts (jack edges) enables the Jitterbug transformation between the cuboctahedron and the octahedron by either pinching the six squares with jack staples or releasing them. As shown in Fig. S3b, scaffold strands connected one vertex of each triangle to a vertex of another triangle, creating binding sites for the jack staples.

Fig. S3c shows edge connectivity: vertex A has four connections, while vertex B has only three connections because when two edges meet at vertex B, they form a jack edge that uses up one strand, reducing the available connections. Joints between triangle edges contain 1 single-stranded nucleotide (ssnt), whereas joints between square edges use 10 ssnts, following design suggestions from a previous report[7] on wireframe DNA origami (Fig. S3d). The design was created in Cadnano and Scadnano, with the detailed Scadnano layout shown in Fig. S5.

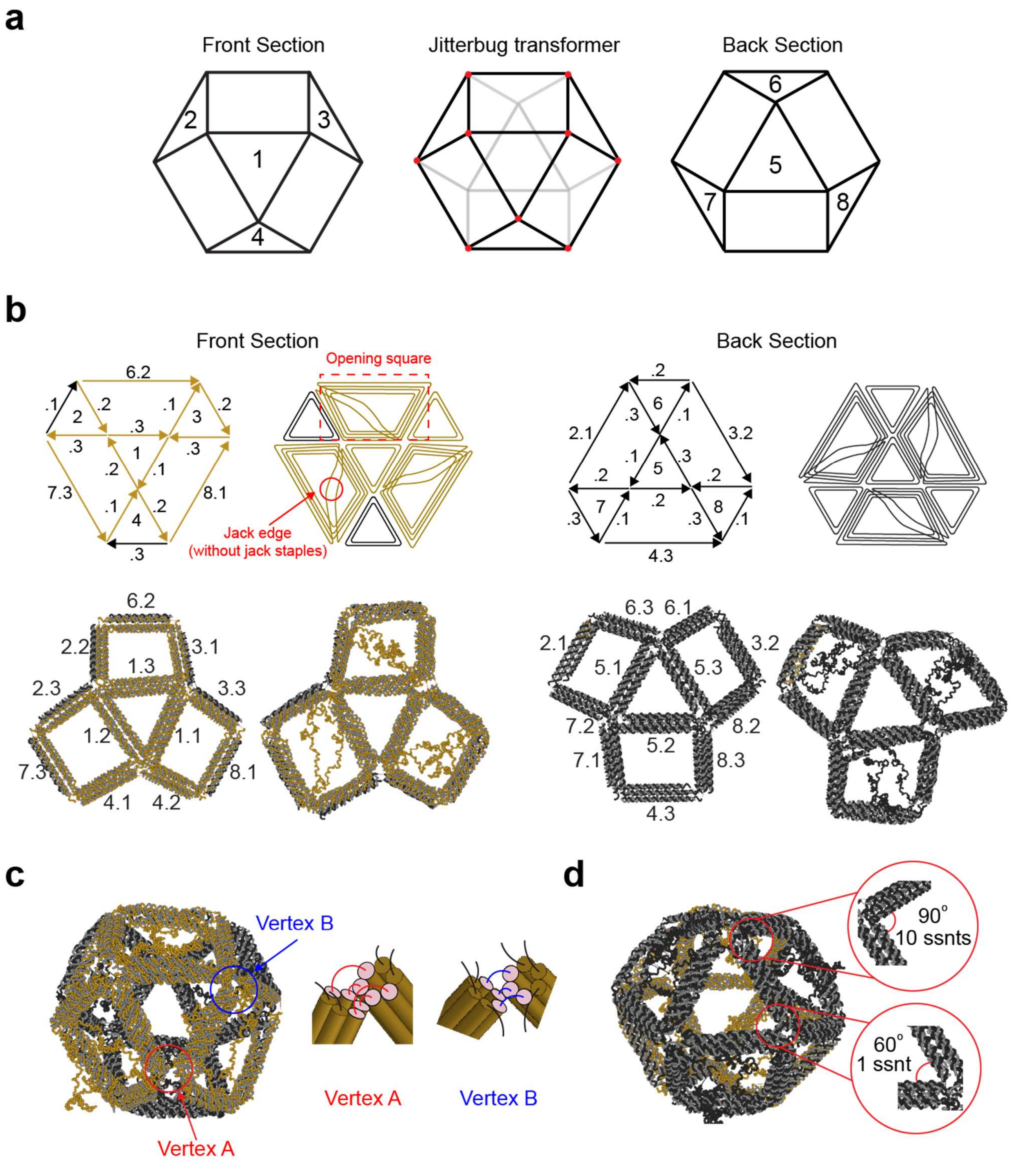


**Figure S3. Wireframe DNA origami design of the Jitterbug transformer. a,** Structural organization showing front and back sections of the Jitterbug transformer with triangular faces labeled 1 through 8. **b,** Design schematics of the wireframe DNA origami architecture. Gold strands represent the 7k scaffold, while black strands denote the 9k scaffold. Arrows indicate the 3' ends of the DNA strands. **c,** Edge connections: vertex A connections link two edges with four DNA strands, while vertex B connections use three strands. **d,** Joint design: square vertices incorporate 10 ssnts (0°–90°), while triangular vertices contain 1 ssnt (always at 60°).

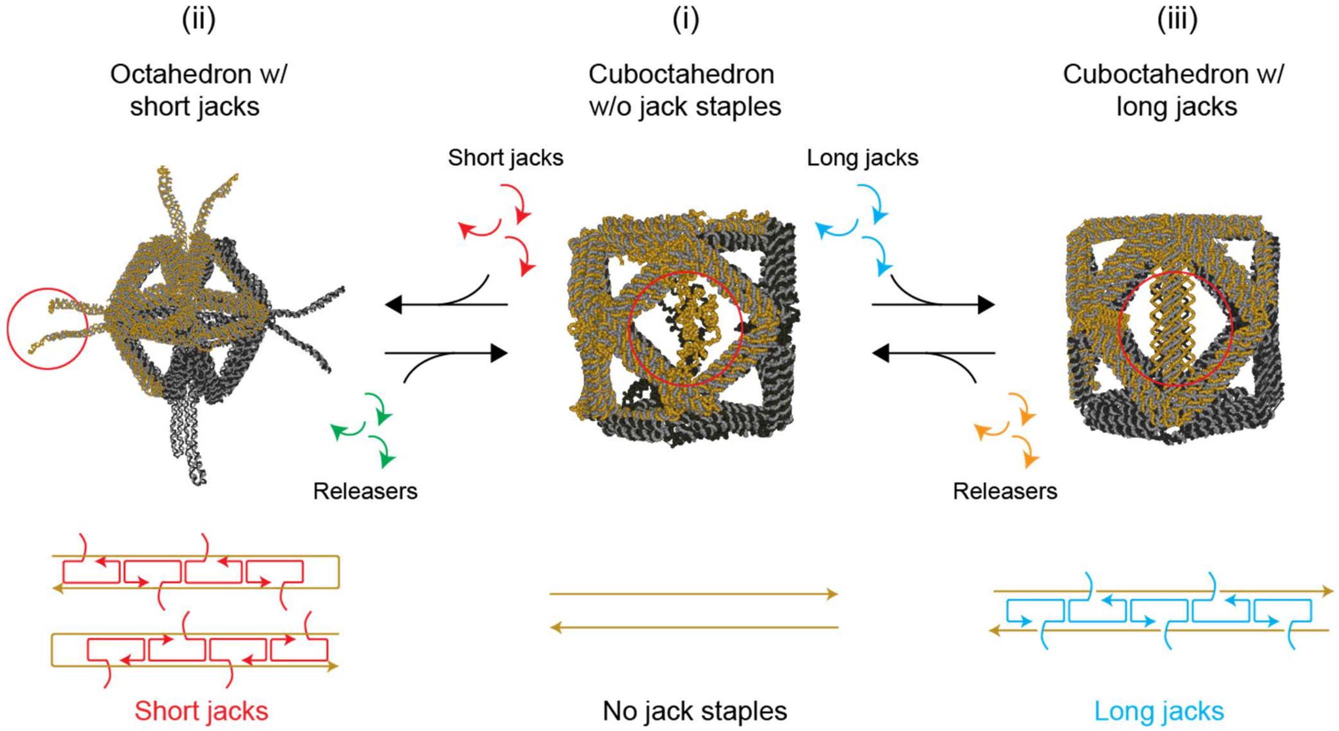


**Figure S4. Jack staple arrangements and toehold-mediated strand displacement mechanisms.** (i) Initial cuboctahedral configuration without jack staples, representing the baseline structural state prior to jack staple incorporation. Subsequent reconfiguration is determined by a specific set of jack staples introduced. (ii) Octahedral conformation achieved by inserting short jack staples (red), which induce contraction of the jack edges. Four short jack staples (red) stabilize the octahedral geometry, and the structure can be reverted to the initial state through addition of complementary releaser strands (green) that target the toehold overhangs of the short jack staples. (iii) Cuboctahedral conformation by incorporation of long jack staples (blue), which maintain extended jack edges. Reconfiguration to the initial state is achieved by introducing corresponding releaser strands (orange) that displace the long jack staples via TMSD.

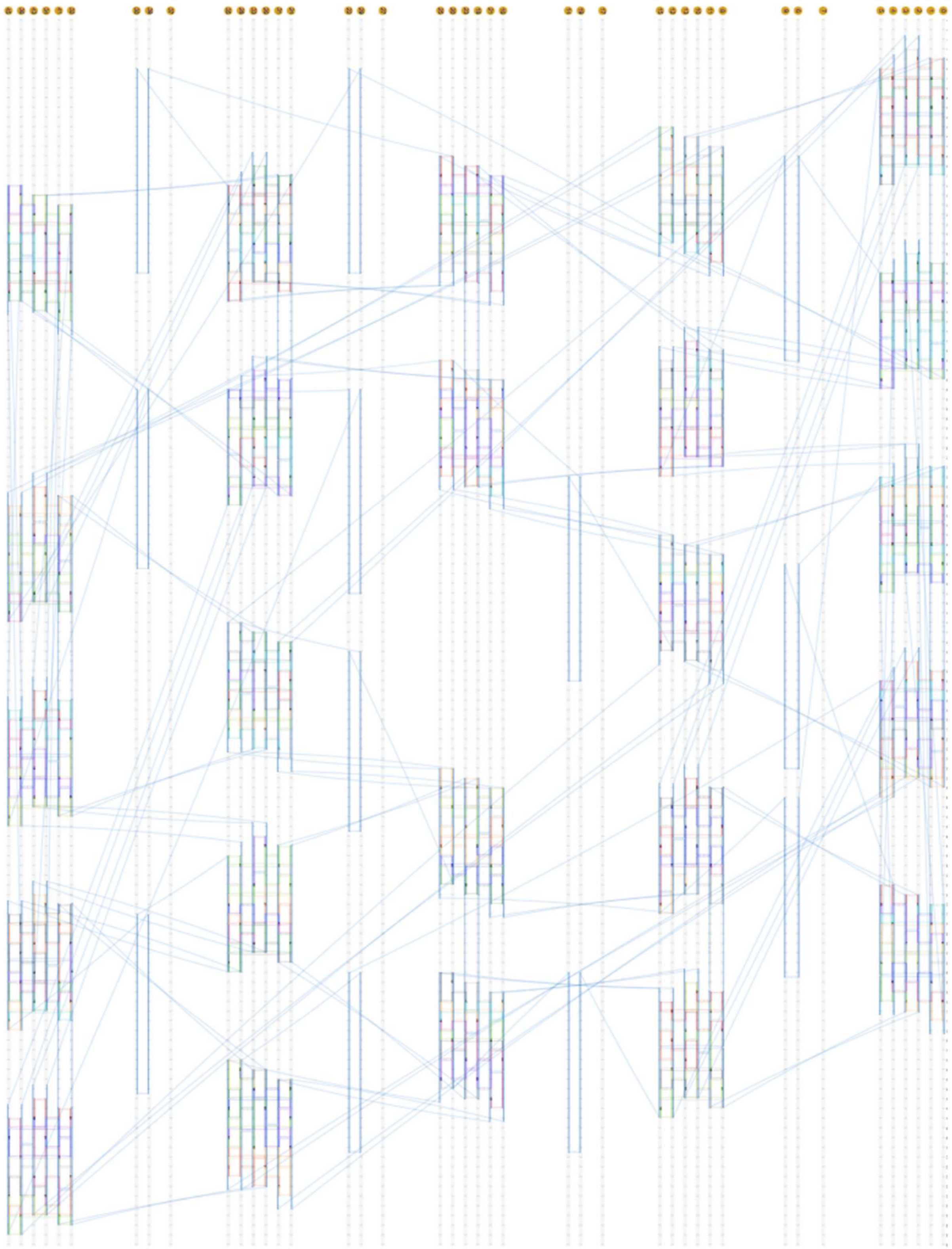

**Figure S5.** Wireframe DNA origami design of the Jitterbug transformer created in Scadnano.

### S3. Computational Design and Analysis

#### S3.1 Minimizing sequence interference to improve structural yield

Due to their large size and incorporation of 6HB edges, the designed origami structures required two scaffolds: a 7249-nt M13mp18 scaffold (7k scaffold) and a custom-designed 9072-nt strand (9k scaffold). Since both scaffolds share similar sequences with considerable overlap, experimental yields varied significantly depending on the specific staple sequences used. Sequence overlaps among staples led to increased interference, resulting in a higher occurrence of misformed or partially assembled structures[8].

This problem was addressed by reducing staple interference. Staple interference was defined as the maximum sequence overlap between any two staples within a given set. To minimize interference, multiple staple permutations were generated by shifting the starting index along the circular 7k scaffold sequence while keeping the staple sequences for the 9k scaffold portion constant. Permutations exhibiting the lowest sequence overlap (approximately 10 nucleotides) were selected and further filtered to eliminate staples prone to unintended hybridization with other scaffold regions or unpaired single-stranded portions. This process significantly improved the yield of Jitterbug DNA structures. The overall workflow for this staple interference minimization is illustrated in Fig. S6.

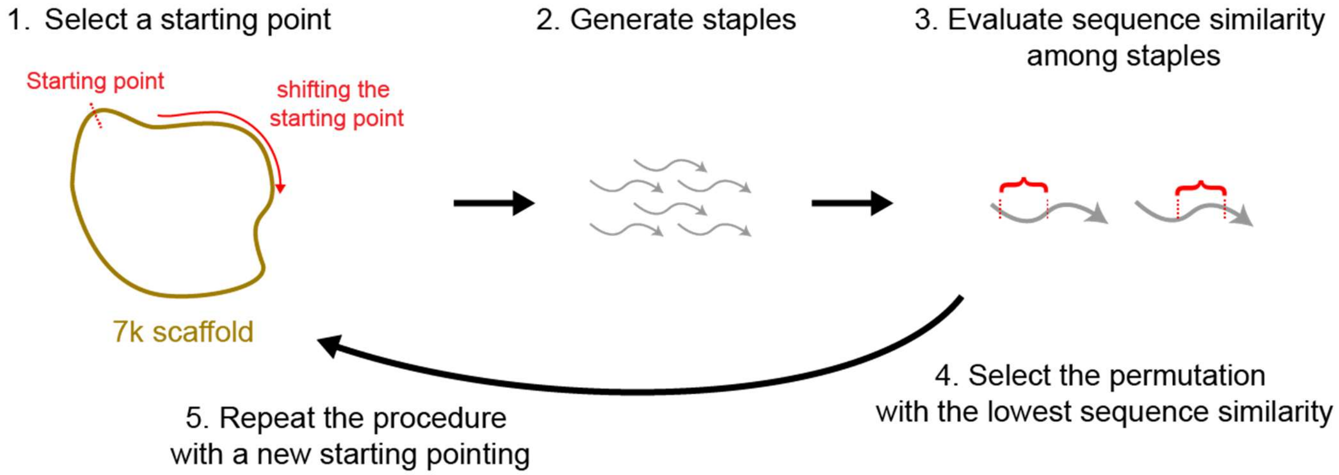


**Figure S6.** Workflow diagram of the iterative process to identify staple sets with minimal inter-staple interference.

**S3.2 Reconfiguration analysis using oxDNA simulation**

To investigate equilibrium properties and dynamic motion during reconfiguration, coarse-grained MD simulations were performed using oxDNA software. For computational analysis, the DNA origami design was first exported from Scadnano into oxDNA, preserving the experimental DNA sequences. Edges and jack segments were identified using density-based spatial clustering of applications with noise (DBSCAN), resulting in 24 edges and 12 jack segments (two segments per square face). For each edge, the positions of the terminal hybridized nucleotides (12 nucleotides at each end) were noted. The opening distance—defined as the distance between the centers of mass of these terminal nucleotides at jack vertices—served as an order parameter in subsequent free energy simulations.

The reconfiguration was analyzed by monitoring the evolution of the average opening distance across all six squares upon removal of jack strands from the octahedral state (Fig. 1g). Individual opening distances are detailed in Fig. S7a. The average opening distances were approximately 13 nm for the octahedral and ~37 nm for the cuboctahedral states. The structure completed the spontaneous transition within approximately $5\times10^6$ simulation steps. During the transformation, radial displacements of constituent triangles and their rotation angles about the radial axis (defined in SI Section 2) were tracked (Fig. 1f). However, the origami structure could not achieve the entire theoretical range of motion, due to the finite thickness of edges, contrasting with the ideal Jitterbug model that assumes edges of zero thickness. Upon staple removal, deformation occurred nearly symmetrically, as expected. The symmetry was confirmed by measuring the angles between the fitted planes of three pairs of opening squares (shown in Fig. S3b), which remained at approximately 90° throughout the reconfiguration process (Fig. S7b).

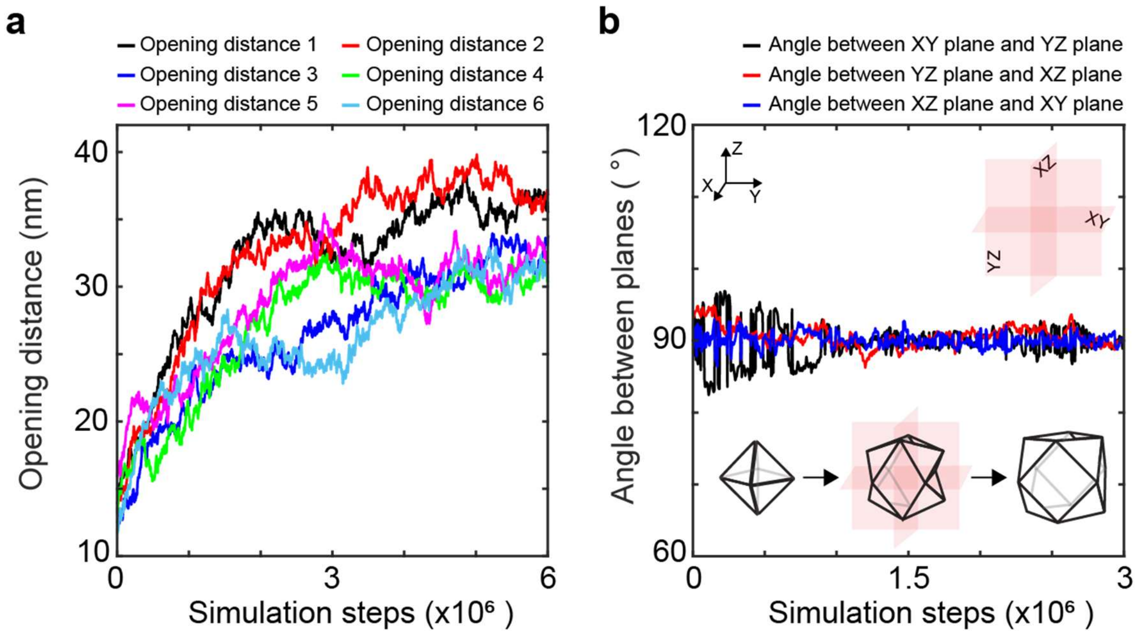


**Figure S7. Kinetic analysis of Jitterbug transformation upon jack staple removal. a,** Evolution of opening distances for all six square faces following jack staple displacement. **b,** Validation of symmetric transformation dynamics. Angular relationships between the three orthogonal normal planes of the Jitterbug structure were calculated throughout the simulation trajectory. The angles were maintained consistently at ~90°, indicating that the engineered DNA nanostructure exhibits conformational dynamics with strong agreement to the theoretical Jitterbug transformation model.

#### S3.3 Poisson's ratio during structural transformation

The three-fold symmetry of the Jitterbug transformer demonstrates a theoretical Poisson's ratio of $\nu = -1$ in all three directions. According to the elasticity theory, Poisson's ratio is defined as:

$$\nu = -\frac{\varepsilon_{trans}}{\varepsilon_{axial}}$$

where $\varepsilon_{axial}$ and $\varepsilon_{trans}$ are axial and transverse strains, respectively. For the Jitterbug transformer, having three orthogonal axes, the effective Poisson's ratio is calculated as the mean of the ratios among strains in these directions:

$$\nu_{jitterbug} = \langle\frac{\nu_{1,2} + \nu_{2,3} + \nu_{3,1}}{3}\rangle$$

where directions 1, 2, and 3 correspond to the three normal axes of the six squares in cuboctahedral configuration, with the origin located at the center of the structure. The effective Poisson's ratio presented in Fig. 1e is obtained by averaging results from ten parallel simulations.

#### S3.4 Free energy simulations

Free energy landscape provides insights into the strain energy stored within the DNA nanostructure, elucidating its spontaneous transition from the octahedral to the cuboctahedral states upon jack removal following our design strategy. To better understand the free energy, we decomposed it into two components: (i) structural free energy inherent to the cuboctahedron and (ii) hybridization energy gained from jack staples, which hold the structure from open to closed states (chemical stabilization).

We estimated the structural free energy of the Jitterbug (without jack staples) using umbrella sampling simulations[9, 10] in oxDNA on NVIDIA 4070 GPU. We assumed symmetric deformations for computation. The average opening distance across all six squares served as the order parameter. MD simulations employed harmonic potential simultaneously applied to all opening squares, each with identical stiffness (11.4 pN/nm; 0.2 simulation units in oxDNA). For each square, 24 nts on each side of the jack are chosen (12 nt per 6HB edge) for applying the harmonic biasing force.

A total of 100 simulation windows were generated, spanning opening distances from 3.6 to 64 nm (4–75 simulation units). Each window was equilibrated over $3\times10^6$ simulation steps, followed by a production run of $9\times10^6$ steps to derive the free energy profile. The resulting distributions of opening distances were averaged across squares and unbiased using the weighted histogram analysis method (WHAM). Figure S8 presents the energy profiles for individual jack segments, while Fig. 1d presents their average profile. The total energy is obtained by summing the contributions from all six squares. The free energy profile shown in Fig. 1d was obtained by unbiasing the average order parameter across all the squares.

$$\Delta G_{total,struc} \approx 6 \cdot \Delta G_{avg,square}$$

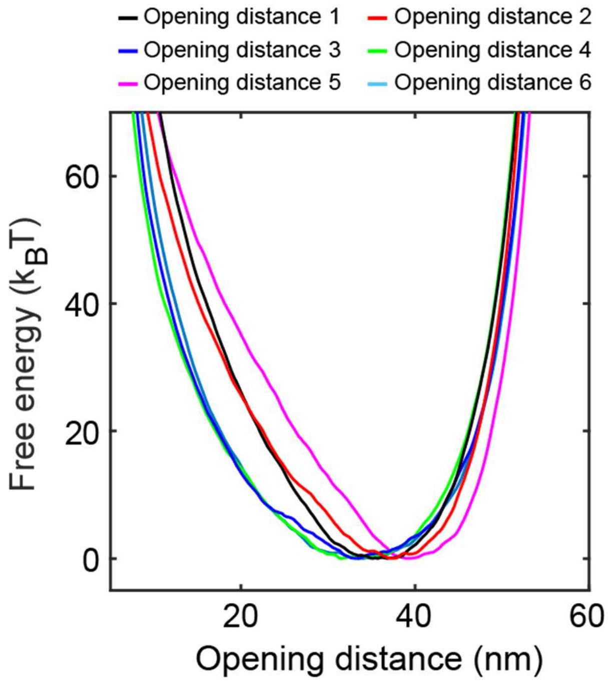


**Figure S8.** Free energy profiles of six opening squares using WHAM.

**S3.5 Stabilization energy for octahedral configuration**

The amount of energy needed to stabilize octahedral configuration is provided via base-pairing. From the free energy profile of cuboctahedral configuration (Fig. 1d), NUPACK calculations suggest that at least 20 base-pairs (~40 $k_BT$) are required for a stable octahedral state.

To further analyze this process, we investigated the dissociation of the first 10 nucleotides in each jack segment. In the octahedral state, the jack segments experience non-zero force due to strain energy stored in the structure, which was estimated to be ~6 pN per jack segment within the 12–20 nm deformation range. To model this, a constant planar force with stiffness 0.057 pN/nm ($10^{-3}$ simulation units) was applied using repulsion planes. Virtual-move Monte Carlo (VMMC) simulations using oxDNA were then performed on a single jack duplex, tracking the dissociation of the first 10 bp (5 bp on each helix shown in Fig. S9). The order parameter was defined as the number of dissociated base-pairs, ranging from 0 to 10 bp. Ideal weights were identified through iterative updates over 10 iterations.

Simulations were conducted in two overlapping windows (0–6 bp and 4–10 bp) for 1 billion steps per window. Convergence was achieved when free energy profiles stabilized, showing negligible changes across different data portions. The simulations also printed the opening distance of the jack segment (distance between the first set of scaffold nucleotides, indicated by arrows in Fig. S9a). Free energy profiles were mapped as a function of distance using the same weights from the original simulations, with samples placed into 1.2 nm wide windows. The resulting free energy was doubled to account for contributions from both jack segments (Fig. S9). As illustrated in the octahedral part of the free-energy plot in Fig. 1b, the total free energy of the system was thus expressed as:

$$\Delta G_{total} = \Delta G_{struc,cubo} + \Delta G_{hyb,jack\ segments}$$

The VMMC profile was integrated with the structural component by aligning them along the x-coordinate. Since base-pairing provides stabilization energy, this energy term carries a negative sign. Given that the fully hybridized octahedron represents an energy minimum, we aligned the minimum from the VMMC simulations with the opening distance of the octahedron obtained from MD simulations, which was approximately 13 nm.

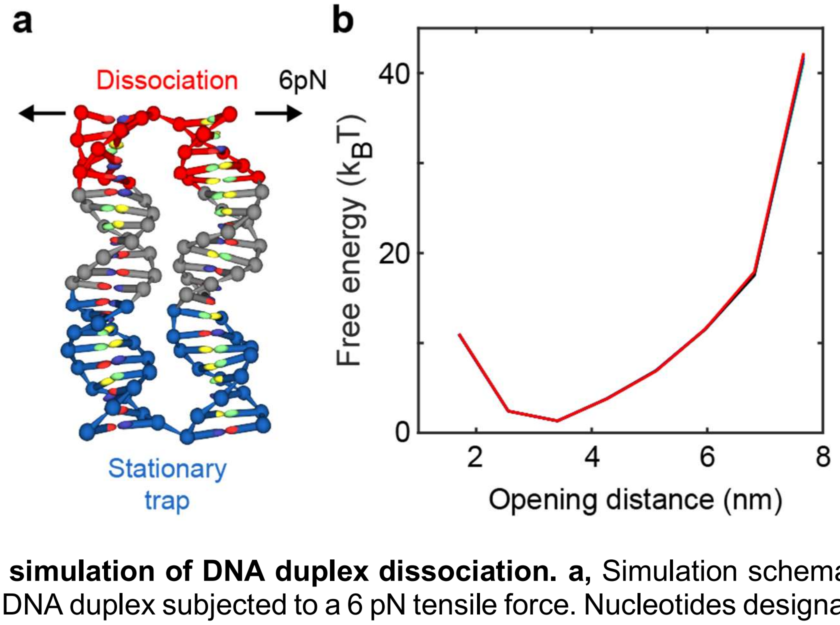


**Figure S9. VMMC simulation of DNA duplex dissociation. a,** Simulation schematic showing a two-helix bundle (2HB) DNA duplex subjected to a 6 pN tensile force. Nucleotides designated for dissociation analysis are highlighted in red, while blue nucleotides are constrained using harmonic potential wells to maintain positional stability. **b,** Free energy profile as a function of opening distance for the staple duplex across multiple independent simulation segments. The negligible variation between resulting curves demonstrates simulation convergence and statistical reliability.

**S3.6 Elastic energy stored in Jitterbug**

To emulate the behavior of an 'ideal' Jitterbug transformation, we estimated elastic energy stored within the structure. Here, the triangular panels were designed to be perfectly rigid, and each opening square was constructed with two alternating vertex types as illustrated in Fig. 1h. To analyze the distribution of elastic energy, the geometry was decomposed into three components: (i) the rigid triangular edges, (ii) vertex A, featuring four 10-nt single-stranded (ssDNA) connections that mediate reconfiguration, and (iii) vertex B, with three 10-nt ssDNA connections that primarily rotate without significant deformation.

For a detailed understanding, the Jitterbug structure was reduced to its simplest building block: a pair of triangles connected by vertices A and B. As described in SI Sections 2 and 3.2, vertices are constrained to normal planes. This confinement was mimicked using repulsion planes with a stiffness of 0.005 simulation units, restricting vertices along two orthogonal planes. Umbrella sampling combined with WHAM was then used to calculate the free energy of this building block as a function of the opening distance.

Harmonic biasing potentials (stiffness: 11.4 pN/nm, 0.2 simulation units in oxDNA) were applied over 100 windows, spanning opening distances from 3.6 nm to 64 nm (4–75 simulation units). Each window was equilibrated for 2 million steps and then simulated for additional 7 million steps. To compare this energy to that of a full opening square (Fig. 1j), the result was multiplied by a factor of two because one opening square consists of two triangular pairs. The resulting free energy profile for this triangular pair (denoted vertex AB) is shown in Fig. S10.

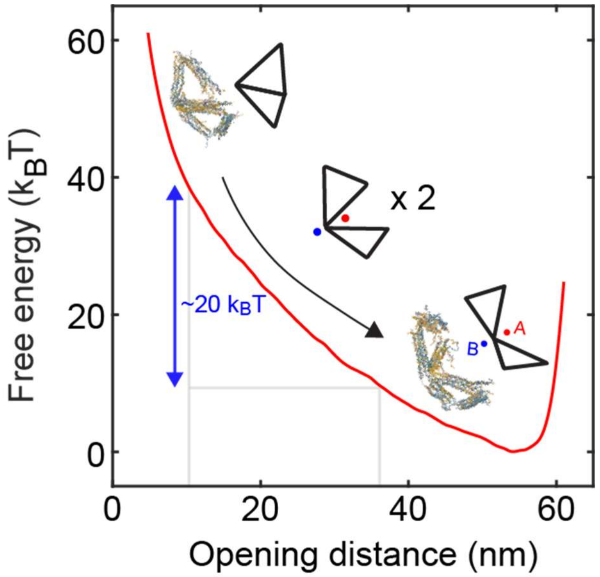


**Figure S10. Free energy profile for opening dynamics of a triangular pair.** The energy profile represents the transformation of a triangular pair, equivalent to half of a square face opening event. The calculated free energy change is ~20 $k_BT$. This result is multiplied by two for a full square in Fig. 1j.

#### S3.7 Comparison of energy estimations

We compared estimated energies involved in the Jitterbug transformation from several methods: (i) umbrella sampling, (ii) NUPACK (for hybridization energy), (iii) VMMC simulations, and (iv) mechanical work.

- Umbrella sampling**:** We unbiased the umbrella sampling simulations using WHAM and measured the free energy difference between the closed (13 nm) and open (37 nm) states:

$$\Delta G_{structural} = G_{structural,13nm} - G_{structural,37nm} \approx 40.5 k_B T$$

- NUPACK: We used NUPACK to estimate the number of base pairs required to provide a minimum energy for stabilization of the octahedral state. Approximately 10 bp per jack segment (20 bp per opening square) were sufficient:

$$\Delta G_{NUPACK,,per\ bp} \approx 1.3\ kcal/mol$$
$$\Delta G_{NUPACK,2\times 10bps} \approx 24\ kcal/mol \approx 44 k_B T$$

  This estimate accounts only for hybridization energy; more staples were used experimentally to offset the entropic cost of bringing scaffold ends together.

- VMMC: The free energy difference between the fully hybridized and 10 bp dissociated states was calculated:

$$\Delta G_{vmmc,2\times 10bps} \approx 40.5 k_B T$$

- Mechanical work**:** The elastic energy was calculated by averaging the force required to maintain the opening distance during the square opening process (Fig. 1e) and integrating over the deformation range (13–37 nm) under the assumption of quasi-equilibrium (Fig. 1k):

$$\Delta G_{mechanical} = \int_{13nm}^{37nm} F\, dx = \sum_{13nm}^{37nm} F\, \delta x \approx 42.4 k_B T$$

Overall, these parallel approaches converge on energy per square within a few $k_B$T.

#### S3.8 Analysis of the jack staple binding process using DNAfold simulations

To reconcile the difference between the theoretical hybridization energy estimate and the number of staples required experimentally, we performed coarse-grained simulations of a simplified jack design using DNAfold[11]. Figure S11 presents the layout of the strands for simulations. We varied the number of staples from 1 to 4 and measured the relative binding times of different segments across 10 simulations for each case. Since DNAfold software supports only square lattices and coarse-grains the system into 8-nt segments, the design was adapted to use 48-nt staples; this adjustment does not significantly affect the observations. The relative binding time for each segment was defined as:

$$t_{rel} = \frac{t_{segment}}{t_{mean}}$$

where $t_{rel}$ is relative binding time, $t_{segment}$ is binding time of the specific segment, and $t_{mean}$ is mean binding time across all segments. The simulations (Fig. 1i) revealed that staples closest to the looped end of the scaffold (i.e., the rightmost staples) bind first, while the leftmost strands hybridize last. This means that the initial binding reduces the entropic cost for bringing the separated scaffold ends closer together, thereby facilitating the binding of subsequent staples.

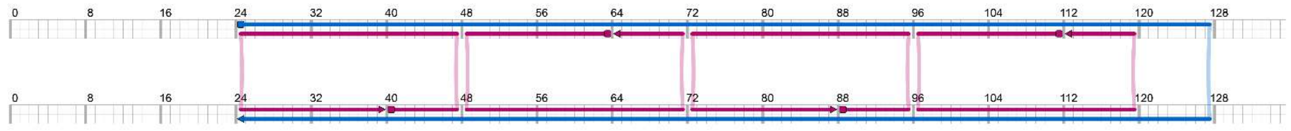


**Figure S11.** Scadnano design of the simplified jack structure for DNAfold simulations.

### S4. Additional AFM Images

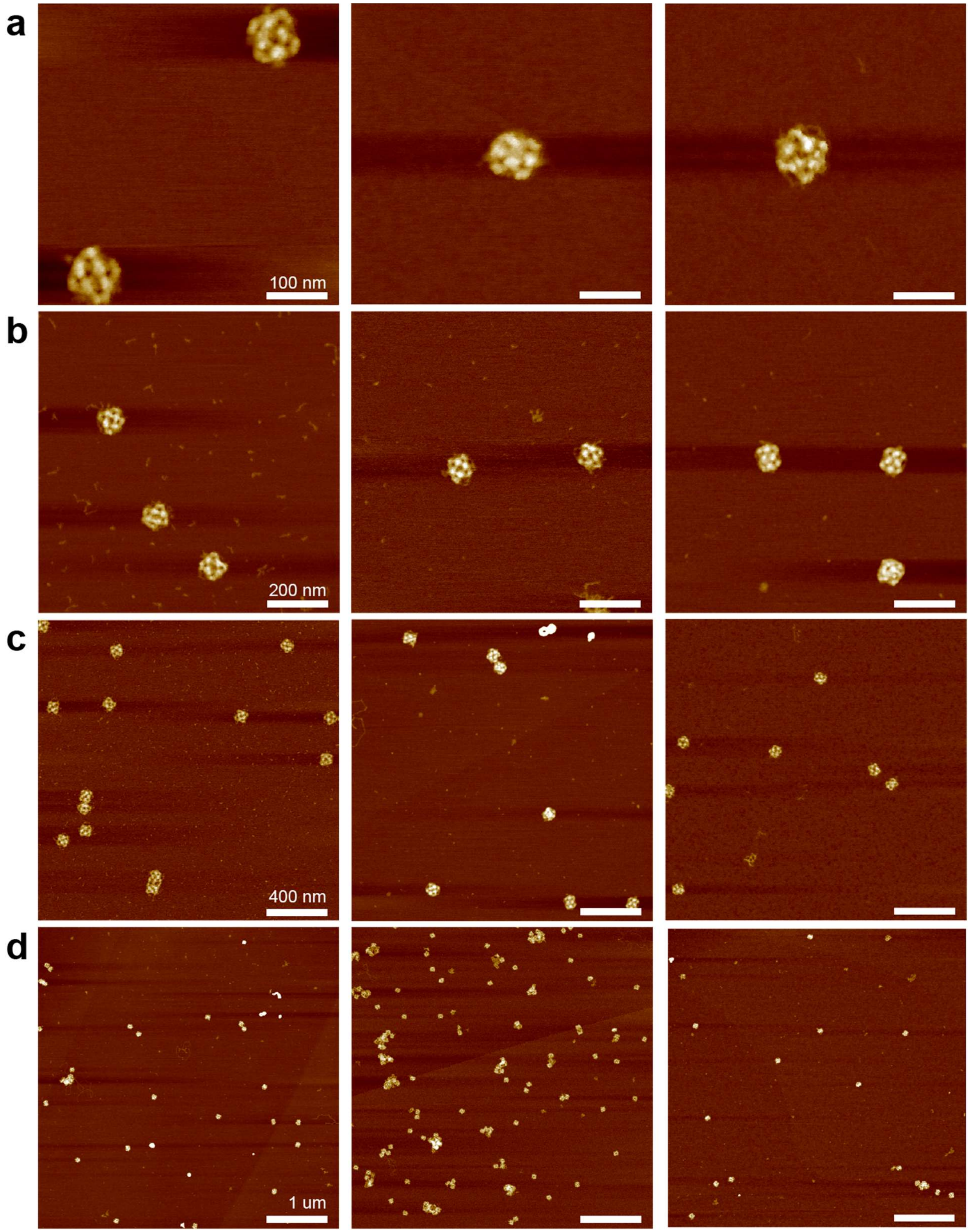


**Figure S12. AFM images of initial cuboctahedra without jack staples.** Images correspond to Fig. 2c(i) at different magnifications. Scale bars: **a** 100 nm, **b** 200 nm, **c** 400 nm, **d** 1 μm.

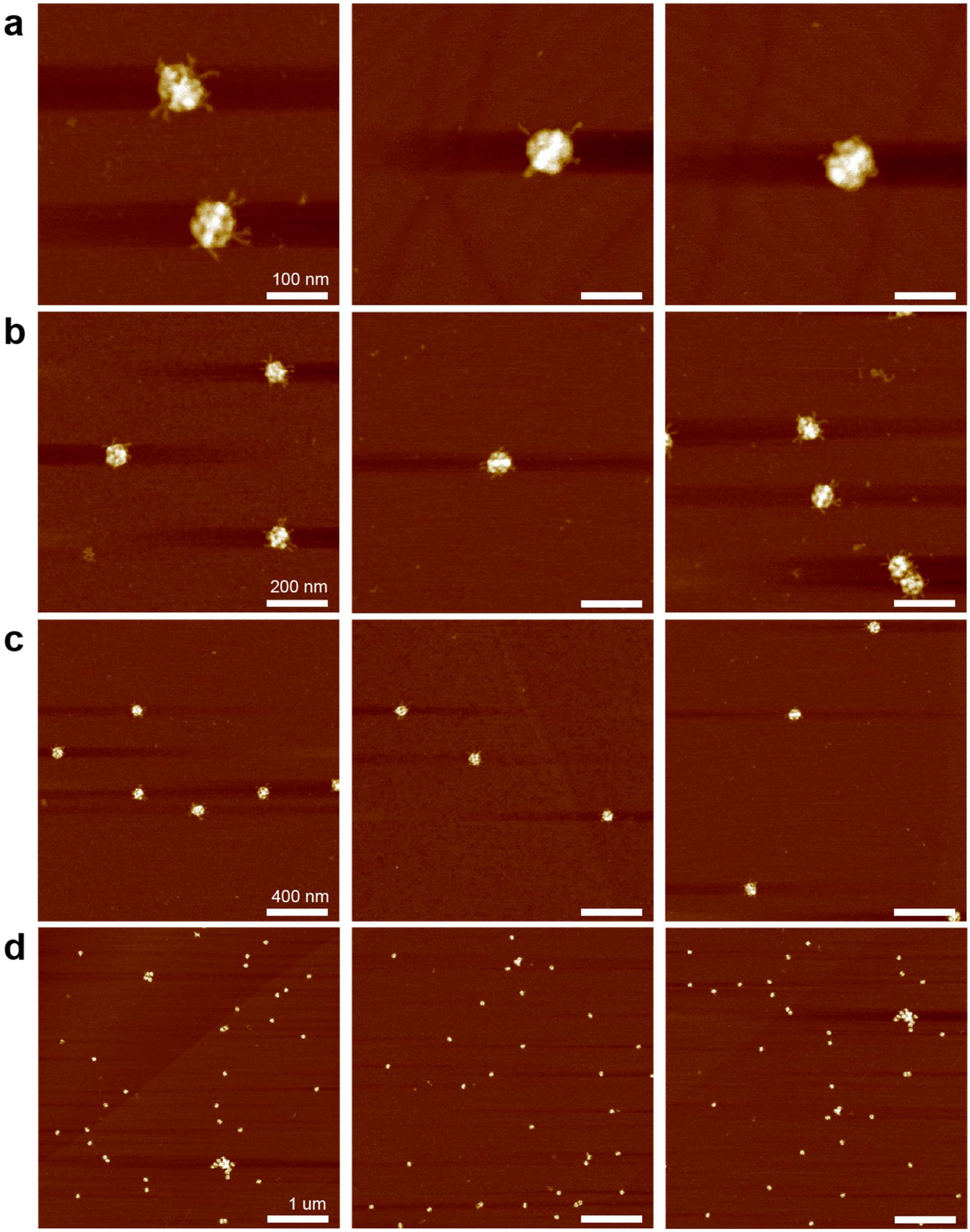


**Fig. S13. AFM images of octahedra with short jack staples.** These AFM scans correspond to those in Fig. 2c(ii). Scale bars: **a** 100 nm, **b** 200 nm, **c** 400 nm, **d** 1 µm.

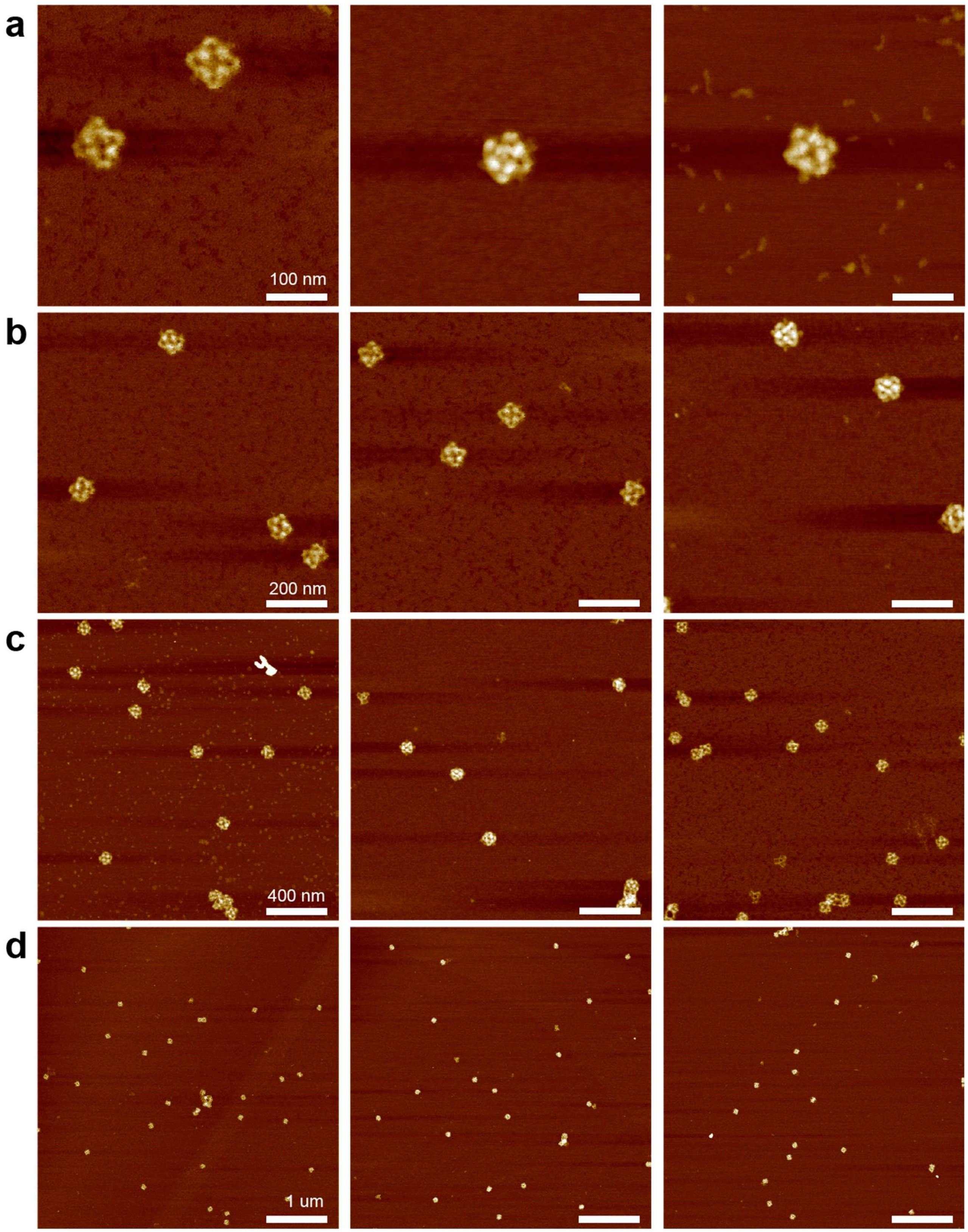


**Figure S14. AFM images of cuboctahedra with long jack staples.** These images correspond to Fig. 2c(iii). Scale bars: **a** 100 nm, **b** 200 nm, **c** 400 nm, **d** 1 μm.

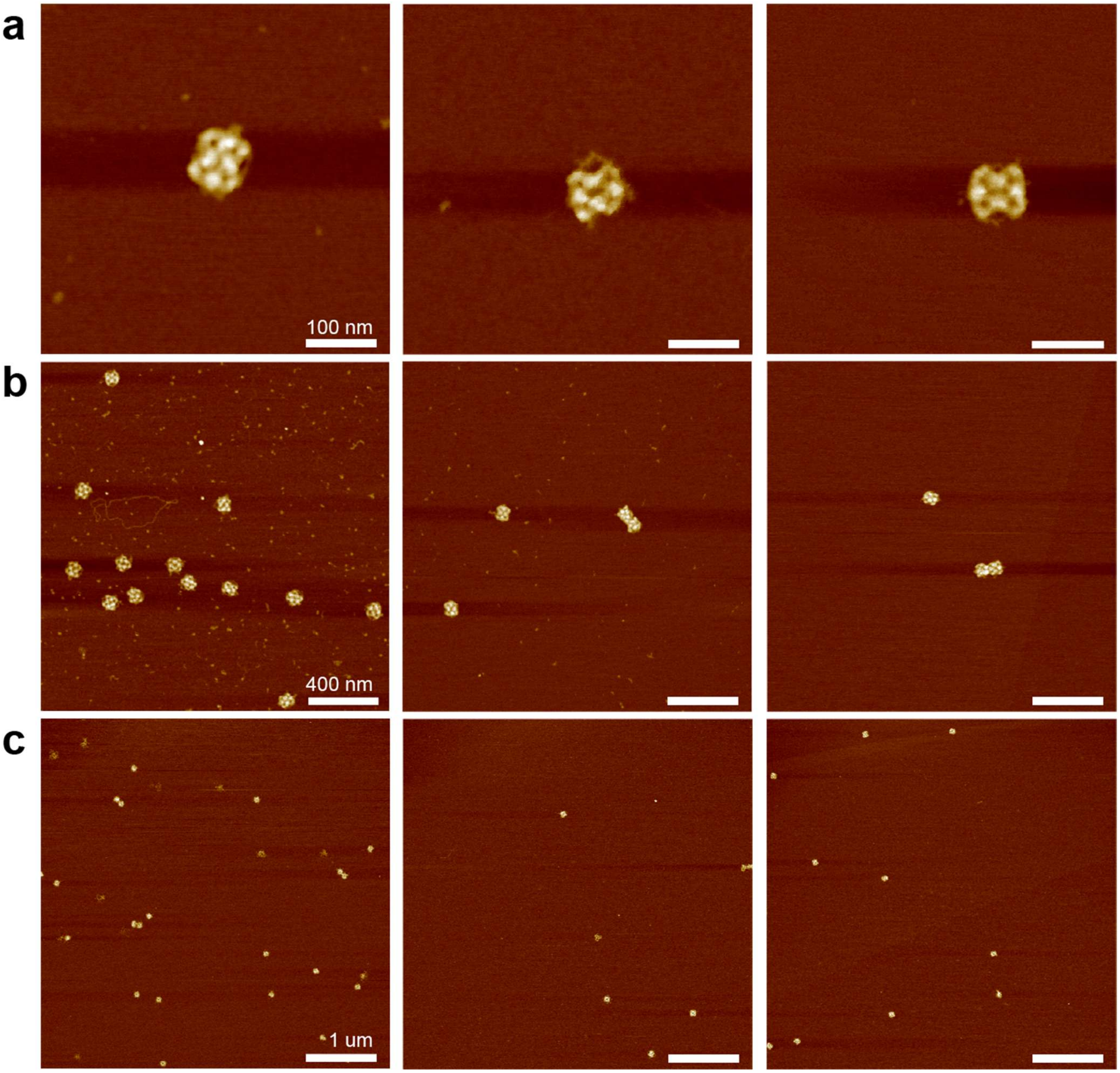


**Figure S15. AFM images of reconfigured cuboctahedra without jack staples.** Scale bars: **a** 100 nm, **b** 400 nm, **c** 1 µm.

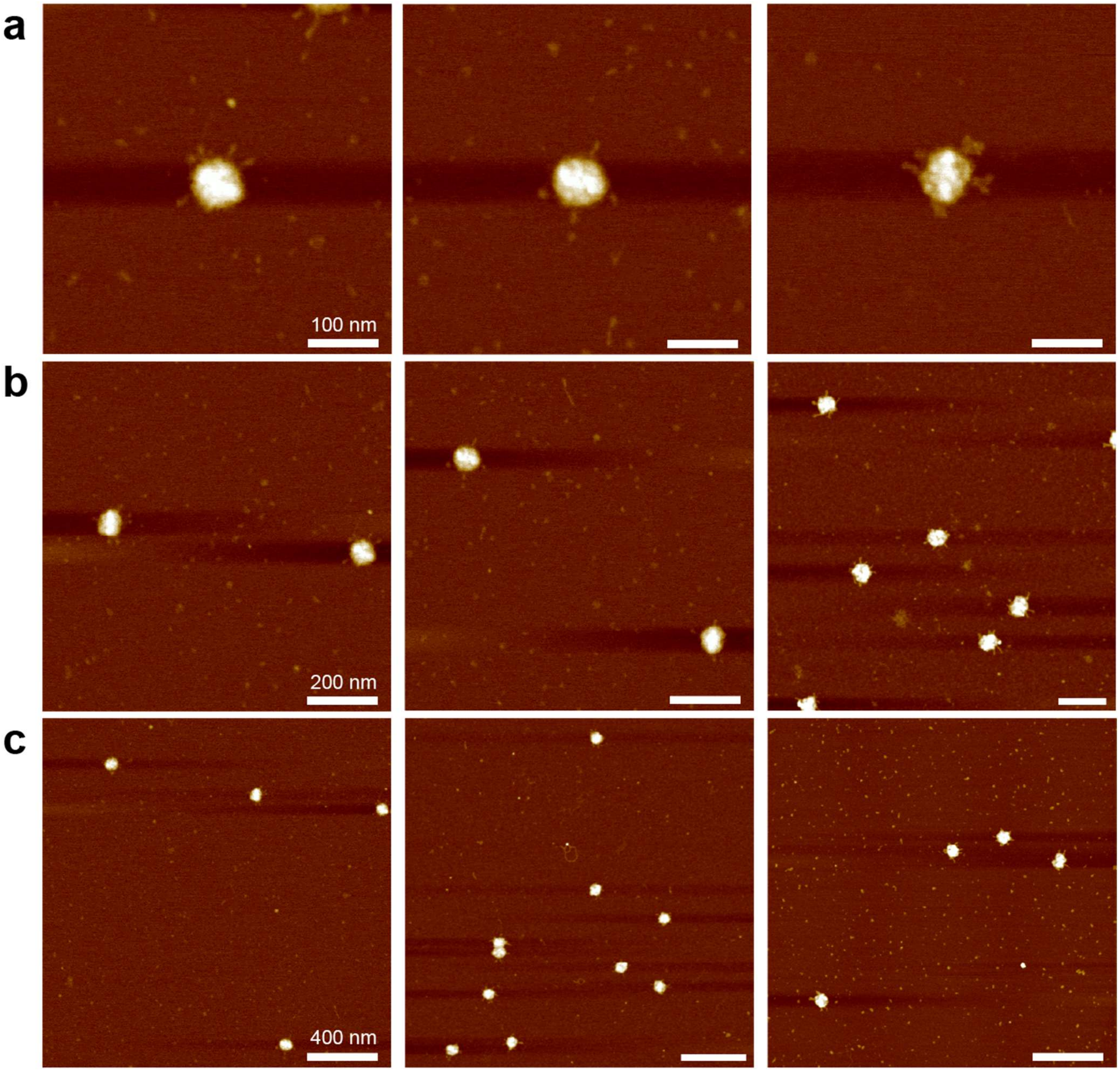


**Figure S16. AFM images of octahedra reconfigured from cuboctahedra with long jacks.** Scale bars: **a** 100 nm, **b** 200 nm, **c** 400 nm.

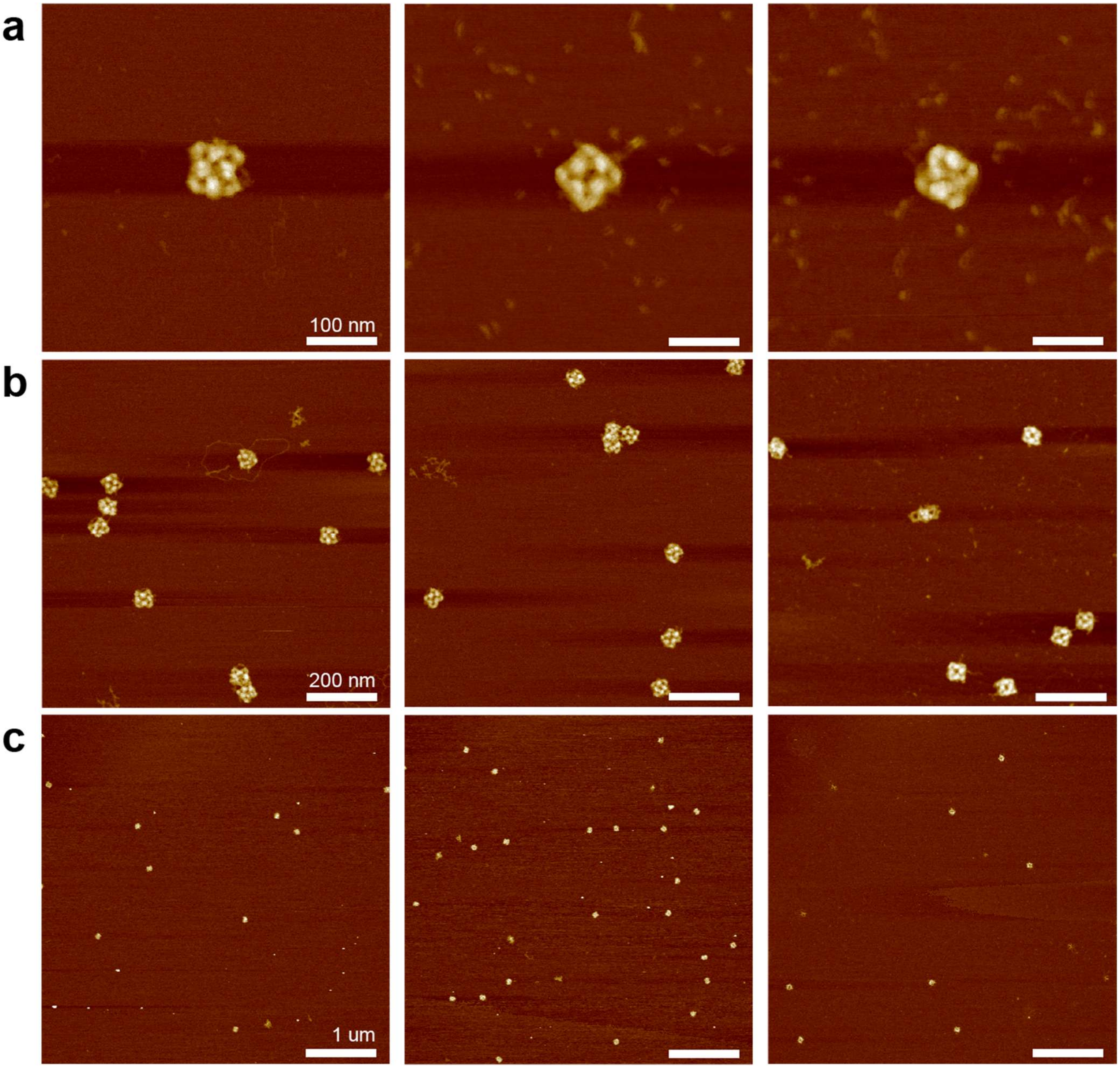


**Figure S17. AFM images of cuboctahedra with long jack staples reconfigured from octahedra.** Scale bars: **a** 100 nm, **b** 200 nm, **c** 1 µm.

### S5. Additional TEM Images

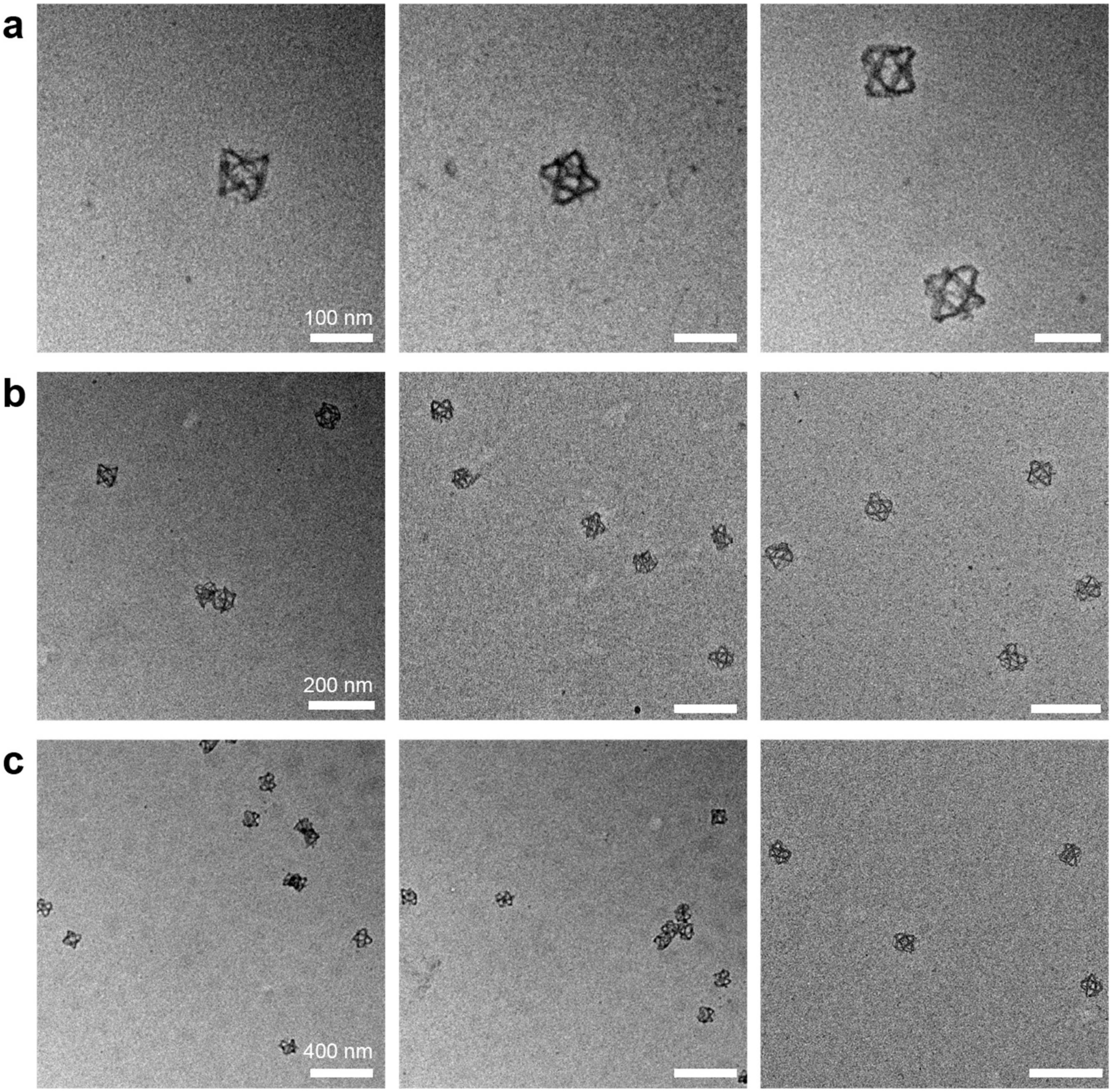


**Figure S18. TEM images of initial cuboctahedra without jack staples.** Scale bars: **a** 100 nm, **b** 200 nm, **c** 400 nm.

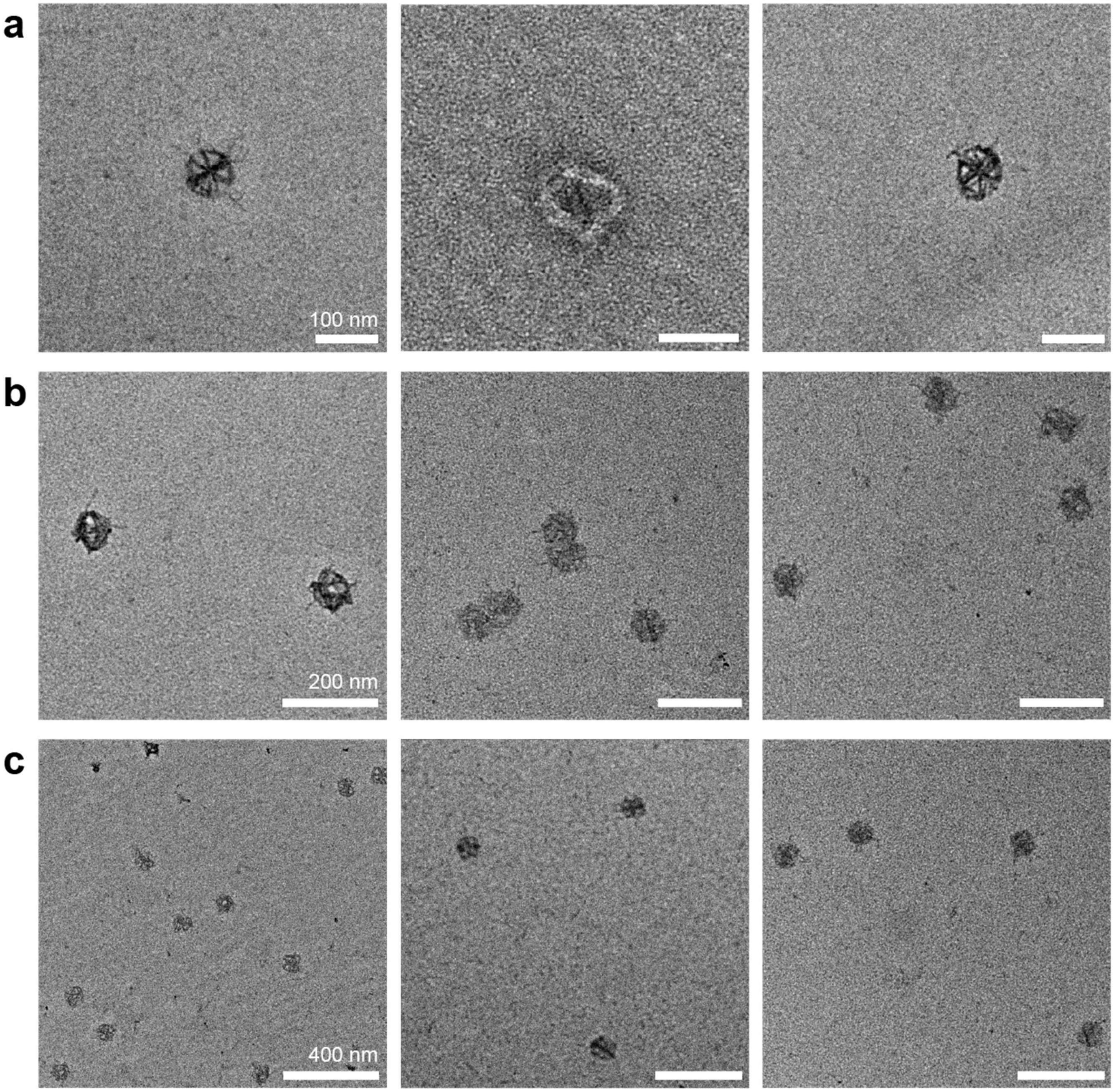


**Figure S19. TEM images of octahedra with short jack staples.** Scale bars: **a** 100 nm, **b** 200 nm, **c** 400 nm.

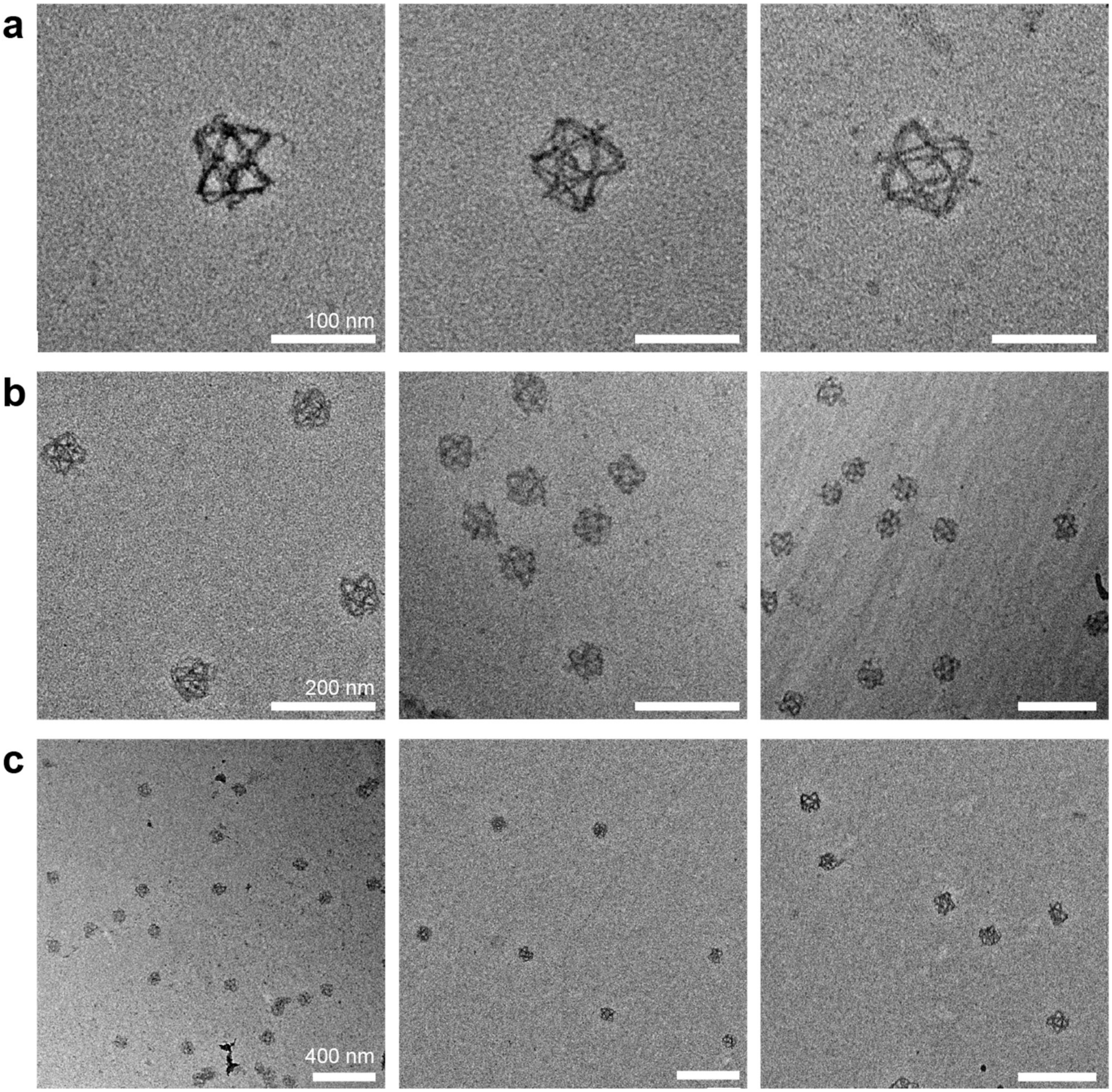


**Figure S20. TEM images of cuboctahedra with long jack staples.** Scale bar: **a** 100 nm**, b** 200 nm**, c** 400 nm**.**

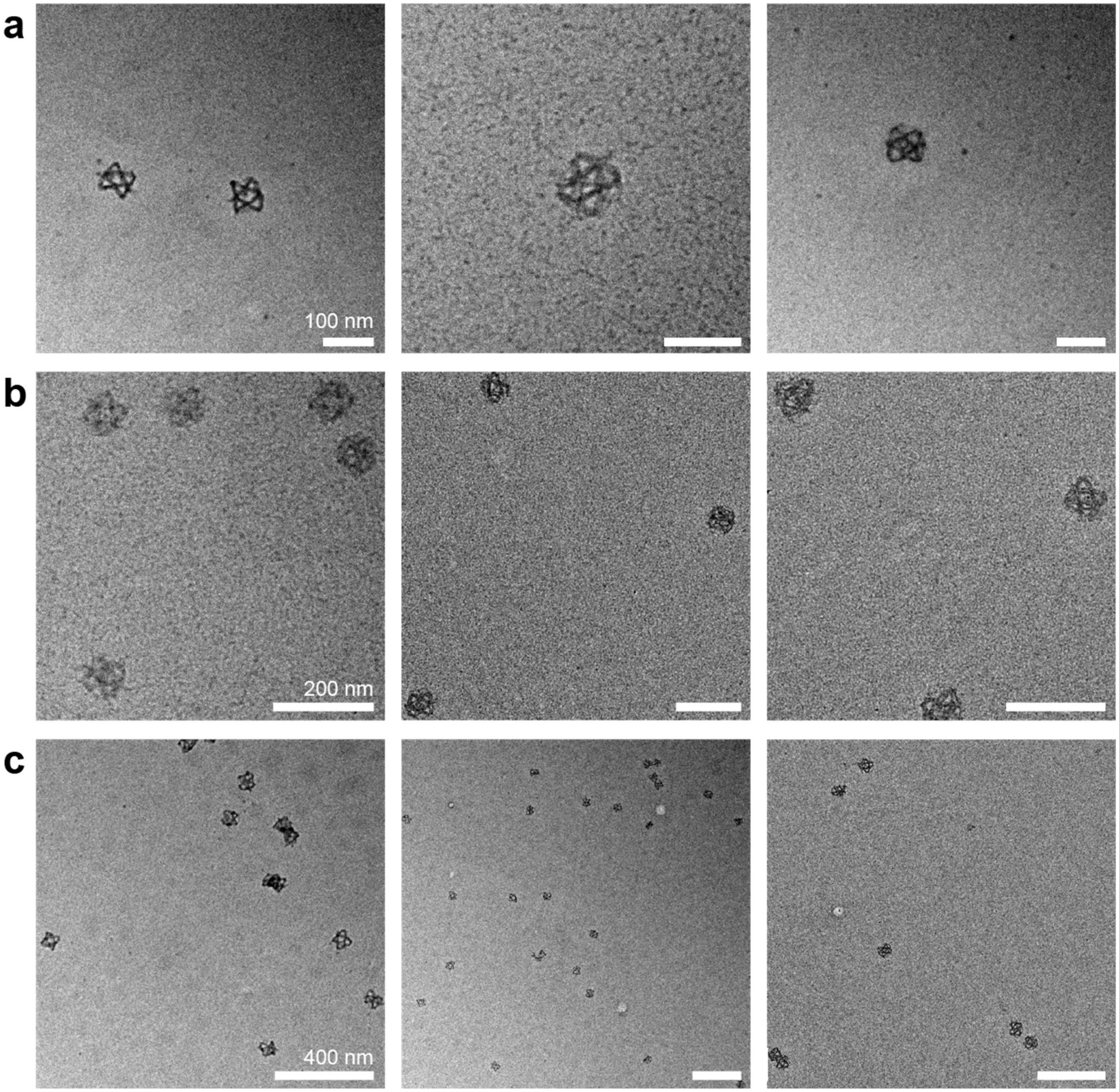


**Figure S21. TEM images of reconfigured cuboctahedra without jack staples.** Scale bar: **a** 100 nm**, b** 200 nm**, c** 400 nm**.**

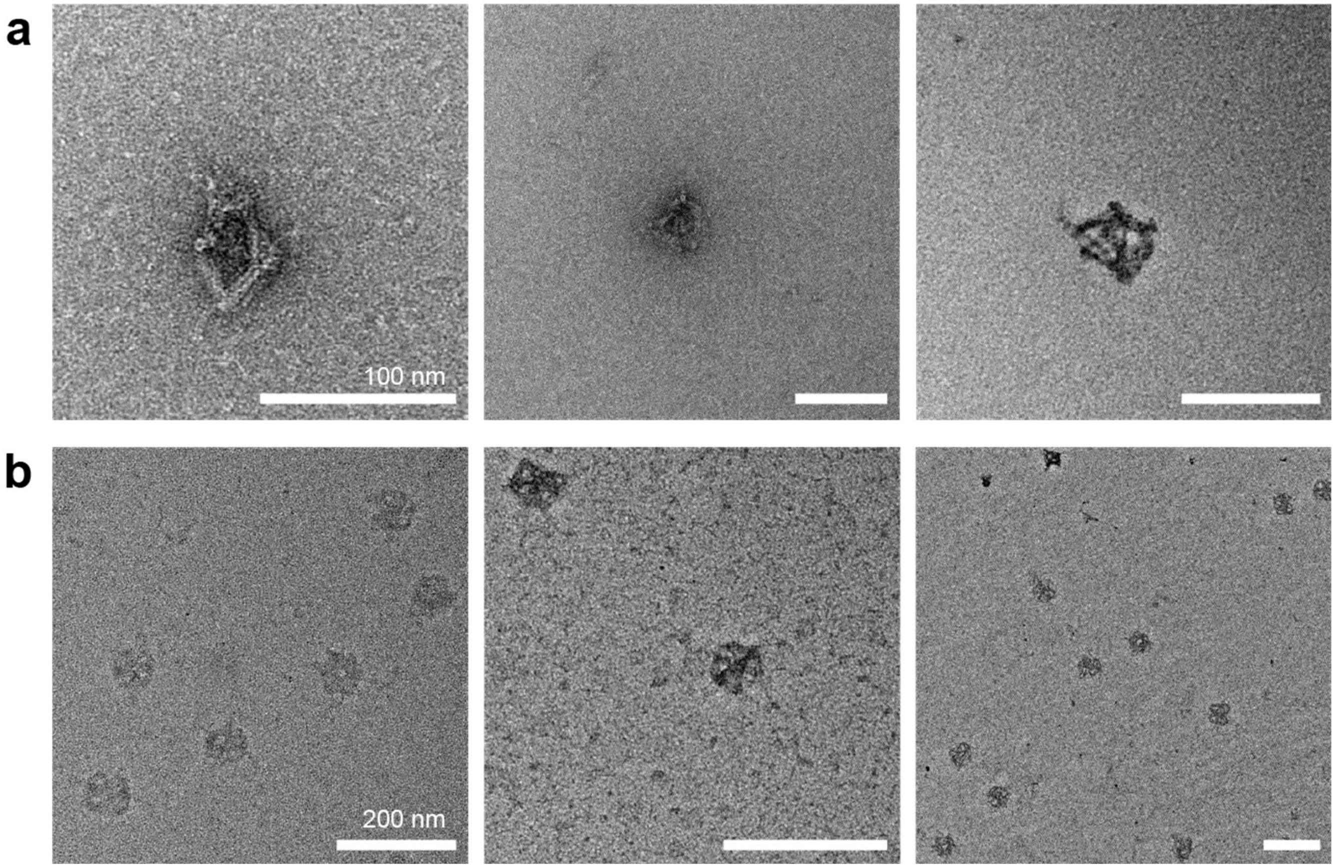


**Figure S22. TEM images of octahedra with short jack staples reconfigured from cuboctahedra with long jacks.** Scale bar: **a** 100 nm**, b** 200 nm**.**

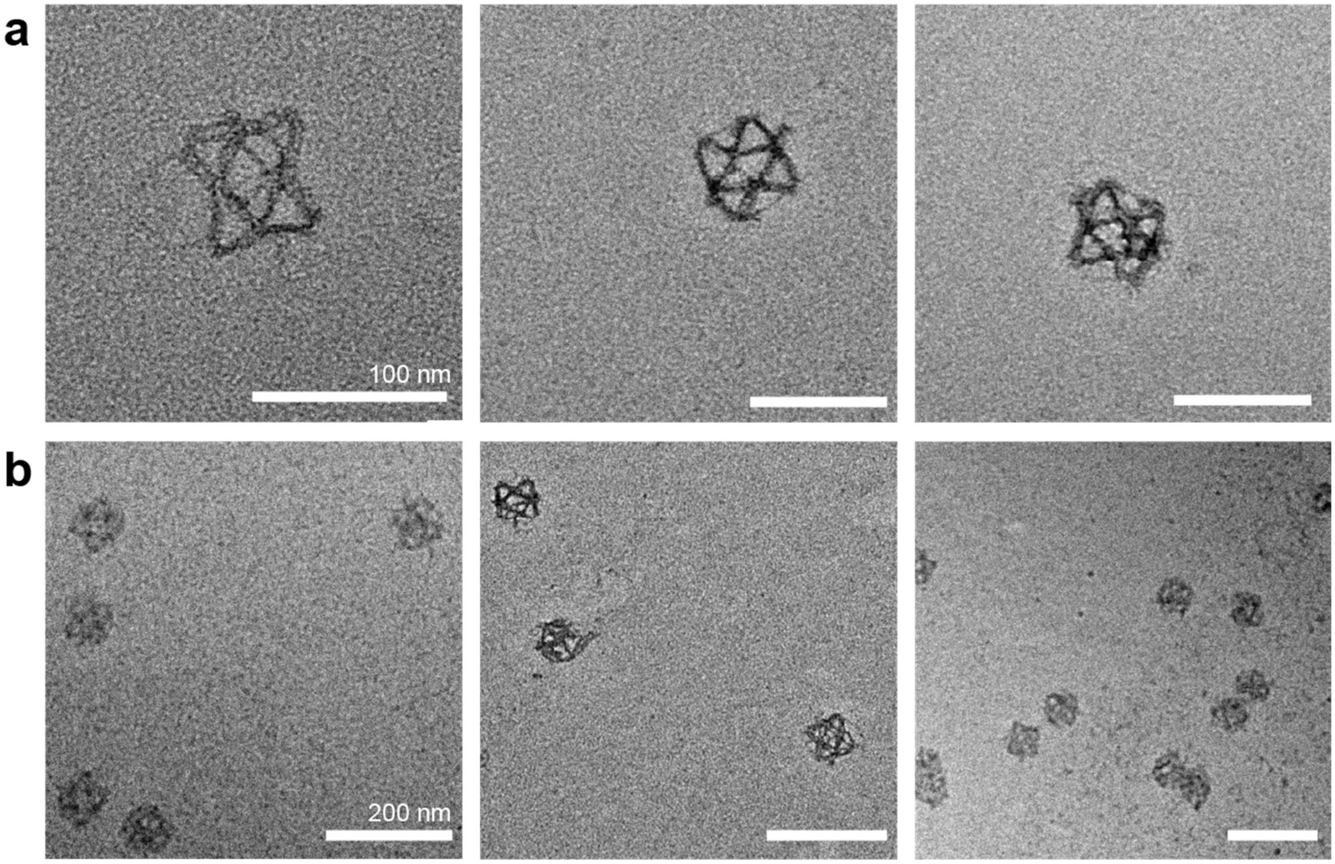


**Figure S23. TEM images of cuboctahedra with long jack staples reconfigured from octahedra.** Scale bar: **a** 100 nm**, b** 200 nm**.**

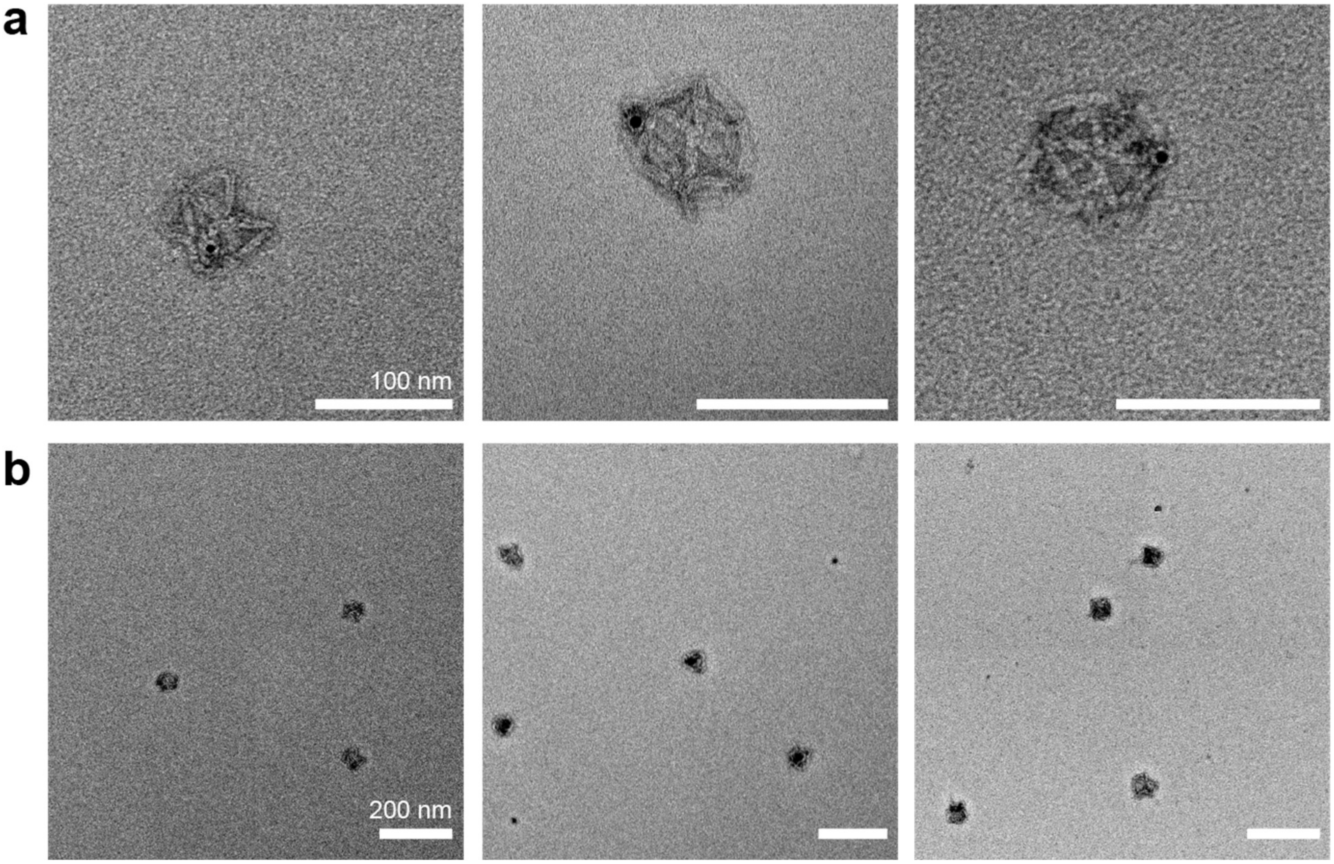


**Figure S24. TEM images of cuboctahedra encapsulating 5-nm AuNPs.** Scale bar: **a** 100 nm**, b** 200 nm**.**

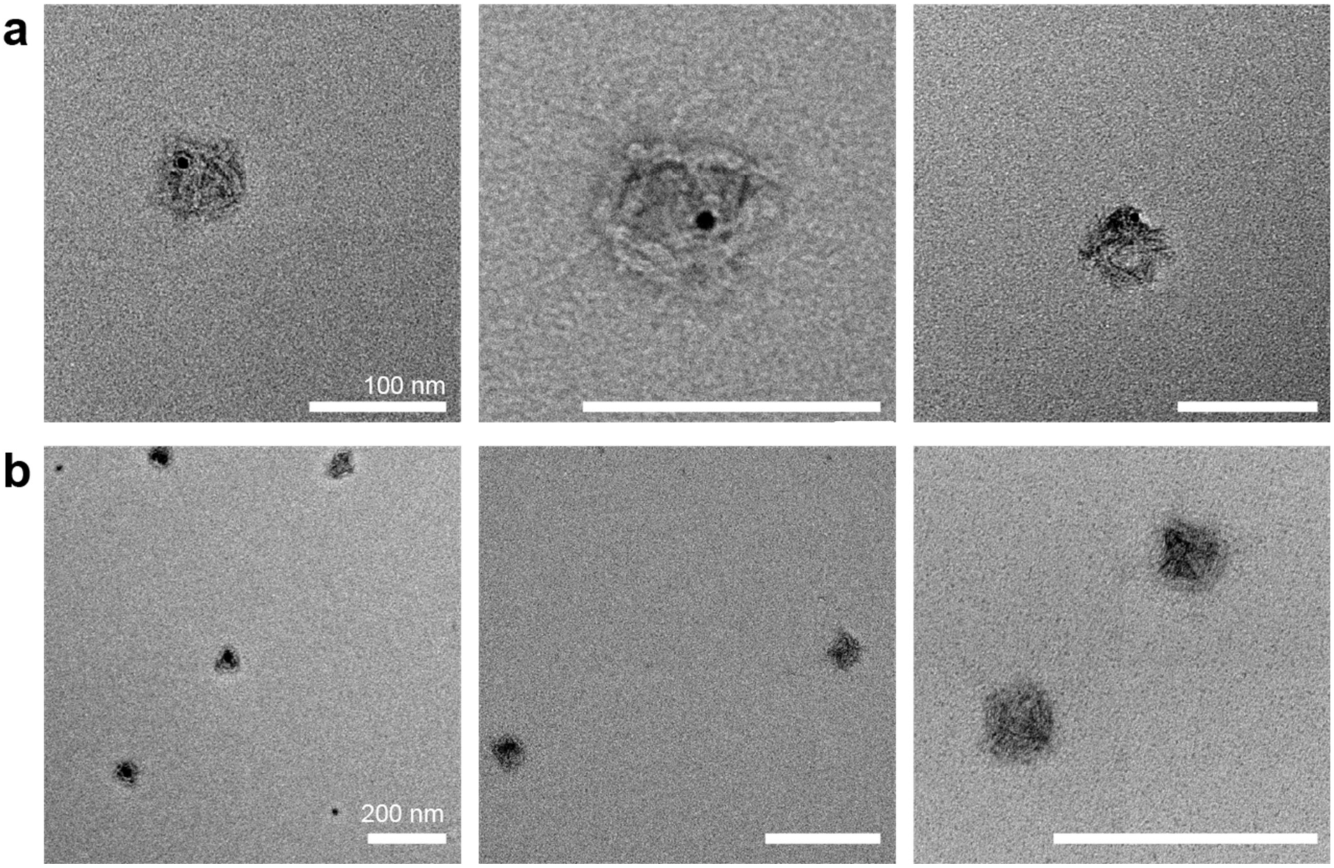


**Figure S25. TEM images of octahedra encapsulating 5-nm AuNPs**. Scale bar: **a** 100 nm**, b** 200 nm**.**

### S6. Reconfiguration Induced by External Stimuli

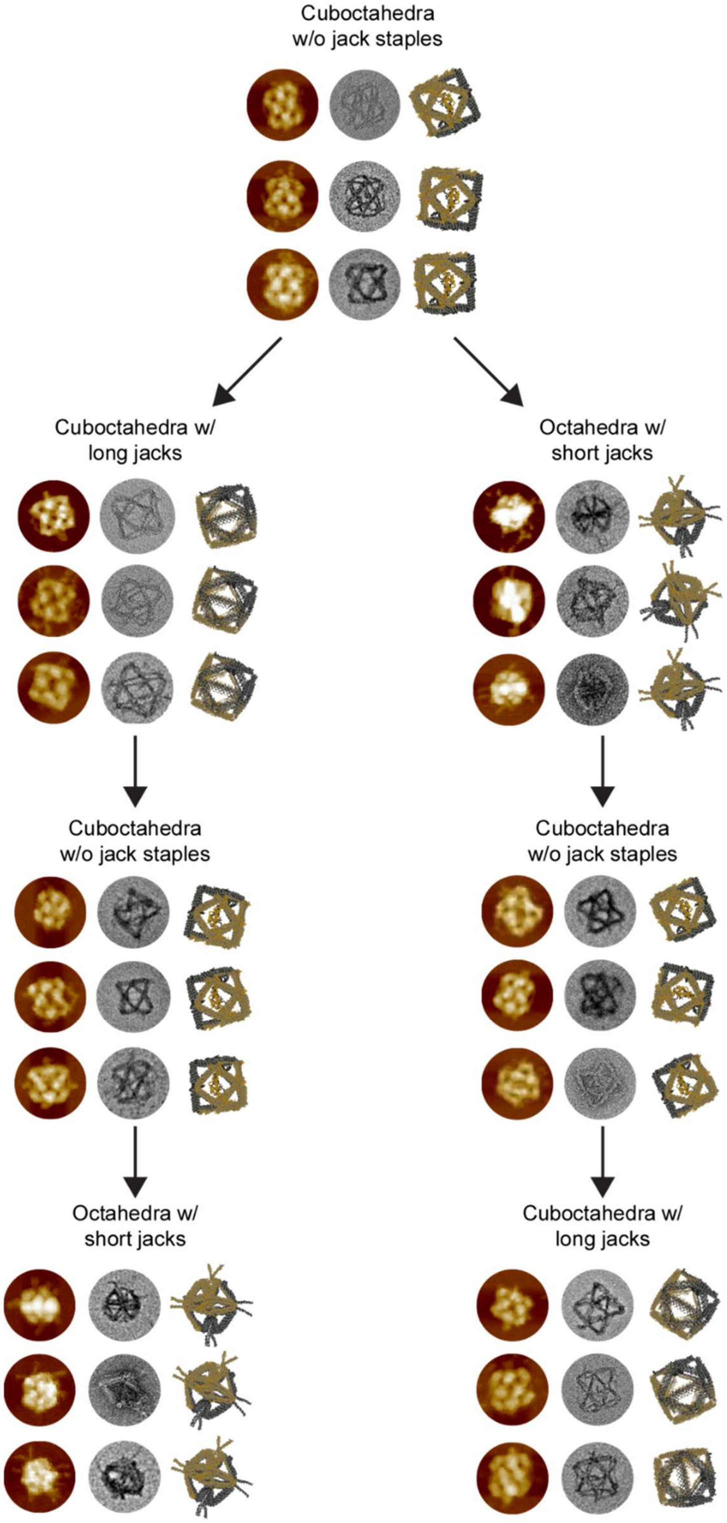


**Figure S26. Reconfiguration via TMSD and reannealing.** Two parallel reconfiguration pathways are demonstrated, both originating from the initial cuboctahedral configuration without jack staples.

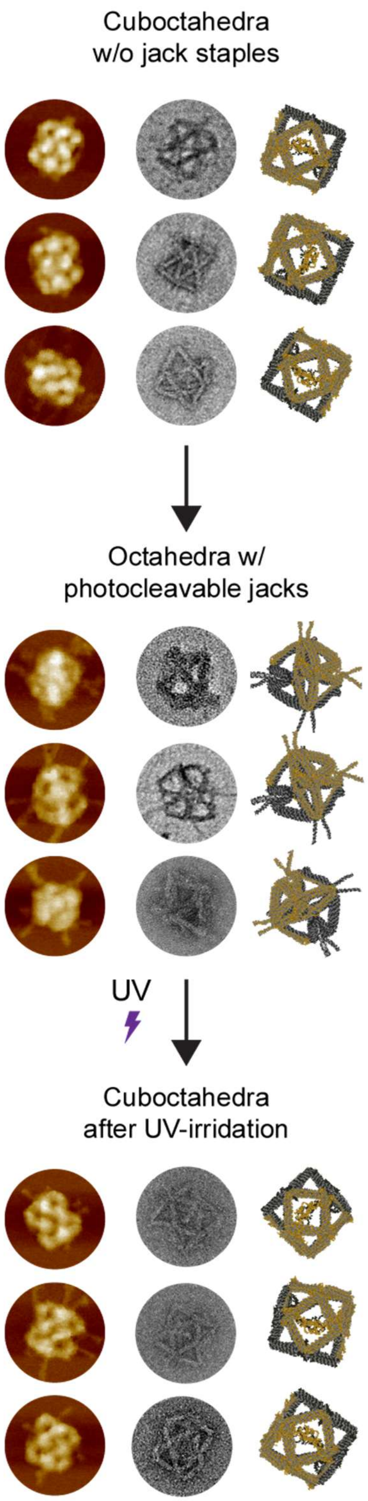


**Figure S27. UV-induced structural transformation.** Photocleavable reconfiguration pathway initiated from cuboctahedral structures without jack staples. UV-responsive octahedra were assembled using photocleavable jack staples. Upon UV irradiation, photocleavage of the staples induced a spontaneous transition into cuboctahedral geometry.

**S7. DLS Measurements**

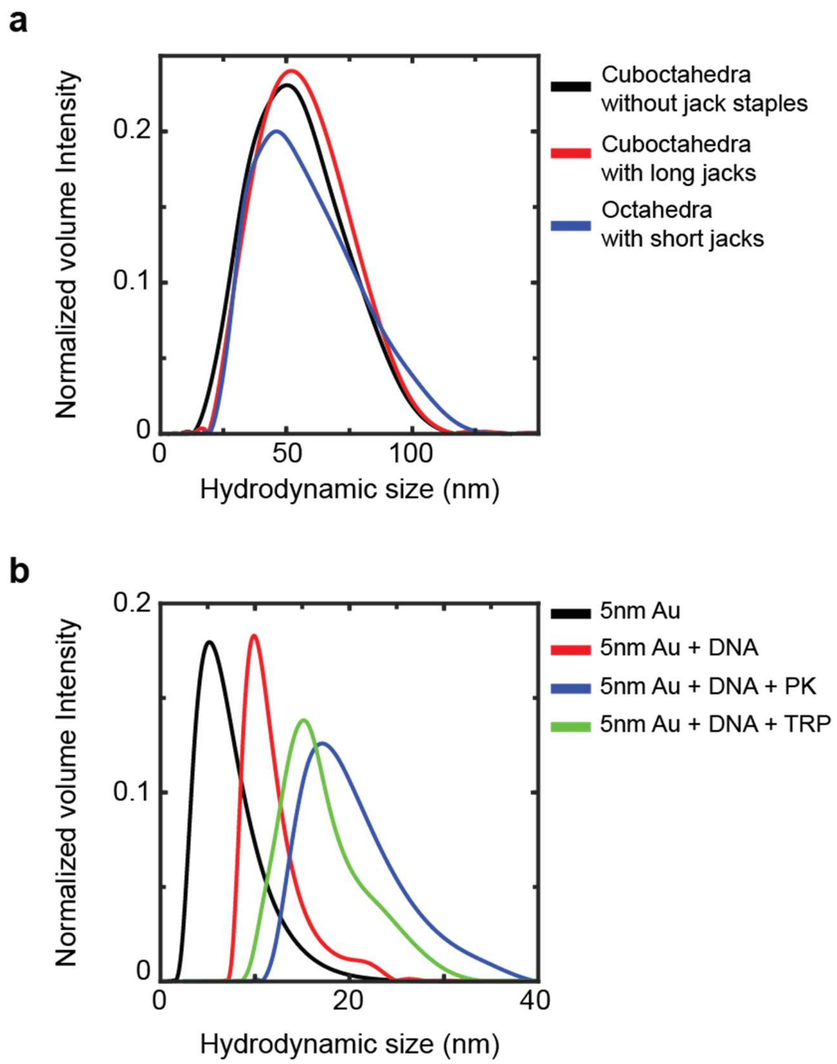


**Figure S28. DLS measurements of DNA nanostructures and AuNPs. a,** Hydrodynamic diameter distributions of three distinct configurations: cuboctahedra without jack staples (black, peak at ~50 nm), cuboctahedra with long jack staples (red, peak at ~52 nm), and octahedra with short jack staples (blue, peak at ~46 nm). **b,** Hydrodynamic diameter distributions of functionalized AuNPs: bare 5-nm citrate-stabilized AuNPs (black), DNA-functionalized AuNPs with thiolated oligonucleotides (red, peak at ~10 nm), DNA-AuNP conjugates with PK (blue, peak at ~16 nm), and DNA-AuNP conjugates with trypsin (green, peak at ~14 nm).

### S8. Gel Electrophoresis and Structural Yield Estimation

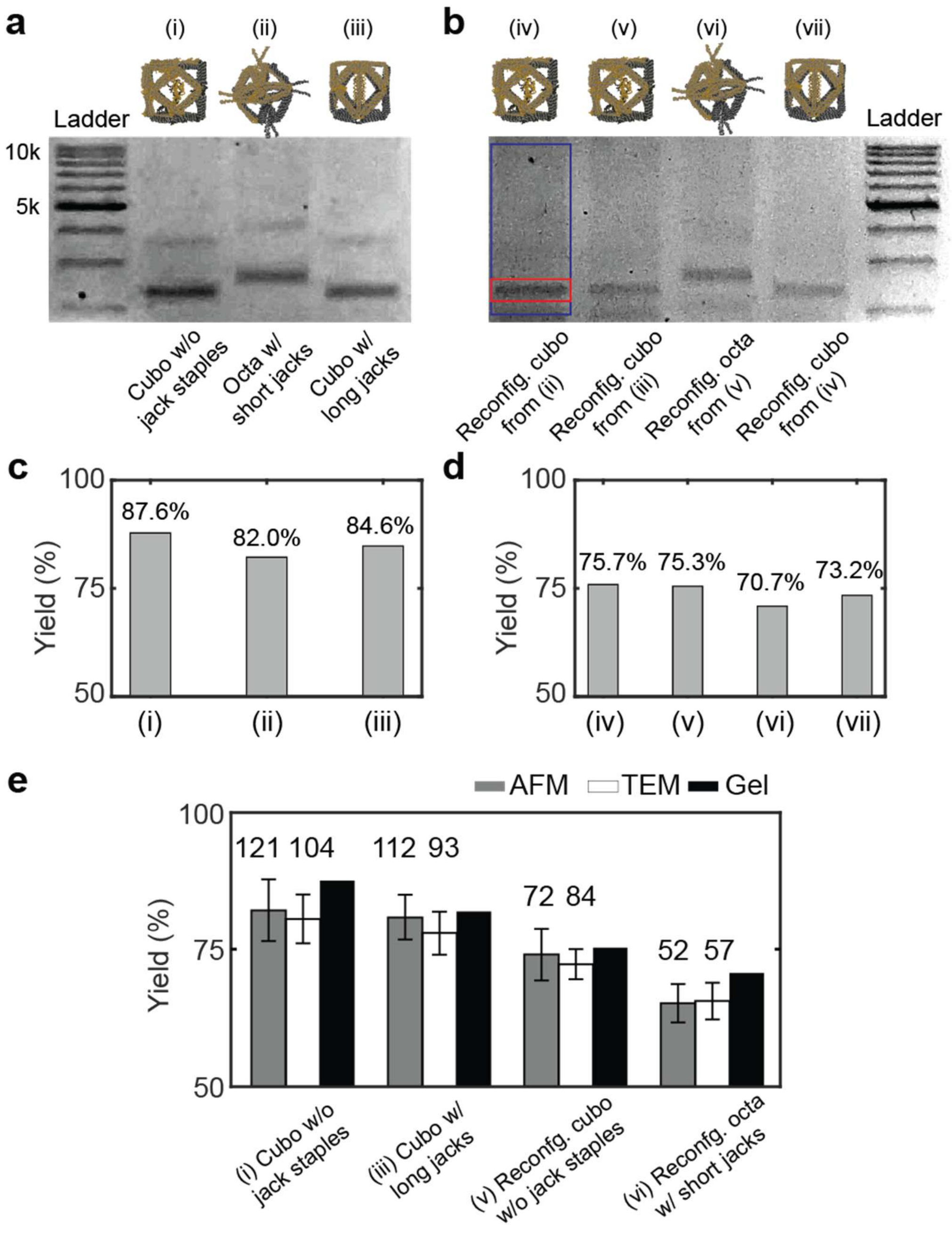


**Figure S29. Agarose gel electrophoresis analysis and structural yield**. **a,** Gel image of DNA nanostructures corresponding to Fig. 2c: (i) cuboctahedra without jack staples, (ii) octahedra with short jack staples, (iii) cuboctahedra with long jack staples. **b,** Gel image of reconfigured DNA nanostructures via TMSD: (iv) cuboctahedra reconfigured from octahedra, (v) cuboctahedra reconfigured from cuboctahedra with long jacks, (vi) octahedra reconfigured from cuboctahedra, and (vii) cuboctahedra reconfigured from cuboctahedra without jack staples. **c,** Structural formation yields for initial configurations (i)–(iii) determined from gel band intensities. **d,** Formation yields for reconfigured structures (iv)–(vii). **e,** Experimental yields of the Jitterbug structures in a series of reconfiguration, estimated using AFM, TEM, and agarose gel electrophoresis. The transformation began with initial cuboctahedron (without jack staples) which were maintained with extended jacks. The structure was then reconfigured into cuboctahedron without jacks via TMSD using long-jack releaser strands. Finally, octahedral structure was formed with short jack staples. The number of DNA structures analyzed for yield estimation is indicated.

The yield per lane was evaluated using ImageJ software. Yield (%) was calculated as[7, 12]:

$$Yield\ (\%) = \frac{Band\ intensity}{Lane\ intensity}$$

Band and lane intensities were estimated after background subtraction with the rolling ball method[13] (using a radius similar to well width) and conversion to grayscale.

### S9. FRET Measurements

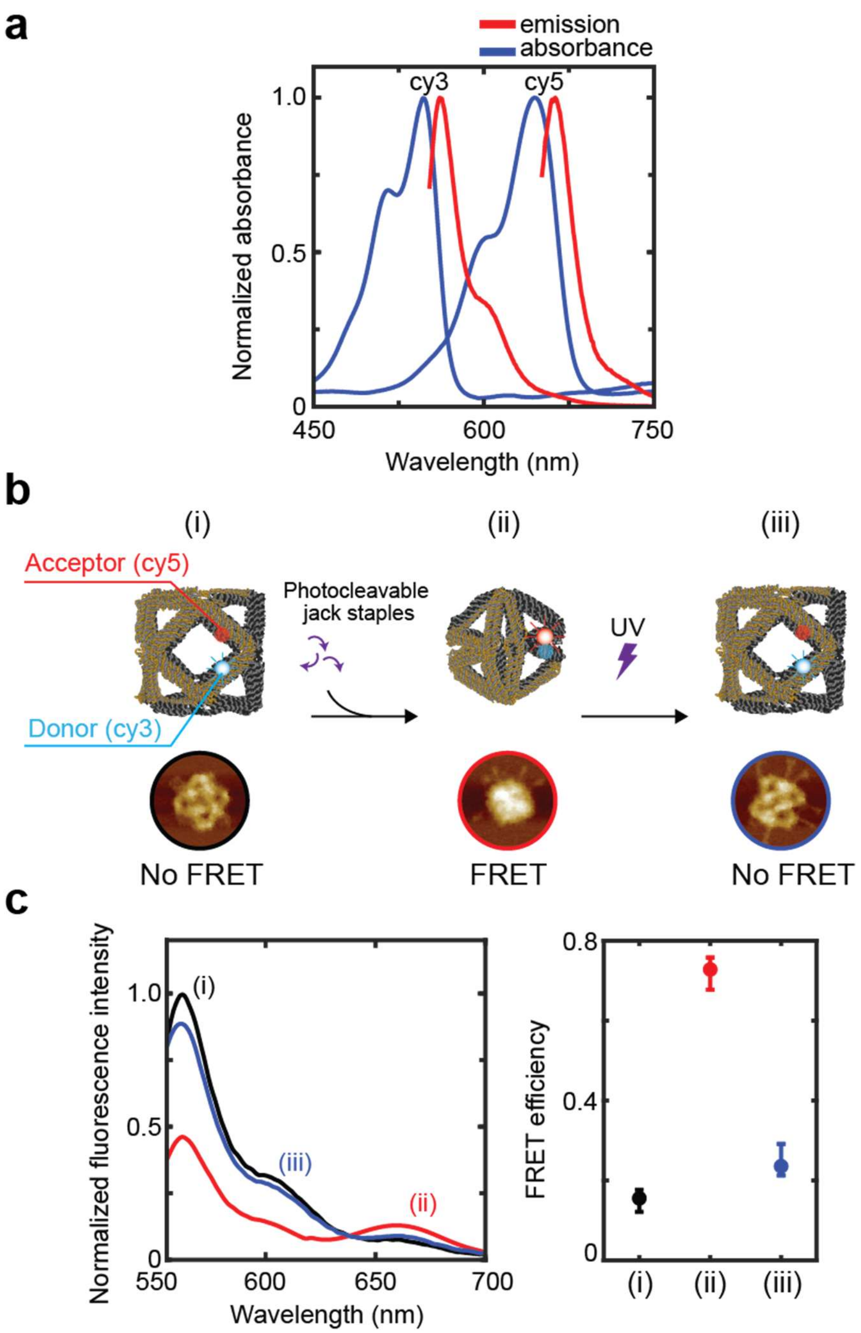


**Figure S30. FRET analysis during reconfiguration. a,** Absorption and emission spectra of FRET pair components. Donor fluorophore (Cy3) exhibits absorption maximum at ~547 nm and emission maximum at ~562 nm, while acceptor fluorophore (Cy5) shows absorption and emission maxima at ~645 nm and ~662 nm, respectively. **b,** FRET response during UV-induced reconfiguration: (i) initial cuboctahedral state without jack staples, (ii) octahedral conformation with photocleavable short jack staples, and (iii) cuboctahedral state following UV-induced jack staple cleavage. **c,** Fluorescence spectra and FRET efficiency. Donor fluorescence intensity decreased from 1.0 to ~0.45 upon octahedral transformation and recovered to ~0.83 following UV-induced reconfiguration. Corresponding FRET efficiencies were approximately 0.18 (initial cuboctahedra), 0.77 (octahedra), and 0.25 (final cuboctahedra), respectively.

Among multiple methods[14-18] calculating FRET efficiency, the approach developed by Hoppe et al.[17] was used. The calculation assumes constant mean quantum efficiency of the fluorophores, and that fluorescence emission can be transferred between different filter combinations using scalar factors to correct for excitation and emission contributions[17, 19]. Contributions from excitation light and fluorescence emission were corrected using scalar factors. Under these assumptions, FRET efficiency (E) was determined as:

$$E = (\frac{I_F - \beta I_D}{\alpha I_A} - 1)(\frac{1}{f_A})$$

where $I_A$ is the intensity at the acceptor excitation and acceptor emission, $I_D$, is the intensity at the donor excitation and donor emission, and $I_F$ is the intensity at the donor excitation and acceptor emission. $\alpha$ is a proportionality constant relating acceptor fluorescence at the acceptor excitation to the donor excitation $(I_F/I_A)$ , $\beta$ is a proportionality constant relating donor fluorescence detected at the acceptor emission relative to that detected at the donor emission $(I_F/I_D)$, and $f_A$ is the ratio of concentration of donor-acceptor complex to total concentration of the acceptor (free plus complexed).

### S10. Absorbance Spectra of AuNPs

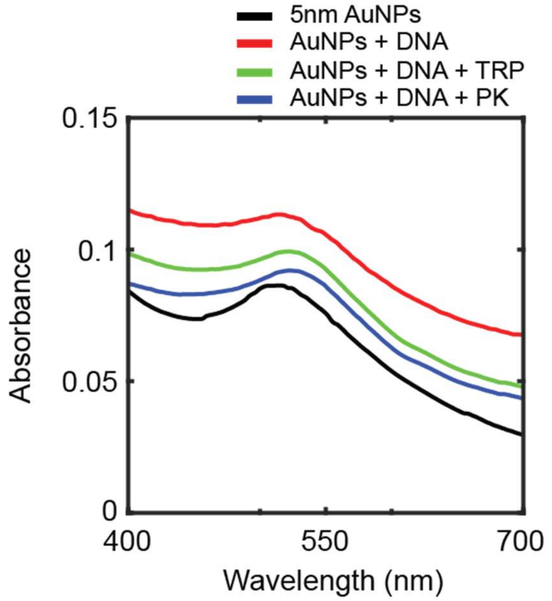


**Figure S31. Absorption spectra of functionalized AuNPs.** Progressive spectral shifts are observed upon sequential functionalization: citrate-stabilized 5-nm AuNPs (black, $\lambda_{max}$ = ~513 nm), AuNPs conjugated with thiolated DNA strands (red, $\lambda_{max}$ = ~515 nm), AuNPs functionalized with DNA and trypsin (green, $\lambda_{max}$ = ~522 nm), and AuNPs decorated with DNA and proteinase K (blue, $\lambda_{max}$ = ~522 nm). The red shift in the surface plasmon resonance peak position indicates successful surface modification and increased local refractive index upon biomolecule conjugation.

The number of DNA strands conjugated to each gold nanoparticle was quantified following previously reported methods[2, 20, 21]. Since the thiolated DNA oligonucleotides were not labeled with any dyes, their concentrations were determined by UV-visible absorbance measurements using extinction coefficients calculated with OligoAnalyzer software (Integrated DNA Technologies), consistent with manufacturer specifications.

Following conjugation and removal of excess DNA strands, bound oligonucleotides were displaced from the AuNPs by overnight incubation with 1 M DTT in PBS buffer at room temperature. The resulting solution was centrifuged at 10,000× g for 15 minutes to pellet the AuNPs. Released oligonucleotides in the supernatant were purified using NAP-5 size-exclusion columns (Sephadex G-25, GE Healthcare) and reconstituted in fresh TAE buffer. DNA concentrations were determined by comparing absorbance values at 260 nm to standard calibration curves prepared with known oligonucleotide concentrations in identical buffer conditions. This analysis indicated an average conjugation density of approximately 14 DNA strands per 5-nm gold nanoparticle.

#### S11. Control Experiments for Nanopore Formation on Synthetic Vesicles

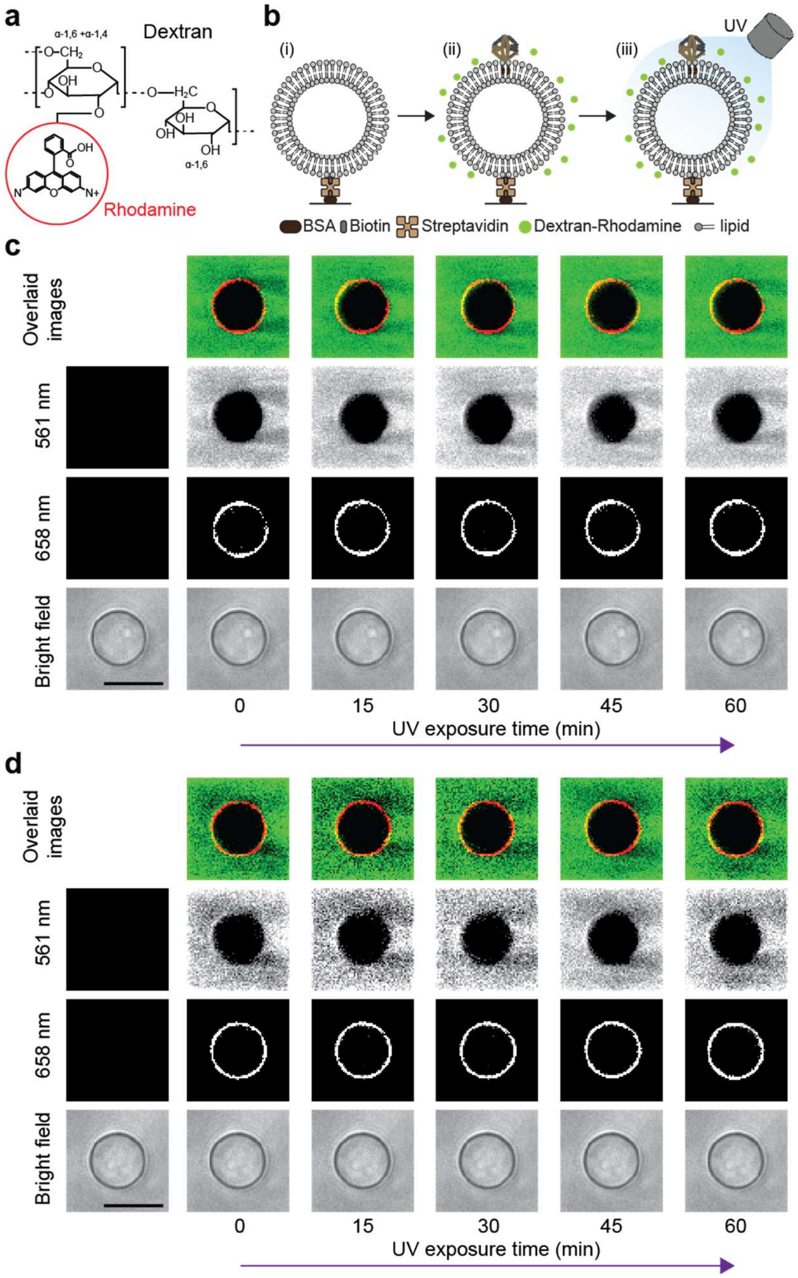


**Figure S32. Control experiments for nanopore formation on synthetic vesicles. a,** Molecular structure of dextran-rhodamine (DR). **b,** Experimental procedure. (i) Giant vesicles containing biotinylated lipids were immobilized on streptavidin-passivated microfluidic imaging chamber via biotin-streptavidin interactions using surface-bound biotinylated BSA. (ii) Vesicles were bound with octahedral Jitterbug structures (Cy5-labeled, lacking photocleavable linkers) through cholesterol-modified anchoring, followed by introduction of 10 kDa DR molecules into the channel. (iii) After 60 minutes of UV irradiation, DR remained excluded from the vesicle, confirming membrane integrity. **c** and **d,** Representative images of giant vesicles under brightfield illumination, Cy5-labeled Jitterbug fluorescence (658 nm excitation), and 10 kDa DR fluorescence (561 nm excitation). Composite images display DR in green pseudo color and Cy5-Jitterbug in red pseudo color. Scale bars: 10 μm for all brightfield images.

**S12. Nanopore Formation and Recovery on Giant Vesicles**

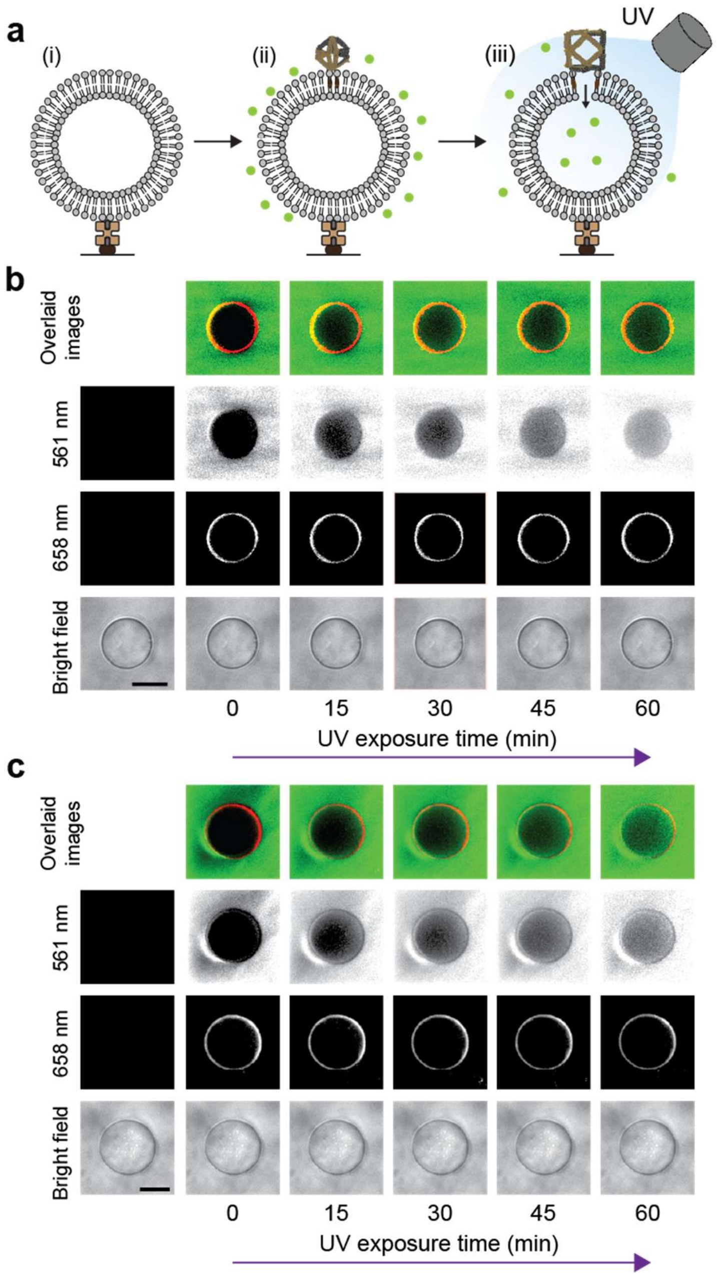


**Figure S33. Controlled membrane permeability experiment with 10 kDa DR**. **a,** Workflow for evaluating nanopore formation. (i) Giant vesicles were immobilized on the surface of an imaging chamber. (ii) UV-resposnive octahedra (Cy5-labeled) bound to the vesicles using cholesterol taggs, followed by introduction of 10 kDa DR molecules into the channel. (iii) After UV irradiation (60 minutes), DR entered the the vesicle, confirming successful nanopore formation and membrane permeability. **b** and **c,** Representative images of giant vesicles under brightfield illumination, Cy5-labeled Jitterbug fluorescence (658 nm excitation), and 10 kDa DR fluorescence (561 nm excitation). Composite images show DR in green pseudo color and Cy5-Jitterbug in red pseudo color. Scale bars: 10 µm for all brightfield images.

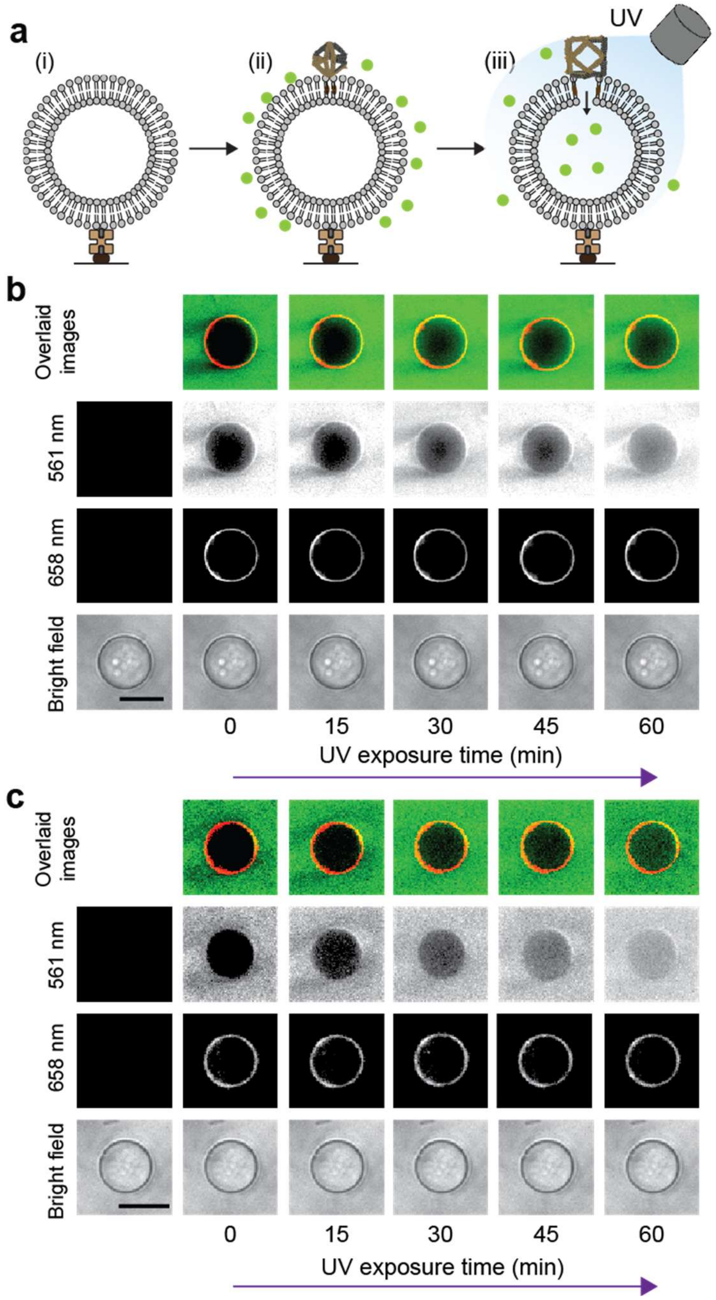


**Figure S34. Nanopore formation experiment with 40 kDa DR**. **a,** Experimental procedure. (i) Vesicles were immobilized on the microfluidic channel via biotin-streptavidin interactions. (ii) Cy5-labeled DNA transformers with photocleavable jack staples were introduced and bound to vesicles through cholesterol-modified DNA anchoring. 40 kDa DR molecules were also introduced into the chamber. (iii) After 60 minutes of UV exposure, DR entered the vesicle interior, confirming nanopore formation and membrane permeability. **b** and **c,** Microscopy images of giant vesicles under brightfield illumination, Cy5-labeled Jitterbug fluorescence (658 nm excitation), and 40 kDa DR emission (561 nm excitation). Overlaid images show DR in green color and Cy5-Jitterbug in red. Scale bars: 10 μm.

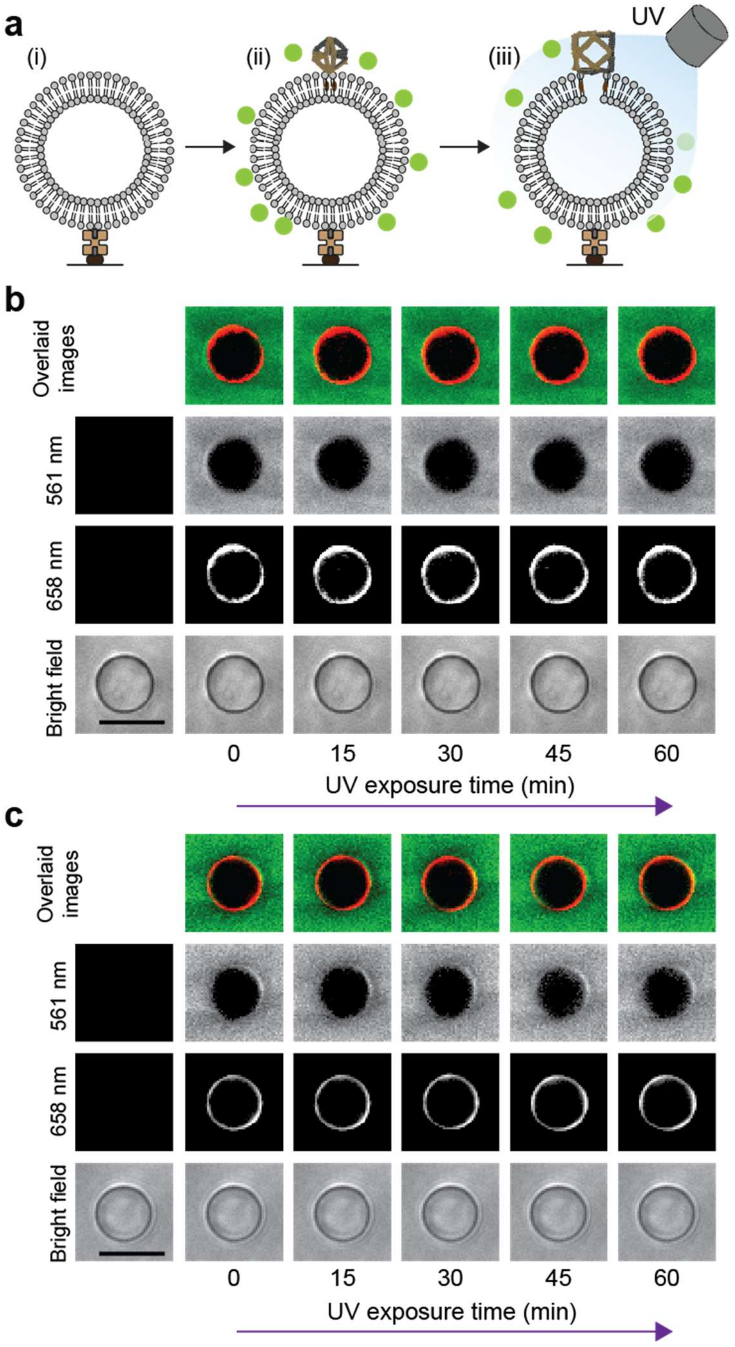


**Figure S35. Size-selective permeability experiment. a,** Workflow for evaluating size-seletive membrane permeability. (i) Giant vesicles were immobilized on the surface-passivated microfluidic chamber. (ii) Octahedral DNA bound to vesicles via cholesterol groups. 70 kDa DR molecules were introduced into the channel. (iii) After UV irradiation, DR remained excluded from the vesicle interior, suggesting that 70 kDa molecues were too large to pass through the nanopores. **b** and **c,** Images of vesicles under brightfield illumination, Cy5-labeled Jitterbug fluorescence (658 nm excitation), and 70 kDa DR fluorescence (561 nm excitation). Overlaid images display DR in green and Jitterbug in red. Scale bars: 10 µm.

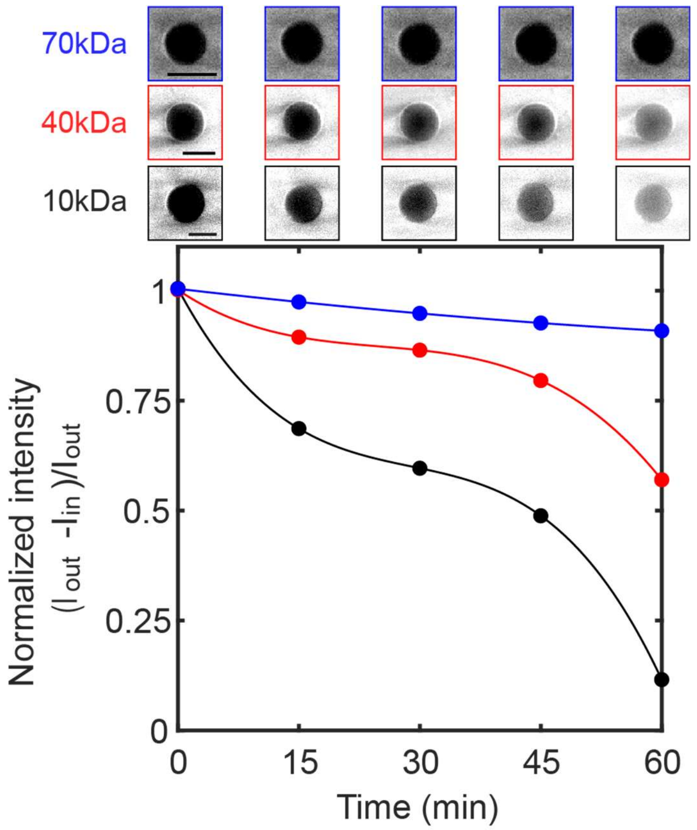


**Figure S36. Size-dependent DR influx into vesicles.** DR fluorescence intensity (561 nm excitation) was measured at 0, 15, 30, 45, and 60 minutes. Vesicle images were captured for 10, 40, and 70 kDa DR at each time point. Following 60-minute UV exposure, 10 kDa DR entered vesicles more rapidly and in a greater amount than 40 kDa DR, while 70 kDa DR showed negligible influx. $I_{out}$ and $I_{in}$ represent DR fluorescence intensity outside and inside vesicles, respectively. Scale bars: 10 μm.

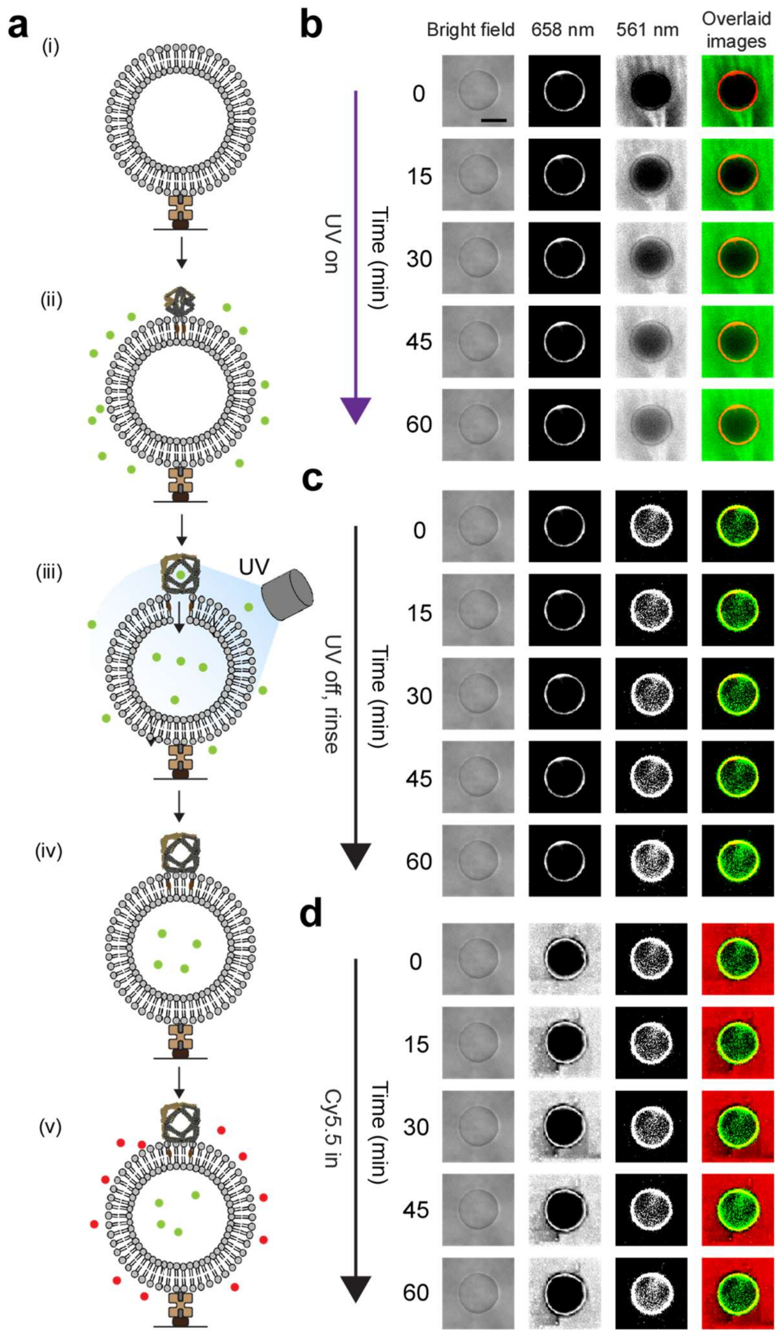


**Figure S37. Formation and recovery of nanopores. a,** (i) Vesicles were immobilized on the surface of an imaging channel. (ii) UV-activable octahedra and 10 kDa DR were introduced into the channel. (iii) DR molecules entered the vesicles during the 60-minute UV irradiation. (iv) In the next 60 minutes after UV light off, the channel was rinsed and extravesicular DR molecules were removed. (v) Cy5.5 dyes were subsequently introduced into the channel. **b-d,** Images of vesicles under brightfield illumination, Jitterbug fluorescence (658 nm excitation), and DR fluorescence (561 nm excitation). Overlaid images present DR in green color, Cy5-Jitterbug in red, and Cy5.5 dyes in red. **b,** DR entry into vesicles was monitored during continuous UV irradiation. **c,** After UV light turned off and extravesicular DR removed, no membrane leakage was observed, indicating a nanopore closure. **d,** Subsequent introduction of Cy5.5 fluorophores demonstrated their exclusion from the vesicles. The results indicate a complete restoration of membrane integrity. Scale bars represent 10 µm.

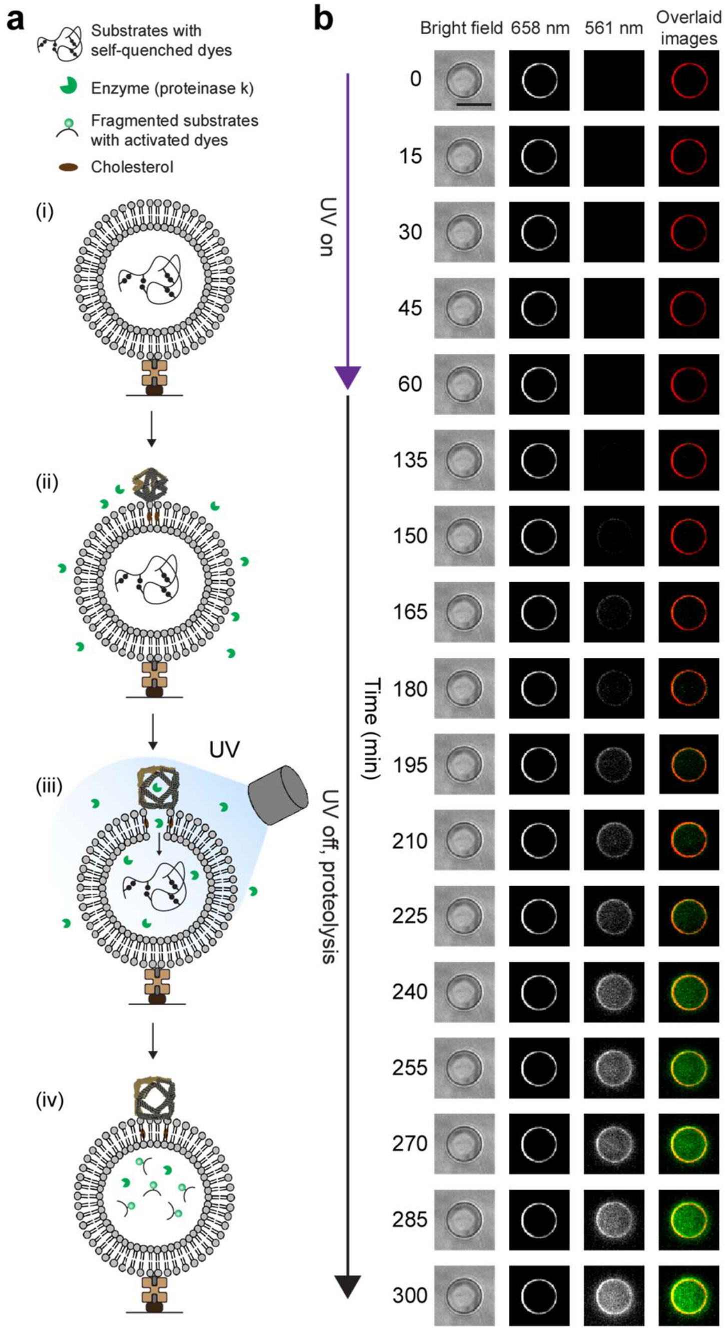


**Figure S38. UV-triggered nanopore formation and proteolytic reactions in giant vesicles. a,** Experimental procedure. (i) Vesicles containing self-quenched fluorescent casein-Bodipy substrate were immobilized on the microfluidic imaging chamber. (ii) Jitterbug DNA and PK enzyme solution were introduced into the channel. (iii) Vesicles were subjected to 60 minutes of UV irradiation to induce DNA transformation and nanopore formation, allowing PK to enter the vesicle cavity and digest its substrate. (iv) After UV light turned off, the channel was washed with buffer to remove extravesicular PK. Nanopores were resealed and intravesicular proteolysis was subsequently monitored. **b,** Images of vesicles under brightfield illumination, Jitterbug fluorescence (Cy5 fluorescence excited at 658 nm), and PK-digested substrate emission (excited at 561 nm). Composite images show Bodipy (PK activity) in green and Jitterbug in red Scale bars represent 10 μm.

### S13. Enzyme Transport into Vesicles

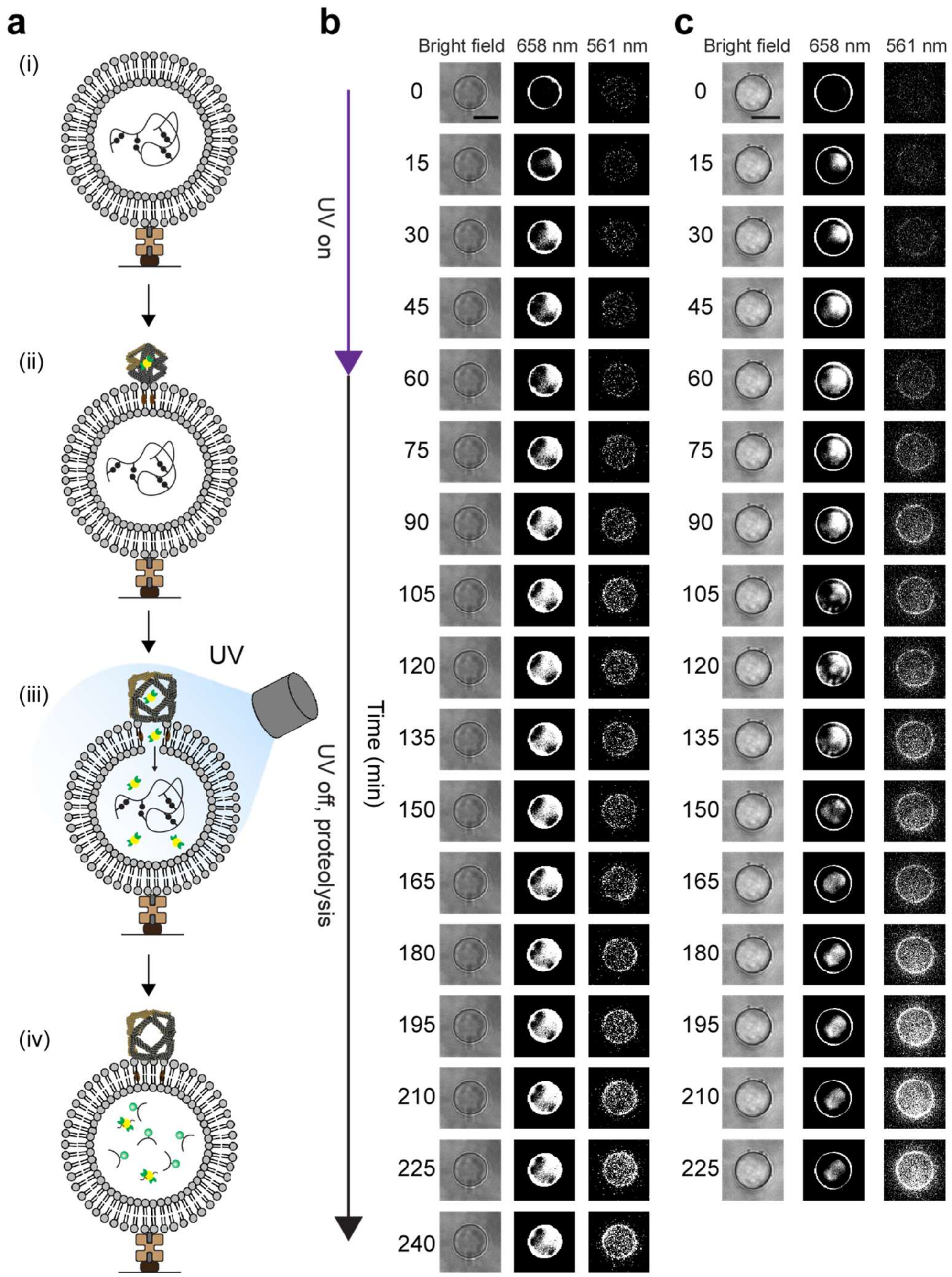


**Figure S39. Two sets of experiments of Au-enzyme transport and proteolysis within vesicles (substrate: Casein, enzyme: PK). a,** Experimental workflow. (i) Giant vesicles containing Bodipy-conjugated casein were immobilized on surface. (ii) DNA octahedra encapsulating Au-enzyme complexes were bound to the vesicle membrane. (iii) UV irradiation induces Jitterbug transformation, generating nanopores that enabled release of Au-enzyme complexes into the vesicle. (iv) Following nanopore closure, intravesicular proteolysis was monitored by time-lapse fluorescence increase from substrate cleavage. **b** and **c,** Microscopy images of vesicles under brightfield illumination, 658 nm excitation (Cy5-labeled Jitterbug fluorescence and Cy5.5-tagged Au-enzyme fluorescence), and 561 nm excitation (Bodipy fluorescence). Scale bars represent 10 μm in all brightfield images.

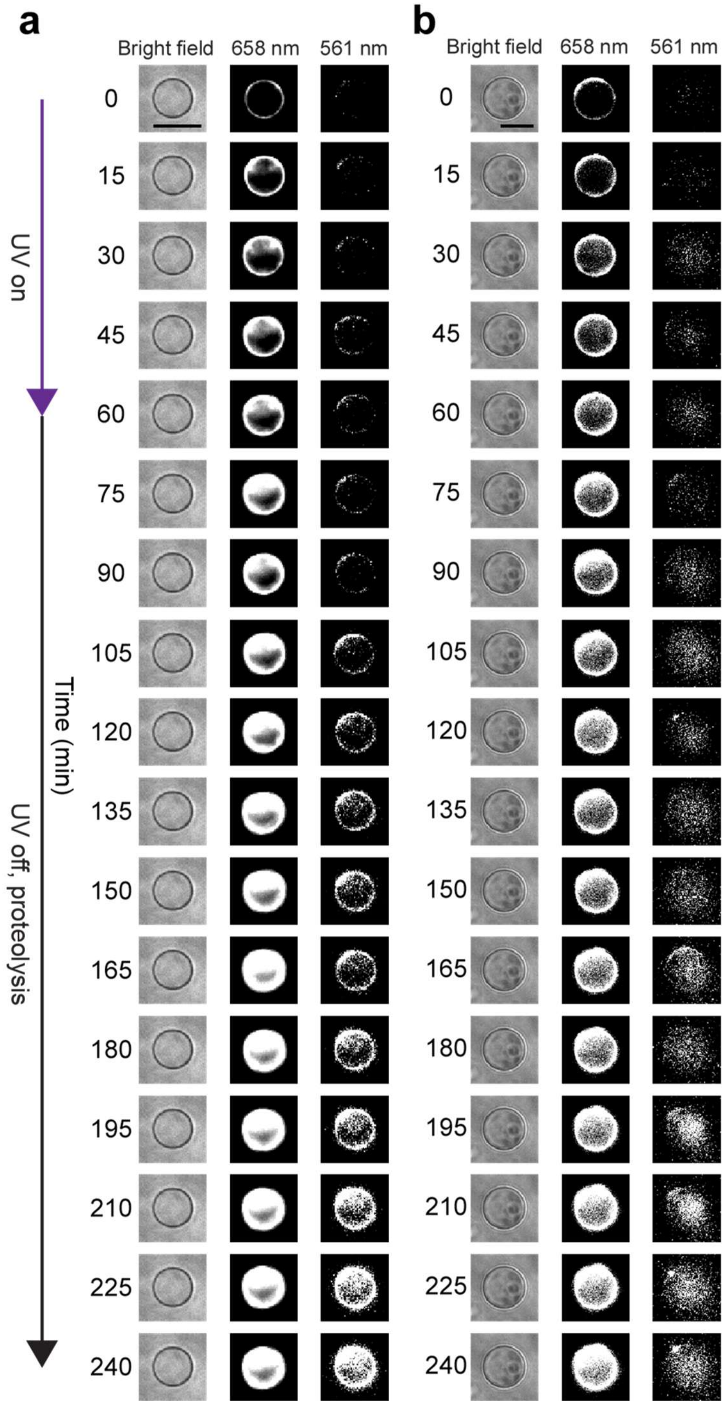


**Figure S40. Two experiments of Au-enzyme delivery and proteolytic activity within vesicles (substrate: BSA, enzyme: PK).** The experimental workflow follows Fig. S39a, using TMR-conjugated BSA as the fluorogenic substrate (instead of Bodipy-conjugated casein). **a** and **b,** Vesicle images under brightfield illumination, 658 nm excitation (Cy5-labeled Jitterbug fluorescence and Cy5.5-tagged Au-enzyme fluorescence), and 561 nm excitation (TMR fluorescence). Time-lapse imaging was performed at 15-minute intervals to monitor proteolytic activity as indicated by enhanced TMR fluorescence within vesicle interiors. Scale bars represent 10 μm.

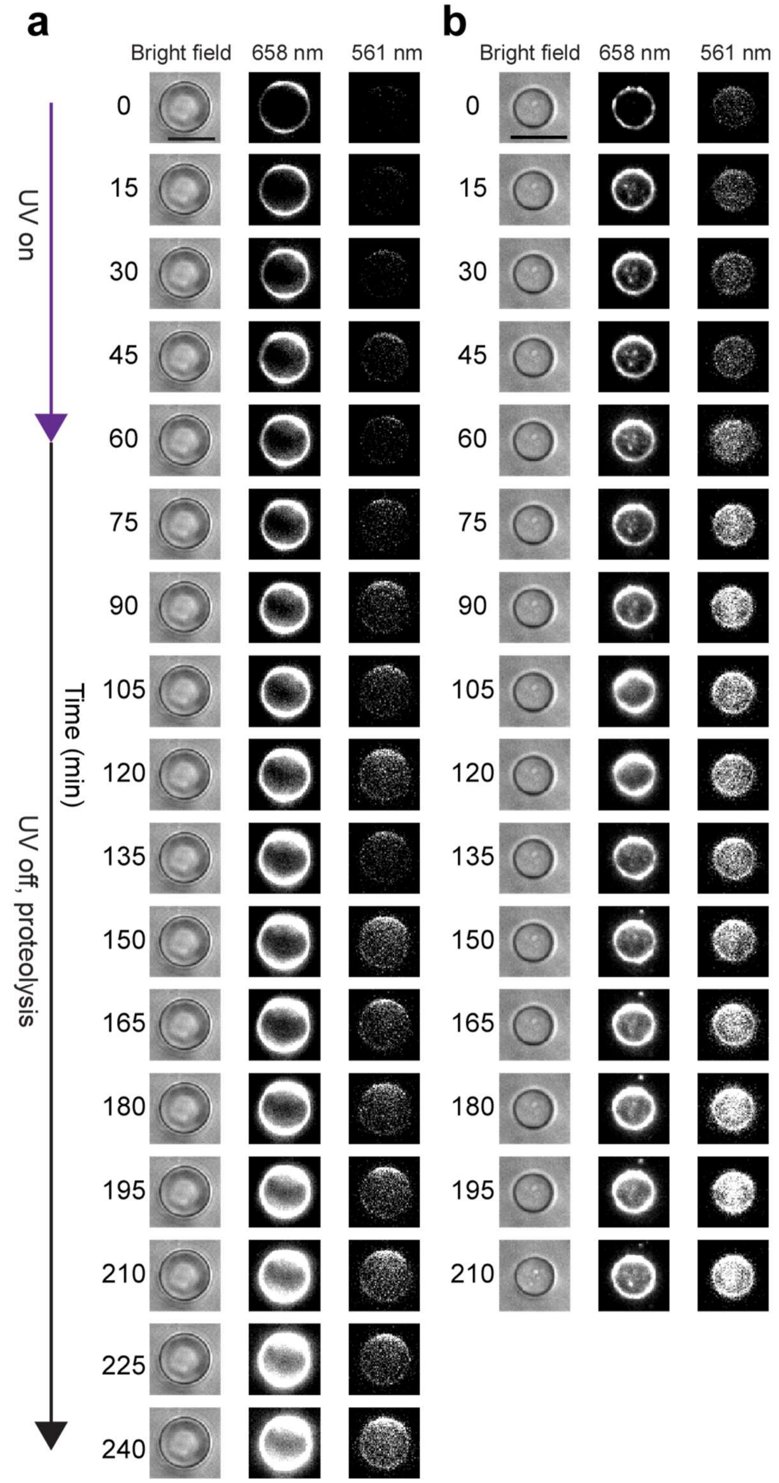


**Figure S41. Two experimental sets of Au-enzyme transport and proteolysis within vesicles (substrate: Casein, enzyme: trypsin).** Experimental procedure is given in Fig. S39a with different substrate-enzyme combinations. **a** and **b,** Microscopy images of vesicles under brightfield illumination, 658 nm excitation (Cy5-labeled Jitterbug fluorescence and Cy5.5-tagged Au-enzyme fluorescence), and 561 nm excitation (Bodipy fluorescence). Scale bars represent 10 μm.

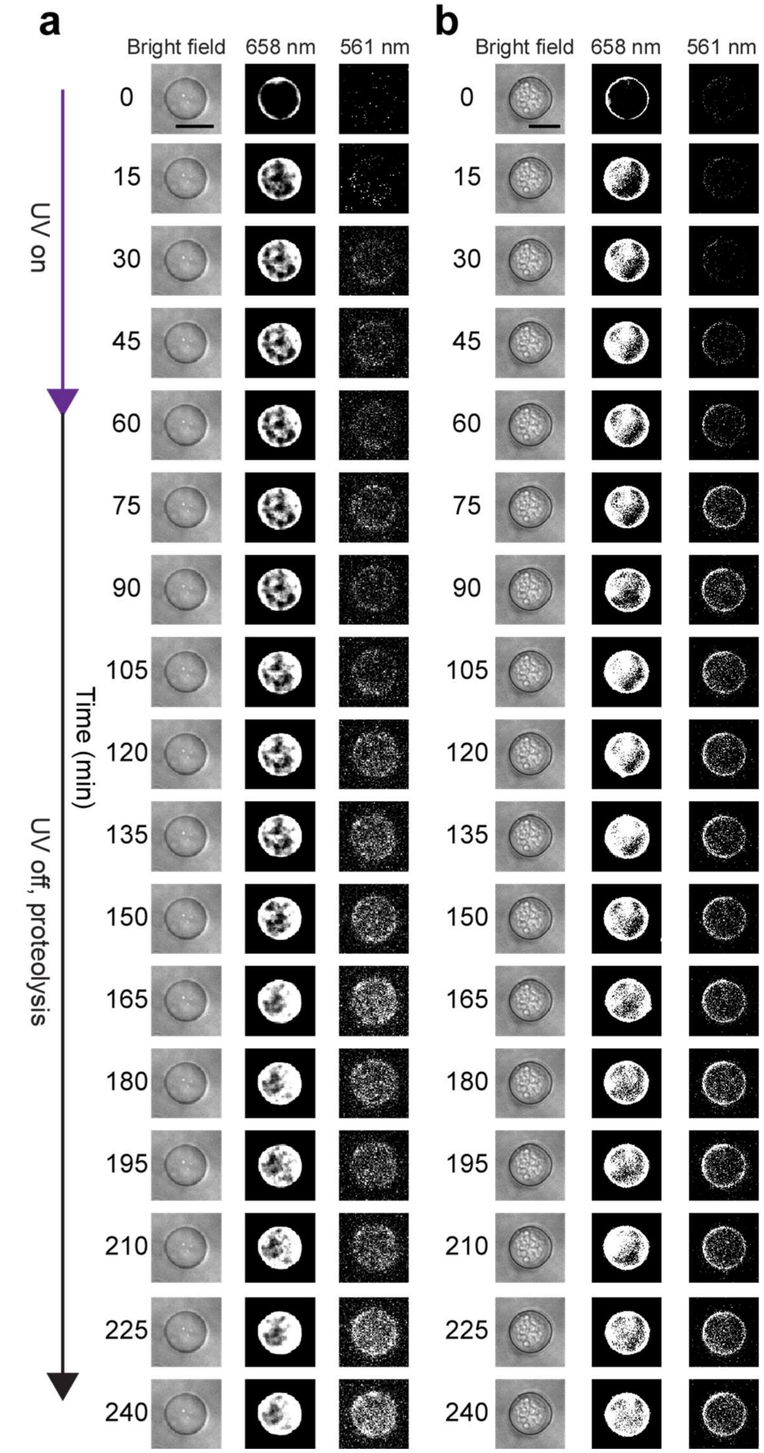


**Figure S42. Au-enzyme transport and proteolytic activity within giant vesicles (substrate: BSA, enzyme: trypsin). a** and **b,** Microscopy images of vesicles under brightfield illumination, 658 nm excitation (Cy5-labeled Jitterbug fluorescence and Cy5.5-tagged Au-enzyme fluorescence), and 561 nm excitation (TMR fluorescence). Scale bars represent 10 μm.

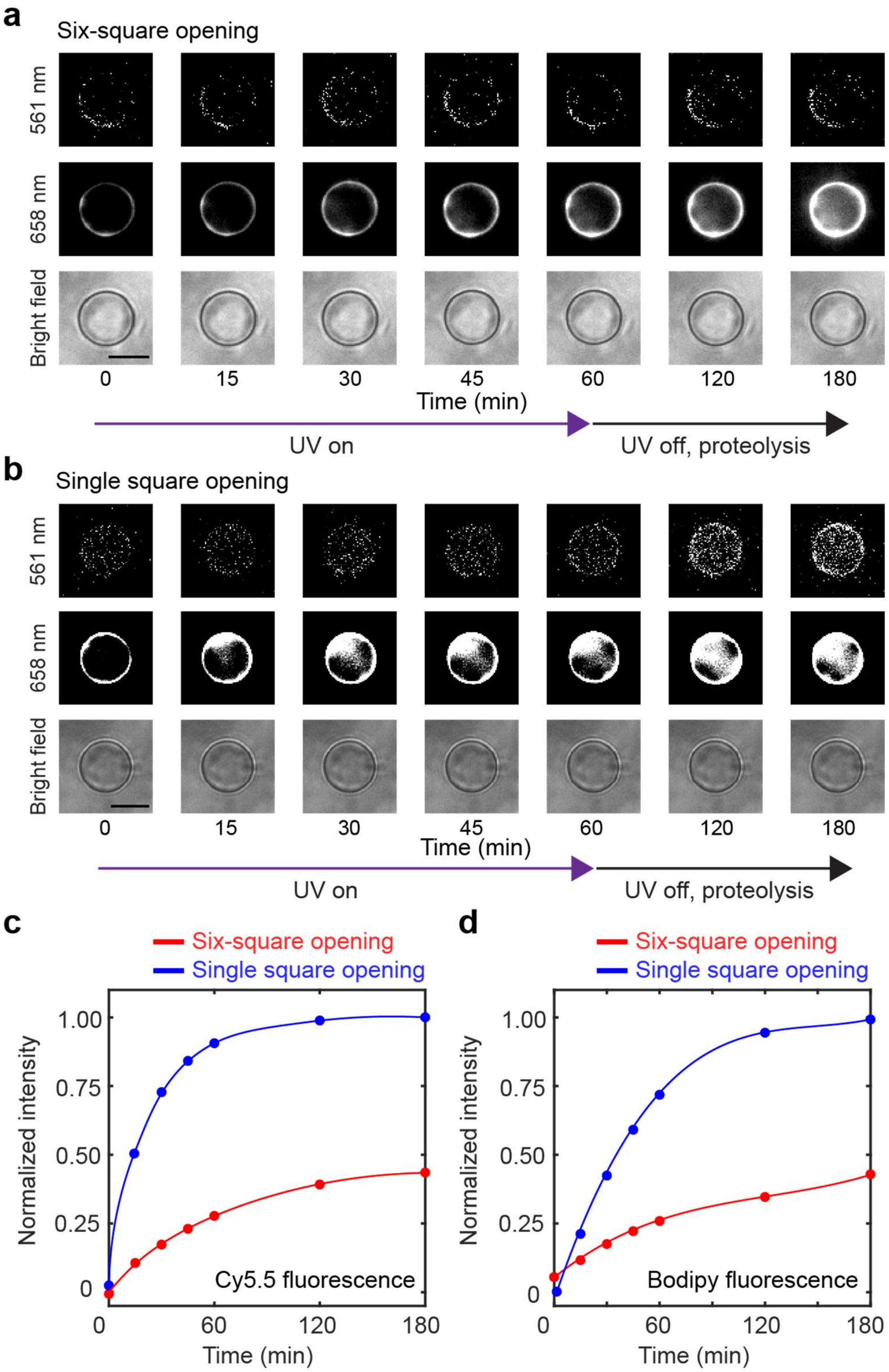


**Figure S43. Enzyme transport via single-square vs six-square opening in DNA transformers (substrate: Casein, enzyme: PK).** Enzyme delivery and proteolysis in vesicles by opening **a,** all six squares and **b,** only one square interfacing lipid membranes. **c,** Fluorescence intensity of Cy5.5-labeled Au–enzyme complexes (658 nm excitation) inside vesicles from **a** and **b**. **d,** Bodipy fluorescence intensity (561 nm excitation) inside vesicles for cases **a** and **b**. It is evident that opening a single square is more efficient in delivering Au-enzyme complexes than six-square opening. Scale bars in brightfield images represent 10 μm.

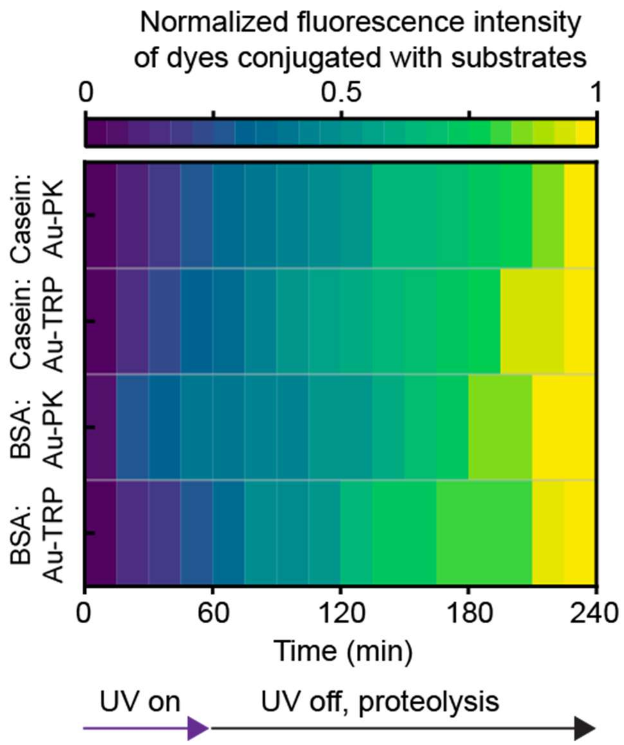


**Figure S44. Heatmap of normalized fluorescence intensities for several enzyme-substrate combinations.** The fluorescence intensity for each proteolytic reaction from Fig. 5b was normalized to the respective maximum value. The quenched dyes conjugated to the substrates became activated by Au-enzyme complexes, resulting in increased fluorescence intensity inside the vesicle over time.

#### S14. DNA Sequences

**Table S1.** Regular edge staples bound to M13mp18 scaffold (cy 3 ●).

| Name | Sequence (5'→ 3') |
|---|---|
| 11_1 | AACATGTCCTTTTGTATTTTCATCGTAGGTCAATCACCTTCA |
| 11_2 | CGACTCTAGAGGGCTTAATTGCTGAATATTGCGGACGAGAAC |
| 11_3 | GCTGTTTACGACGGCCAGTGCCAAGCTTTAATCATTATCATT |
| 11_4 ● | [cy 3] AGTACCGCACTCATTGGCTTAGAATCCCCGGGTACTAAACCA |
| 11_5 | AACCGAACCGTTTTATAAGAGGTCATTTTAATGCTCTTCCTG |
| 11_6 | AAGCAAGCTGACCAACCAGGCGCATAGGGTCTGGCGTAGCTC |
| 11_7 | ATGCAGACGGGTATCGAGCTCGAATTCGGCATGCCAACAACC |
| 11_8 | CCAAGAAACGCGCCTCTGTCCAGACGACAGCGAGTTGCAGGT |
| 11_9 | TCAAGAGACGCCATCAAAAATAATTCGCCTGGCTGATAAGGG |
| 11_10 | TAGCCAGTGTACAGACTTTGAAAGAGGACACATGTTCAGCTA |
| 11_11 | AAATGTGGACAATAAACAAGATGAACGGCTTTCATCAACATT |
| 11_12 | CGTCGGAAAGTAATTGTTTATCAACAATCTTTCCTGGTCATA |
| 12_1 | GGACTAAAAAAGTAGGCAACATATAAAAGGAATTAATTGTGT |
| 12_2 | TTTTAAGAGACTTTTTCATTTATCCGGTCTTACCGAAAGACA |
| 12_3 | AGGCGTTCAATGAATATTTTGTCACAATCATTAAAAACGAGG |
| 12_4 | CAAGAAATTAGCGAACCGCGCCCAATAGTAGCCGGGGTGAAT |
| 12_5 | ATGCCACAACGGAGGCACCATTACCATTCAAACGTAGAAAAT |
| 12_6 | ATAGAAGGCGAGGAAGTTTCCATTAAACCTGATAAGAGCCAG |
| 12_7 | CGCAGACAATCATTACCTCCCGACTTGCCCCAATAATAAGAG |
| 12_8 | CGAAATCCGCGACCACTTGAGCCATTTGGAAACGCAAGCCCT |
| 12_9 | ACATACAACCAGTAATTTGTATCATCGCGGGTAAAGCTTTGA |
| 12_10 | CAAAATCTAAAGGTAGCAGATAGCCGAATACAGAGATACGTA |
| 12_11 | CCACGGAGTCACCGTGCTCCATGTTACTCAAGCAAGAACGCG |
| 12_12 | TATCACCATAAGTTATAGCAATAGCTATATTCTAAATCAGAT |
| 13_1 | CTAAAACTACCAAGCGTGCTTTCCTCGTAAGGAGCGGGCGCT |
| 13_2 | GCGATTAGAAAGAGGGCAACAGCAATAGATTAGAGCACCAAC |
| 13_3 | CATGTAGATAATATGCCGATTAAAGGGAGAAAGCCGTCCACG |
| 13_4 | AAGAAAAAAACCAAGTATTGGGCGCCAGGCAAGCGGGCGAAC |
| 13_5 | TAGATAAAAATATCTTTAGGAGCACTAACAACTTGATTGCCC |
| 13_6 | TTCACCGTGAGACGGCAAAAGAATACACTAAAAATTTACGAG |

| | |
|---|---|
| 13_7 | TCACCAGCCTGGCCAGGGAAGAAAGCGATAGAATCACCCCCA |
| 13_8 | CTGGTTTGGTTTGCTCAATAATCGGCTGGATAAGTCAGGAAC |
| 13_9 | AGGGCGCCGTATAACGCGAAACAAAGTAACGAAGGCCGTCAA |
| 13_10 | GAGCTAAGAAAGGACTGAGAGAGTTGCAGGTGGTTTTTCTTT |
| 13_11 | GTGGCGAACAGGAGCCCATCCTACACTCATCTTTGAGAGCGG |
| 13_12 | GGTACGCCCCGATTTAGAGCTTGACGGGTTTTAGACCTGAAC |
| 22_8 | AGACTTTACAGTTGCCCGCCGCGGGCACAGACAATCAGTTGG |
| 22_1 | GTAAGAATACGTCTTAATGCGCCGCTACCACCACAAAAGGAA |
| 23_9 | CTGAAACCCTGCCTATTTCGGAAGTTGAAAATCTCTTTCGAG |
| 31_1 | ATTAATTGTATAAAGAAAAAGCCTGTTTAAAGCACAAATCGG |
| 31_2 | CTTACCAGCGTTGCTCGTGCCAGCTGCAGGTTCCGTAAATCG |
| 31_3 | ATCAAAATCATTTCACATCAAGAAAACAGAACCATTAGGGTT |
| 31_4 ● | [cy 3] GAATTATTTATTTGTTATACTTCTGAATCCCGAGACACCCAA |
| 31_5 | AACGCGCGCGAAAATCCTGTTTGATGGTTTAATGAAACTCAC |
| 31_6 | CAAAATCAAACCTGGCTCACTGCCCGCTTTAGAACCTACCAT |
| 31_7 | AGAATAGAATGGAAGGGTTCCAGTCGGGCCTTATAGGTCGAG |
| 31_8 | GAGTGTTGTTTGGACACGTAAAACAGAAAATCGCGTTACATT |
| 31_9 | GAACCCTATTACTAGCCAACGCTCAACAGTGAGCTATCGGCC |
| 31_10 | GTGCCGTAGTATCATATGCGAAGATGATTTTTTGGAATCAAA |
| 31_11 | ATCAAGTGAAACAAAATTACCTGAGCAAAAGTTATACAAATT |
| 31_12 | TAACAATCAGGGCGATGGCCCACTACGTAAATTAACAGAGGC |
| 33_1 | AGTCACGATTGTTATGGGGTGCCTAATGTAGGGCTAATATAA |
| 33_2 | GGGTAACCACACAACATACCGGAATCGTAAGGCGAGGGCGCA |
| 33_3 | GTGCTGCCATAAATATAGCGTCCAATACTGGAGCCGGAAGCA |
| 33_4 | AACAGTTCTCTTCGCTATTACGCCAGCTCCTCAAAATAGTAA |
| 33_5 | GAATCGCTAAAGCCTCCGCTCACAATTCGCCAGGGATAGGTC |
| 33_6 | TAAAGTGCATATTTTTTTCGAGCCAGTATAATGGGTTTTCCC |
| 33_7 | CAACACTAGACTGGATTCATTGAATCCCGGCGAAACTGCCAG |
| 33_8 | AATGTTTATCATAATACATAACGCCAAACGTGCATGGGGGAT |
| 33_9 | AGTACCGGAACAAACGGCGGATTGACCGATAAGAGTAATTGA |
| 33_10 | ACGTTGGAGAGGCAAACAACGCCAACATGTGCATAGTAAGAG |
| 33_11 | TCGTAACAGGAATTACGAGAATTTAGGCTGTAGATTTAAGTT |
| 33_12 | TTTGAGGATGCAGACCCTCGTTTACCAGGGGGGTATGCTTTA |

| | |
|---|---|
| 41_8 | AGATTAGCAGAGAGAAAGGGCGATCATTAAAGCCATTGAGCG |
| 42_8 | CGTTTTAGAGTAGAAACATCCAATTTGTTAAAATTCCAATTC |
| 42_1 | AAACGTTAATATTAAATCATACAGGCAATAGCATTTTTAGTT |
| 62_1 | AAGAAACCTGAGAGAACCACCAGCAGAAAGTGAATAACCTTG |
| 73_5 | CGCATAACTTGATACCGATAGTTGCGCCGACAATTTTTGTTT |
| 73_1 | TTAGCGGTTAACGGAAATAGCAGCCTTTAGCCATATTATTTA |
| 81_9 | CAAAAATGCAACTGAGCCAGCTTTCCGGGACCCTGTAATACT |

**Table S2.** Regular edge staples bound to p9072 scaffold (cy 5 ●, overhangs for cy 5 strands, overhangs for cholesterol moieties, overhangs for photocleavable strands, and **spacers**).

| Name | Sequence (5'→ 3') |
|---|---|
| 21_1 | GTCTCCCCTGTTAGCTCAAGTATGCTATAATGTAATTGAGGT |
| 21_2 | CTGTATTAACGCGCCTGGTCGTCGTGCGATGCACAATAGTAA |
| 21_3 | CTGCTAATCCACCTCAACATGTCTAGTGGCAGACTTACTAAA |
| 21_4 | AGGCCGTAATGTTTTAGCGGACGCATAATTTTCTCAAGCCAG |
| 21_5 | TCGTGGTATCAATTTAGGCTTGAGATAACTGGCCTGCTGAGG |
| 21_6 | CTCTGCATGAGCAGAGAGCAAAAATGTCCTTAGAACCGAGCGGGCTCTTGGGCAACG |
| 21_7 | ATCCTAACTCTGACGCCTGGCTATCGGGTGGGGAAGGAGCGA |
| 21_8 | CTCTGCATGAGCAGAACTGCGAAAGTCAACTAGAGGTGTAAGGCGGAAGGCCTTACG |
| 21_9 | CTCTGCATGAGCAGATTTATTTCGTAGGTTTACAACTGGTTATTAGGTTTCTAAGGT |
| 21_10 | TCTTACATTTGTAGTTGCCAGGACTTAACCATACTAACTTTG |
| 21_11 | TATTTAAATGCCCAAACAAAAAGATACTGCGTTGAGAGAGAA |
| 21_12 | CTCTGCATGAGCAGAGTGATCAAGTCGTATGGCTAAGTTTCCCCAGTTACTACCCCT |
| 32_9 | TACTTCCCTGTCCATAGGGAAAATTCCCTTTAACGTAAAAAG |
| 43_1 | CGGTTACAGATAGGTGTCAGAAGTAAGTACTGATCTAAGGGC**TT**CGATGGCCCT |
| 43_2 | CCTCGCCTGCACTCGACTACCATCTTCACAGTTTGCTCCTTC |
| 43_3 | AAAACCGGCGAAGGCTTATACAGAGCAAACGTCAAGCGAGTTTTTCTGAACCGC |
| 43_4 | AGTCCACTATTAAAGTTGTGCAAAAAAGCGGTTAGGAACAAG |
| 43_5 | TCTTTTACGATCGTGTTGAGTGTTGTTCGGCCCGTCTGGGCT |
| 43_6 | GGTCCTCCTTTCACAAAATGCCGCAAAACCGAACATGTACTG |
| 43_7 | CGCAACGCCCCCATGAACGTGGACTCCATACATAATTTGAAA |
| 43_8 | ACATGATTTGTTGCGGGAAGCTAGAGTACTACGTTATAAGTA |
| 43_9 | GACACGGGGACGTACCGTAGAACGACGCAAGGGAATTCAGCATTTCTGAACCGC |
| 43_10 | AGAGCCTGGAAGGCCAGCGTTTCTGGGTGAGTTAATAGTTTG |

| | |
|---|---|
| 43_11 | AGTAGTTCGCCAGCAAAAACACTTTGGTGTTGGTGGGTGCGC |
| 43_12 | TTGGGTCTGTTGCCCATTGCTACAGGCAGATCAAGAGGGCGA |
| 51_1 | TAAATTGCATCGTAACAAGGAGTCTCAGTGGTAGCTCTTGGT**TT**<u>CGATGGCCCT</u> |
| 51_2 | AGCGCCAACAGGTGATTAGTAACACATGGCCCACGGTCAATG |
| 51_3 | TAGTCCCTCCGCCTGCGTATGAAAGTAGAAAAAGAGGGGCAG**TT**<u>CGATGGCCCT</u> |
| 51_4 | <u>CTCTGCATGAGCAGA</u>TTTCGAACGGGTGCAGAGCGAGATCCACTCGGATTGTTGGTA |
| 51_5 | <u>CTCTGCATGAGCAGA</u>TCCAGCACGTCGCTCAGCAGATTACGCGTTGTTGATGGAGCT |
| 51_6 | GCTGAAAGGACGTATATTAGGCGATTCGACGACTAGGTGGTT |
| 51_7 | <u>CTCTGCATGAGCAGA</u>AAGGAGGCTCCGAATTGGCTGCTCCACCTGCCCTCACAGTCG |
| 51_8 | CATTGCTATCAAGCAAACAAACCACCGCATACGTGCGGGGGA |
| 51_9 | <u>CTCTGCATGAGCAGA</u>GGAGTAGTTGCAAGAAAGATGAATCTAGGTGAATGCCAGGAT |
| 51_10 | TTTTTGTTCAGTTTATGCATAGGGCTACTAGTATACAGGTAA<u>TTTCTGAACCGC</u> |
| 51_11 | AAGTCTGATCCGGCGCATCTCAGTCATGCGACCTTACCTTCG |
| 51_12 | GCTCTTGTGCGGTGCAGTAAATTTCTGCCGCGGAACTCTTCG |
| 52_1 | GCAATAATTTCCTCGTTAGCGCGAAGACACGGGTAAACAACC |
| 52_2 | CACCCCGACAGGAAGACGTCCCAAAATATAGTCTCGTGCGTC |
| 52_3 | GTTGAAGAACAGGGGCTTTTAGAAATACGATTAAGCTGTCCC |
| 52_4 | ACTCCATTTTGAGCTCTGGACTTGACACTCGCGGCGGGTCCT |
| 52_5 | ACTTCGGGACAACCTCGAAGAACCCGGCCCATAATGGAACCT |
| 52_6 | CCCCAGGTCGCTTATGTTGGTGGACATCGGACGGCCATTAGG |
| 52_7 | CACGAAGGAGTGGTCCGCCCAATATCATCACTCAGGAGATTC |
| 52_8 | AGCCGTATGTTCTTAGTGGAGAAGAATTACCGGAGCTACAGT**TT**<u>CGATGGCCCT</u> |
| 52_9 | CAGCGTCCAGTCCCTGCAAATCTAAGTCATAGAGCTAAGAAT |
| 52_10 | GCCTAGACAAGCAGATGAGTCTAGCATTACAGCCTCGTCGGA |
| 52_11 | CACCGCTCCGGCAGAGAGATATCAACTGATTCGACACAAGCC |
| 52_12 | TGGGTAAATGGAAAGTGGATGTCGAGCACATAAGTCCTCCGC |
| 53_1 | TGGCGGACCTGTTCCATGAGGAGTATGAAGCATGCCCTAAAC |
| 53_2 | TGATCTAAAATGTCAGTCGCGCAAACGGACCTTTTTACTTCT |
| 53_3 | AGTCGATTCGCTTCTGATCCTGATTGAACACCGGGTTTTGCG |
| 53_4 | CGTGTCCTTATCATGTGAATATAAGAGATTAGGTACGCACTG |
| 53_5 | CGACCGGATTGTAGTAGTCATAGTGTCCACTCTACCCCTGCC |
| 53_6 | TTTTAACGCAATAGCCATGCCGTGGGACTAAGAAAGAGGGGT |
| 53_7 | AACGACACACGGGCGAACGGTATTATAGTACTGTTAACAAAG |

| | |
|---|---|
| 53_8 | AAATCAGCTGCTTTCCGGGGTCTCGTGCCGTGCTTCTTACGT |
| 53_9 | GCGATGTAGATAGAGTACATCAAACAATATAATAAGAGCCAA |
| 53_10 | AGGTTAGCATTTGGATTGAGTCACCACTGCATATAGCTACTC |
| 53_11 | AAGAACTGCTCGAATCTTGATCCCTCAGTGACGGAGCACCAT |
| 53_12 | ATGCCTCTTAGGCAGCGGGGCAACATTATCTCCCCGGATGCT |
| 61_1 | ATTGTCTGAGAGCATCCTTAAGAGTTTTGCCGGGCGGCACGA |
| 61_2 | CACCCCTATGGTCTTCACTCATGACACGAACATCCCGCTCTG |
| 61_3 | ACAGGACAAATTTTGTGACATTGGTCTCAACACTAGGGCTTATTTCTGAACCGC |
| 61_4 | ACTATTCGTTATGTCGTTCGTAAATCTCCCATGGTCCTGGGA |
| 61_5 | CTTATCGCTCGGTAGTTCTGAGTTACGGTTGACTATCACCTC |
| 61_6 | TAATGACCTAAGTGTGGCGTGTCTGTCGCATGCGTGCAGAAC |
| 61_7 | AAGACCTCAGACGAGCGCCCTATAGGACGTAGCAGGTGACTA |
| 61_8 | CATACTGTTCAGTTGTCGGTACCAAGCTCCATTCTTAATTAA |
| 61_9 | GCATATCGCGCCTCTGGATCGGCTTGGGCTAGTTGTGCTTGCTTTCTGAACCGC |
| 61_10 | ATCCGGGGGAGACTTTTATGTACCTACAAGATCGTCAAGGTG |
| 61_11 | TAAGTTCGATCCTCAAATCCTTTTCTACTCGCAACACGGACC |
| 61_12 | ACGTCGTACGAATATCCCTACACGGCTATCGGCCCGCTATGG |
| 63_1 ● | [cy 5] AAGACTAATCTCGACGTTAACTGAGCACCGGGTCGCTGCATC |
| 63_2 | GCCTGACGTGTACAGATATCCTCTATGTTTGTTATTACGCTG |
| 63_3 | GAGTGTGTGAGTGGAAACAAGTGGGTTCATTCATAGGCTAATTTTCTGAACCGC |
| 63_4 | GCAGGAGCGCAAGTAACTCTCATGTACATGATTACCTTCGAG |
| 63_5 | CGCTATCCCGTGACGGTCCATCTGATGTTCGCAACAAACCAC**TT**CGATGGCCCT |
| 63_6 | CTCTGCATGAGCAGATCCGCGAAGAGGTCATACAGCTTCCGAGTATTTGTTAGGTAA |
| 63_7 | GCAAATGCGCACGGGTGAGTCCGTTGGTATTCCAAGCTGGGG |
| 63_8 | CTCTGCATGAGCAGATGCCTGCGCAGTCCAAGGATCGAGCTTACCTTGGAAGACGAA |
| 63_9 | CTCTGCATGAGCAGATAACGATCATGCGGATAACTTGCAGGGCGTTCTCCCGCCGAC |
| 63_10 | GCCTCCATTTAGGCCTGCAATTGGTGAAAGTAATAAGAGGTT |
| 63_11 | GGCTGTTCGAGTCAGGTGCAGCGAGCTCTTGGTTATGGAGAG |
| 63_12 | CTCTGCATGAGCAGACGACGAGTCCTGTCGACGCTCACTATCCAAGTGCCGTGCCAG |
| 71_1 | AAATTCGTCTCATGACTCTTCCTTTTTCGCAGCCGTATTGTT |
| 71_2 | GTTATTGCGTTAAAAGGCCGAAATCGGCTCGCCCCGAATCTG |
| 71_3 | ATTCCGAAAGGATCTTTTAAATCAATCTTTAGCTAAGAATGC |
| 71_4 | TATCAAACCACTGGTCTTGCGTGTACGTGGATACACGATTCA**TT**CGATGGCCCT |

| | |
|---|---|
| 71_5 | ATCAAAAAAATACGAAAACGGATCTAGGAAAATCCTTTGTTA |
| 71_6 | AAGACGTAACCAATTTTTTGTTAAATCAGCGCTGATACGCGA |
| 71_7 | ATGAGGGAGACCCGGACAATCATTTTTTCACAGGGTAATGCC |
| 71_8 | TCCTCCAAAATCTGAGGGTAGCGCAGGTTTTGGTCATATGAG |
| 71_9 | GTAGCTGTACTCATAGCGGATACATATTTTAATATCTTATAA |
| 71_10 | ATGATGGAATATTATTGAATAAATTAAATAAACTATCTTTCT |
| 71_11 | CTCTTCAAATGAAGTTCACCTAGATCCTTTGCATTTATCAGG |
| 71_12 | TAAACTTTATGAACGTTACTGCCTTGAAAAAGTATATGAGAT |
| 72_1 | CTCTGCATGAGCAGATCTTAAAAAGGCCATGACGAGCATCACACGACCCTATCTCAG |
| 72_2 | GTGAGGCGCCGCGTTGCTGCCCTTGGTGCGGCGCCCGCTAAG |
| 72_3 | GAAGGCACATACCAGATCGCTAGCCACCTCGCGTTTTTCCAT |
| 72_4 | CTCTGCATGAGCAGATAGACCACGGCGGAGACAAGGTATTCCTGGCTCATCCGGTGA |
| 72_5 | ACCGGATGCCCCCCGGAACCGTAAAAAGTTCTATAGCTACGG |
| 72_6 | AGGCTCCACCTGTCCTCATAGCTCACGCAGTAGACCGGGAGA |
| 72_7 | CAGTGGACGGCTCGAAATTAATCATCGTAAATGGGTGATCGG |
| 72_8 | CGCAGCTACGAAAAAAGAAGATCCTTTGATAACCGAGGGCGT |
| 72_9 | CTCTGCATGAGCAGATTCGGTGCTTTCACCCTTCTTTCCGTTATGTAGGTGCCGCTT |
| 72_10 | CCCAGCAGCGCTTTCGCCTTTCTCCCTTCGGGGTCTGACGCT |
| 72_11 | ACTCGCGATCTTTTCTACGGGAAGCGTGTTGGCTACCCTAAC |
| 72_12 | CTCTGCATGAGCAGAATAGAACGGATCTCCTCACGTTAAGGGATTTCCAAGGGACTT |
| 82_1 | AATAACGACCAGTAAAGGCCAATTATGACTATCCACCAGTCA |
| 82_2 | ATATATCATCTCCTGGCGCCAGCTGCAGTCCAATCCCGCAAC**TT**CGATGGCCCT |
| 82_3 | TTTTCGGACATAGGCCACAATTTGCCCAACCTTAGCTTAGACTTTCTGAACCGC |
| 82_4 | GCTCGCCGAAGCAACCGCCTTTTAAGCTGGGTTATGATTCGT |
| 82_5 | GTCATAAATGTAACAGCACACCTACCCCTGTTCGTTGATCCG**TT**CGATGGCCCT |
| 82_6 | TTACAGGCTCGCAGTCGGTATCGCCGTATATGGTGGCTATAC |
| 82_7 | GACGTGTGCGTCGGTTGTAGAAGGCGACGGCGTCCGCAGATG |
| 82_8 | CGGGAGCATACGATCCACCGGCTACATCTTCCTACAAAGTCA |
| 82_9 | GCAGTAGAGACTTGCTCGGGGATCCAAACGTGTTCTACCGGCTTTCTGAACCGC |
| 82_10 | CAAGCGGATAAGCAGAAAGGGTCAGCGGAACTAGTTCACACT |
| 82_11 | GAAACTGTGTGTCCAGGTGCGTATTAGATTCCTTGACGTGCA |
| 82_12 | CAATCTGGTTCATCACATGTGCCGAGGGAGCAACTACTCATC |
| 83_1 | CGGAGCCAGTAAGTGTGAATTGCCGGCCCTATTGGTACGACG |

| | |
|---|---|
| 83_2 | TATAACTAGATCTACAATCCTACCTATTACACGCTATTGTCC |
| 83_3 | GTCGAGGCCGCCGCCACCGTCTCAAGACATTTGAGCCAAAGCTTTCTGAACCGC |
| 83_4 | ACCACACTGCCGTACCACTACGTGAACCACCGAAGTTTAGAT |
| 83_5 | CGTCTGCCCTATGATTCTAGGGAGGCACTCCGGGATCAGAAG |
| 83_6 | GATGGCGGGTTCGTCGCCCCGTAGGGCATCAAGTTTTTTGGG |
| 83_7 | AGGCCTTATCACCCTAATCCTAGTATCCTTTCTGAGACTAGC |
| 83_8 | GCTGCTTCGATGGCAAGCACTAAATCGGCGCTGCGAGAGCAG |
| 83_9 | AAGCGTTGGCGGCCATTGTGTGGGGCCAACTCTCGTTATCCGTTTCTGAACCGC |
| 83_10 | TATCTAAGAAACAAATTCTCTCTAGAGCGAAGTCCAGTGATG |
| 83_11 | CGGGCCAGATCGCCGCTTAATGTTCGCTAGCCGGGCGAGTAT**TT**CGATGGCCCT |
| 83_12 | TTCCGGTCGAATGACAGAAGATCCCGCGGGAATGTCGTAACC**TT**CGATGGCCCT |

**Table S3.** Regular edge staples connecting the two scaffolds (cy 5 ●).

| Name | Sequence (5'→ 3') |
|---|---|
| 22_1 | CGGTCAATGCATCAAGAGATAGAACCCTATTAGTCAACCCTC |
| 22_2 | TCGAACAGACGCAACCTTTGCCCGAACGAATATCATTTAATG |
| 22_3 | AGCGAACAAACCAGGGGATGCGACCATCTGTAATGATTCGAC |
| 22_4 | AGTTTGAGCATCACCTTGCTGAACCTCATTATTAACTTGTAC |
| 22_5 | AATCAATATTAAATCCCGCGCGAGACAGCATTTCATGAAAGC |
| 22_6 | AACTCGTATCTGGTATTTTTGAATGGCTTCTGACCCAAGTAC |
| 22_7 | CAAATCAACAAACATGTGAGAGGCCCAAGTTCCATAAGTATT |
| 22_9 | TTGAGGAGATTTAGATGCTGAACCTACTACGCGTTGTACTAT |
| 22_10 | CGCGAACGGCCAACTCTGGGAGTTTCTCTACACAATTTTAAA |
| 22_12 | GGTTGCTAGCGGTCACGCTGCGCGTAACAGGGCGCACCGCAT |
| 23_1 | GTGAATTCGGTCGCGGAAAACATGAAAACCGCTTTATAGCCT |
| 23_2 | ACCCAAAAGTTAATTGTATCGGTTTATCAGTTAAAGAATTAT |
| 23_3 | ACGAGGGAGAAGGAAACCGAGGAAACGCAAGACAGGTGACCT |
| 23_4 | TTTGCGGGATCGTCAGCTAGGCGGCTAGCGCATTGAAGGAAC |
| 23_5 | GAGTAAAGTTTGAATGAGGCTTGCAGGGAGCTTGCCAAAAAA |
| 23_6 | ATCGCACGGTTCGCCGAATAATAATTTTTTCACCCTATTATT |
| 23_7 | GCCGGGAACTTCTCACCCTCAGCAGCGAAATAATAACTCCTT |
| 23_8 | ATATAACGTCATCGTTTCAGCGGAGTGAGATTAAGACGGAAT |
| 23_10 | AAGGCTCGGAATTGGACGATTACATGCCGTGCGAGGGCCGCT |
| 23_11 | AACTAAACAAAAGGAGCCTAACTGGCATGAATAGAAAACCGC |

| | |
|---|---|
| 23_12 | ATTACGCTCAACAGACAAGTCAGAGGGGTTTTCAACATCGGA |
| 32_1 | AGTTACATAAAGAACTCAATTAGAGTGTAAAAGTTTAAAGTT |
| 32_2 | CCTAGAGATCACTACGGATTCGCCTGATCAGGTTTTTCGTTA |
| 32_3 | GATGAATACAATAACTAGCCTTGGCCGGTGACCTGAAGGGGC |
| 32_4 | GGGAGAAATACAGTACTTGGCATGATTGATCGCTGCTAGGAC |
| 32_5 | TTGCCAGCGACGATAAAAATTGCGCCATCATCGATTGATGCG |
| 32_6 | CGGACCTCCGCGTTATTGCGTAGATTTTTGCTTTGCGGATTT |
| 32_7 | ATGGAAGCGGGTCCCCTGCCGCGGTATATAATATCAACGTCA |
| 32_8 | CTACCAATGAAACTAACAGTACCGGCTTTTGCAAACGTTCTG |
| 32_10 | AGAGAATGTTGTAGTAACTTCCGCAGCATTGGGTAAATACCA |
| 32_11 | TAATGGTGAGCCCTTTCGGATGAGATACAAACCTCAGAAGTT |
| 32_12 | TGGATACCCTTCGGAAAAACCAAAATAGCGAGATTTTACATC |
| 41_1 | TGCACCCGGCCGCAAACGAGCGTCTTTCCATTAGACGGGAGA |
| 41_2 | TGCATAACCGCTGTTTCACAAACAAATAAATCCCATTCAACC |
| 41_3 | GATCTTATTCTCTTTTTGCACCCAGCTAAGGGTAAGAATGGA |
| 41_4 | CTGTGACATTGGAAAGGTAAATATTGACTTCATATGGTTTAC |
| 41_5 | ATTAACTCAACGCTGTGTTATCACTCATATGTAACGTCAGAC |
| 41_6 | ATCTTACGAACACCAATTTACCGTTCCAGAGGCAGCCACTCG |
| 41_7 | CTAATATTTGCTATACTGTCATGCCATCCGAAAACTCTCAAG |
| 41_9 | GATTGGCGTCTCTGCTGAACAAAGTCAGCAATTTTGCAGCAC |
| 41_10 | AAGCGCACTTGATATGAGATCCAGTTCGGGTTATGATCCTGA |
| 41_11 | GATTGAGAAAGACAATAACCCACAAGAAGAAGCCTTGCTTTT |
| 41_12 | CAGCGCCGGAGGGAAACGTTCTTCGGGGCGTAAGATAAATCA |
| 42_1 | CAAGTCACAGAACTAGATTGTATAAGCAGTTAAATATTCCAT |
| 42_2 | CACATAGTTCTGAGAAGCAAACTCCAACAAGTTTCCAGCTCA |
| 42_3 | CGTCGTTTGGTAGCCCGGCGTCAATACGCGACCGACTTCAAA |
| 42_4 | GAGTACCCAACTAAAGTACGGTGTCTGGAGGTCAGACTCAAC |
| 42_5 | ATAACAGAGACCGGAATAGTGTATGCGGGGATAATAAATTGT |
| 42_6 | GCGAACCTTGATTCCGCATTAAATTTTTAATATTTACCGCGC |
| 42_7 | TGCGAACATTCGAGGTTGCTCTTTGGCTTCATTCAAATATCG |
| 42_9 | TGACCATGACTTCAGCTCCGGTTCCCAACGTGGTGAATTAGC |
| 42_10 | TTTTTTAAACAGGATTAAAAGTGCTCATGGTGAGTGATTAGA |
| 42_12 | AAAATTACATCAATTCTACTAATAGTAGGGCAAAGTCACGCT |

| | |
|---|---|
| 62_2 | TGCCACGCACCAGACGGGCGGCATTGCTCCCCGCTCGGAACA |
| 62_3 | TTATCAGAGAACGTGGACTAGTATTAACCATCATAGACCGAG |
| 62_4 | CTATTAAATGATGGTCGATAGTGACTCTATTAATATTTTAAT |
| 62_5 | TAAACGAACGCTAGCGCTATTAATTAATAAAACATCGCCATT |
| 62_6 ● | [cy 5] AAACTAAAAGAGGAAGGAGCGGAATTATACCGCCTCAGAGGT |
| 62_7 | TGATAGGTAGGAGGAATCAATATATGTGGATAAAAGCAACAG |
| 62_8 | CCCTGCGTGTACGACAATTCATCAATATGGAACAAGAGTCCA |
| 62_9 | GGAAACAGTACATATAAGCTTCGCATACGTGATAATTCCTGA |
| 62_10 | AAAAATAAAATCGTTGGCATCTATAGCTCGTTGGAGATGAGC |
| 62_11 | CTTCTGTCCGAACGCCAGCAGCAAATGACATTTTGAATAAGA |
| 62_12 | GAGGCGGTCCCAACGTCAAAGGGCGAAATTACCTTCACTCGT |
| 73_1 | GTCAGAGTCCCTCGCTGAGACTCCTCAAGTAACAGTGCCCGT |
| 73_2 | TGGAAGCGTGGCGAAGAAACGATTGACAACAACCACGCTCAA |
| 73_3 | CCCGAAACAAATAGCTCAGTACCAGGCGGGAGTGTGAATAAC |
| 73_4 | AAATAAAAGTGCCAGTTACAAAATAAACACAGAGAACTGGTA |
| 73_6 | AACGTCACCAAATAAACCCGACAGGACTATAAAGCACATTTC |
| 73_7 | TCCCAATAAAATGAGGTCAGTGCCTTGAGAGAAGGTTCCCCC |
| 73_8 | ATAAAAATTTGCCACCTAAATTGTAAGCGAATGTAGCCGTCG |
| 73_9 | ATAAACATAAGAGGTGCGCTCTCCTGTTAAATCGATCGCCCA |
| 73_11 | ATAAGTTGGTTTTGGGGTTCCGCGATACCAGGCGTATTAGGA |
| 73_12 | AGAGGGTTACATGGCTTTTGATGATACAGATAAGTTTTAGAA |
| 81_1 | AGCGGTCACCCTAAAAAAGATTAAGAGGTCGCAAATGGTCAA |
| 81_2 | GCTAAATCGGTTGTGAGCGGGCGCTAGGTTAGAGCGAAGCAA |
| 81_3 | GGAAAGCGCGAAAGACCAAAAACATTATCACCGCTGCCATTC |
| 81_4 | GAAGAAACGGCGAATTACCCTGACTATTGCCATTCTCTGGTG |
| 81_5 | CTGGGCAGTACGCACTTTATTTCAACGCAGTATCGGCCTCAG |
| 81_6 | TAACCTGTGCATCAAGGGAGCCCCCGATGCGCTGGGCATAAA |
| 81_7 | AGCGGATTTTAGCTATATTAGGCAAAGCATAGTCATTGACGG |
| 81_8 | AGGCTGCCAGGTCTCGTGGCGAGAAAAGCATTAGACATAAAT |
| 81_10 | CCGGAAACCTTCATTTGGGGCGCGAGCTCCTCAGACAAGTGT |
| 81_11 | TTTGCGGGCACTCCTTGGGAAGGGCGATGAATGACTGGCTCT |
| 81_12 | GAAGATCGAGAAGCTTCCATCCTCAAAGTATCGAAAGGAAGG |

**Table S4.** Short jack staples.

| Name | Sequence |
|---|---|
| 13j22.1 | TTGACCGAGGGACATTCTCAGAATCCTGAGAAGTGTTTTCAGTAATAAAA |
| 13j22.2 | GACCATTGAGGCCACCGACAGATTCACCAGTCACACGACTATAATCAGTG |
| 13j22.3 | TGCGCATATTTACATTGGGTAAAAGAGTCTGTCCATCACGAAATGGATTA |
| 62j31.1 | GTTGACCACGGAATCATATTTCCCTTAGAATCCTTGAAAAGAATAAACAC |
| 62j31.2 | GTAATCCGGCTTAGATTATGATAAATAAGGCGTTAAATAACATAGCGATA |
| 62j31.3 | ACTGCGATACCGACCGTGAGACGCTGAGAAGAGTCAATATGGTTTGAAAT |
| 11j33.1 | ACTGCTGACATTCAACTATAATCTTGACAAGAACCGGATTAGGAATACCA |
| 11j33.2 | GATCTGACAAATCAACGTAAAGATTCATCAGTTGAGATTATTCATTACCC |
| 11j33.3 | GCATGCTATTACAGGTAGAACAAAGCTGCTCATTCAGTGGAACAACATTA |
| 73j23.1 | TTCGCAGATAAACAACTTTGATATAAGTATAGCCCGGAATGGGATTTTGC |
| 73j23.2 | GACCGATTCCGTACTCAGTTAGTAAATGAATTTTCTGTATAGGTGTATCA |
| 73j23.3 | GCTGCTAATTTCCAGACGGAGGTTTAGTACCGCCACCCTGTTTTGTCGTC |
| 81j42.1 | CATGTCAGAAGCCCCAAAAAGGATAAAAATTTTTAGAACGATAATCAGAA |
| 81j42.2 | AGGTCTACTTAAATGCAATCAATCATATGTACCCCGGTTCCTCATATATT |
| 81j42.3 | ACTACTGGAACTAGCATGTGCCTGAGTAATGTGTAGGTAGGTAATCGTAA |
| 12j41.1 | ATCTGCAGACAGGAGGTTAGCAAGGCCGGAAACGTCACCCGCCAGCATTG |
| 12j41.2 | CATGACTGCGATAGCAGCCAGAACCACCACCAGAGCCGCAATGAAACCAT |
| 12j41.3 | GATTGACCGAGCCGCCACACCGTAATCAGTAGCGACAGACACCACCCTCA |
| 63j21.1 | GTGACCATACCCTACTGTGCAATGTACGGGCACCAGCCGAGGTTGACGCG |
| 63j21.2 | GCTGCATAACTGAAGCTCTCTTCTAGCTTTTTCTTGCACCTGTTAACCTC |
| 63j21.3 | GCTCTGAAAGAATTCAGCAGCAATCGTTGAACCGTACTTCTGTAATTATC |
| 61j53.1 | GGCCATATGTATAAAATCCATCTACTGGGTGTTAGCATCCCCGATGAAGC |
| 61j53.2 | TGCGACTAGAATAGACCTAGCAATTGTCGGGCAGTGATTTTACTCTGAGC |
| 61j53.3 | TGTCAACGTTAACACGAAGATCTGCCTCAAGGATAAAGGCGTCGAAATTA |
| 72j51.1 | AGCTGTCATTACCTTCGGTAGGTCGTTCGCTCCAAGCTGGCTGAAGCCAG |
| 72j51.2 | TTAGGCACCGAACCCCCCAGTATTTGGTATCTGCGCTCTGGCTGTGTGCA |
| 72j51.3 | CGCGATTACTAGAAGAACGTTCAGCCCGACCGCTGCGCCCTACGGCTACA |
| 82j32.1 | CTCTAGAGCACCAGAAAGTAGCTCCCACATTTCCCATAGGCAGCCTGTAT |
| 82j32.2 | TCAGACTGTTACTCGGGAGATCATGAACTCTAGTACGTTGATCCGGAAGA |
| 82j32.3 | CAACTGTGACCAGGGCAATGGGATTGCCATTGTGCCATCACACGGTGGTG |
| 71j43.1 | GCTACTAGGTCTATTAATGGTCTGACAGTTACCAATGCTGCCTCCATCCA |

| | |
|---|---|
| 71j43.2 | CGCGATTAGCACCTATCTAGTGGTCCTGCAACTTTATCCTAATCAGTGAG |
| 71j43.3 | AGCTGTCACGAGCGCAGACAGCGATCTGTCTATTTCGTTGCCGGAAGGGC |
| 83j52.1 | ATGACTGCGCTGGTCTGACCGCGGTGCAACTCACATAGCATCAACGCAGT |
| 83j52.2 | CAGGATTCCCAAAGGCCCCAGCGGTAGCACATACAAACGTGTGGACGATC |
| 83j52.3 | AACCGTTGGGATTCCAGGTTGATGATCCAGTACCCGGTAGGAAGGCCGAT |

**Table S5.** Releasers for short jacks.

| Name | Sequence (5'→ 3') |
|---|---|
| r_13j22.1 | TTTTATTACTGAAAACACTTCTCAGGATTCTGAGAATGTCCCTCGGTCAA |
| r_13j22.2 | CACTGATTATAGTCGTGTGACTGGTGAATCTGTCGGTGGCCTCAATGGTC |
| r_13j22.3 | TAATCCATTTCGTGATGGACAGACTCTTTTACCCAATGTAAATATGCGCA |
| r_62j31.1 | GTGTTTATTCTTTTCAAGGATTCTAAGGGAAATATGATTCCGTGGTCAAC |
| r_62j31.2 | TATCGCTATGTTATTTAACGCCTTATTTATCATAATCTAAGCCGGATTAC |
| r_62j31.3 | ATTTCAAACCATATTGACTCTTCTCAGCGTCTCACGGTCGGTATCGCAGT |
| r_11j33.1 | TGGTATTCCTAATCCGGTTCTTGTCAAGATTATAGTTGAATGTCAGCAGT |
| r_11j33.2 | GGGTAATGAATAATCTCAACTGATGAATCTTTACGTTGATTTGTCAGATC |
| r_11j33.3 | TAATGTTGTTCCACTGAATGAGCAGCTTTGTTCTACCTGTAATAGCATGC |
| r_73j23.1 | GCAAAATCCCATTCCGGGCTATACTTATATCAAAGTTGTTTATCTGCGAA |
| r_73j23.2 | TGATACACCTATACAGAAAATTCATTTACTAACTGAGTACGGAATCGGTC |
| r_73j23.3 | GACGACAAAACAGGGTGGCGGTACTAAACCTCCGTCTGGAAATTAGCAGC |
| r_81j42.1 | TTCTGATTATCGTTCTAAAAATTTTTATCCTTTTTGGGGCTTCTGACATG |
| r_81j42.2 | AATATATGAGGAACCGGGGTACATATGATTGATTGCATTTAAGTAGACCT |
| r_81j42.3 | TTACGATTACCTACCTACACATTACTCAGGCACATGCTAGTTCCAGTAGT |
| r_12j41.1 | CAATGCTGGCGGGTGACGTTTCCGGCCTTGCTAACCTCCTGTCTGCAGAT |
| r_12j41.2 | ATGGTTTCATTGCGGCTCTGGTGGTGGTTCTGGCTGCTATCGCAGTCATG |
| r_12j41.3 | TGAGGGTGGTGTCTGTCGCTACTGATTACGGTGTGGCGGCTCGGTCAATC |
| r_63j21.1 | CGCGTCAACCTCGGCTGGTGCCCGTACATTGCACAGTAGGGTATGGTCAC |
| r_63j21.2 | GAGGTTAACAGGTGCAAGAAAAAGCTAGAAGAGAGCTTCAGTTATGCAGC |
| r_63j21.3 | GATAATTACAGAAGTACGGTTCAACGATTGCTGCTGAATTCTTTCAGAGC |
| r_61j53.1 | GCTTCATCGGGGATGCTAACACCCAGTAGATGGATTTTATACATATGGCC |
| r_61j53.2 | GCTCAGAGTAAAATCACTGCCCGACAATTGCTAGGTCTATTCTAGTCGCA |
| r_61j53.3 | TAATTTCGACGCCTTTATCCTTGAGGCAGATCTTCGTGTTAACGTTGACA |
| r_72j51.1 | CTGGCTTCAGCCAGCTTGGAGCGAACGACCTACCGAAGGTAATGACAGCT |
| r_72j51.2 | TGCACACAGCCAGAGCGCAGATACCAAATACTGGGGGGTTCGGTGCCTAA |

| | |
|---|---|
| r_72j51.3 | TGTAGCCGTAGGGCGCAGCGGTCGGGCTGAACGTTCTTCTAGTAATCGCG |
| r_82j32.1 | ATACAGGCTGCCTATGGGAAATGTGGGAGCTACTTTCTGGTGCTCTAGAG |
| r_82j32.2 | TCTTCCGGATCAACGTACTAGAGTTCATGATCTCCCGAGTAACAGTCTGA |
| r_82j32.3 | CACCACCGTGTGATGGCACAATGGCAATCCCATTGCCCTGGTCACAGTTG |
| r_71j43.1 | TGGATGGAGGCAGCATTGGTAACTGTCAGACCATTAATAGACCTAGTAGC |
| r_71j43.2 | CTCACTGATTAGGATAAAGTTGCAGGACCACTAGATAGGTGCTAATCGCG |
| r_71j43.3 | GCCCTTCCGGCAACGAAATAGACAGATCGCTGTCTGCGCTCGTGACAGCT |
| r_83j52.1 | ACTGCGTTGATGCTATGTGAGTTGCACCGCGGTCAGACCAGCGCAGTCAT |
| r_83j52.2 | GATCGTCCACACGTTTGTATGTGCTACCGCTGGGGCCTTTGGGAATCCTG |
| r_83j52.3 | ATCGGCCTTCCTACCGGGTACTGGATCATCAACCTGGAATCCCAACGGTT |

**Table S6.** Long jack staples.

| Name | Sequence (5'→ 3') |
|---|---|
| 4383_1 | GTGCTTGGCTCACATAGCGCCTCCATCCAGTCTATTAATCCGCGGTGCAA |
| 4383_2 | AGTCAATCAACTTTATCCTGTGGACGATCCCAAAGGCCCAGTGGTCCTGC |
| 4383_3 | GCATGTAGCCGAGCGCAGATTGATGATCCA |
| 4383_4 | GATGGTCTGCCGGAAGGGGTACCCGGTAGGGATTCCAGGTAAACCAGCCA |
| 4383_5 | CCAGACACCATACAAACGTAATCAGTGAGTTATCAGCAACAGCGGTAGCA |
| 4383_6 | CGGGGCACTACCAATGCTATCAACGCAGTGCTGGTCTGAGGTCTGACAGT |
| 3282_1 | CAGTTTTCTTTCCCATAGCCCGATGAAGCGTATAAAATCTAGCTCCCACA |
| 3282_2 | ACGCCTAAGGCAGTGATTGATCCGGAAGATTACTCGGGAAGCAATTGTCG |
| 3282_3 | AGCATAAAATTAACACGAATGGGATTGCCA |
| 3282_4 | CGACGAGCCGTCGAAATTTTGTGCCATCAACCAGGGCAACAGCACCGTTA |
| 3282_5 | AGTCATGACTAGTACGTTTTACTCTGAGCCGATGTTAGAGATCATGAACT |
| 3282_6 | AAGTCTTATGTTAGCATCGCAGCCTGTATCACCAGAAAGCATCTACTGGG |
| 2172_1 | GTACTGGAGCACCAGCCGGCTGAAGCCAGTTACCTTCGGGCAATGTACGG |
| 2172_2 | CGTGCCGTTCTGCGCTCTCTGTTAACCTCACTGAAGCTCAGTATTTGGTA |
| 2172_3 | TTCACTGCACTAGAAGAACAGCAATCGTTG |
| 2172_4 | GAATAATACTACGGCTACAACCGTACTTTAGAATTCAGCTGGTGGCCTAA |
| 2172_5 | CCTGGAGCTTTCTTGCACGGCTGTGTGCAGTTCTTGAAGTCTTCTAGCTT |
| 2172_6 | TGTACCGTCTCCAAGCTGAGGTTGACGCGACCCTACTGTTAGGTCGTTCG |
| 1273_1 | AATGACATAAACGTCACCTGGGATTTTGCTAAACAACTTAGCAAGGCCGG |
| 1273_2 | ACTCATCAATTTTCTGTAAATGAAACCATCGATAGCAGCTTAGTAAATGA |
| 1273_3 | TTAAATTTCTTTCCAGACGACCGTAATCAG |

| | |
|---|---|
| 1273_4 | TCTCTACAGTTTTGTCGTTAGCGACAGAAGAGCCGCCACAACGATCTAAA |
| 1273_5 | GCCAAACGCCAGAGCCGCTAGGTGTATCATAGTTAGCGTCAGAACCACCA |
| 1273_6 | ACCAAGTGTAGCCCGGAACGCCAGCATTGACAGGAGGTTTGATATAAGTA |
| 1181_1 | CATTTCCAAGAACCGGATGATAATCAGAAAAGCCCCAAATAATCTTGACA |
| 1181_2 | GGGAGCGCTACCCCGGTTATTCATTACCCAAATCAACGTTCAATCATATG |
| 1181_3 | GATGGAGAAAACTAGCATGAACAAAGCTGC |
| 1181_4 | TTCATTCTGGTAATCGTATCATTCAGTGATTACAGGTAGAATCGATGAAC |
| 1181_5 | CTCGCCAGAGTTGAGATTCCTCATATATTCAAACAAGAGAAAGATTCATC |
| 1181_6 | CACTGTAATTTTTAGAACTAGGAATACCACATTCAACTAAAGGATAAAAA |
| 1362_1 | TAGGCACTGAAGTGTTTTAGAATAAACACCGGAATCATACAGAATCCTGA |
| 1362_2 | AAAAGAGTGCGTTAAATATATAATCAGTGAGGCCACCGATGATAAATAAG |
| 1362_3 | GATGATAATACCGACCGTGGTAAAAGAGTC |
| 1362_4 | TCATGAGTTGGTTTGAAATGTCCATCACGTTTACATTGGCCTAAATTTAA |
| 1362_5 | GCCGCGCTGTCACACGACACATAGCGATACATCTTCTGACAGATTCACCA |
| 1362_6 | AAGGTGGTATCCTTGAAACAGTAATAAAAGGGACATTCTTTTCCCTTAGA |

**Table S7.** Releasers for long jacks

| Name | Sequence (5'→ 3') |
|---|---|
| r_1273_1 | AATGACATAAACGTCACCTGGGATTTTGCTAAACAACTTAGCAAGGCCGG |
| r_1273_2 | ACTCATCAATTTTCTGTAAATGAAACCATCGATAGCAGCTTAGTAAATGA |
| r_1273_3 | TTAAATTTCTTTCCAGACGACCGTAATCAG |
| r_1273_4 | TCTCTACAGTTTTGTCGTTAGCGACAGAAGAGCCGCCACAACGATCTAAA |
| r_1273_5 | GCCAAACGCCAGAGCCGCTAGGTGTATCATAGTTAGCGTCAGAACCACCA |
| r_1273_6 | ACCAAGTGTAGCCCGGAACGCCAGCATTGACAGGAGGTTTGATATAAGTA |
| r_1181_1 | CATTTCCAAGAACCGGATGATAATCAGAAAAGCCCCAAATAATCTTGACA |
| r_1181_2 | GGGAGCGCTACCCCGGTTATTCATTACCCAAATCAACGTTCAATCATATG |
| r_1181_3 | GATGGAGAAAACTAGCATGAACAAAGCTGC |
| r_1181_4 | TTCATTCTGGTAATCGTATCATTCAGTGATTACAGGTAGAATCGATGAAC |
| r_1181_5 | CTCGCCAGAGTTGAGATTCCTCATATATTCAAACAAGAGAAAGATTCATC |
| r_1181_6 | CACTGTAATTTTTAGAACTAGGAATACCACATTCAACTAAAGGATAAAAA |
| r_1362_1 | TAGGCACTGAAGTGTTTTAGAATAAACACCGGAATCATACAGAATCCTGA |
| r_1362_2 | AAAAGAGTGCGTTAAATATATAATCAGTGAGGCCACCGATGATAAATAAG |
| r_1362_3 | GATGATAATACCGACCGTGGTAAAAGAGTC |
| r_1362_4 | TCATGAGTTGGTTTGAAATGTCCATCACGTTTACATTGGCCTAAATTTAA |

| | |
|---|---|
| r_1362_5 | GCCGCGCTGTCACACGACACATAGCGATACATCTTCTGACAGATTCACCA |
| r_1362_6 | AAGGTGGTATCCTTGAAACAGTAATAAAAGGGACATTCTTTTCCCTTAGA |
| r_4383_1 | GTGCTTGGCTCACATAGCGCCTCCATCCAGTCTATTAATCCGCGGTGCAA |
| r_4383_2 | AGTCAATCAACTTTATCCTGTGGACGATCCCAAAGGCCCAGTGGTCCTGC |
| r_4383_3 | GCATGTAGCCGAGCGCAGATTGATGATCCA |
| r_4383_4 | GATGGTCTGCCGGAAGGGGTACCCGGTAGGGATTCCAGGTAAACCAGCCA |
| r_4383_5 | CCAGACACCATACAAACGTAATCAGTGAGTTATCAGCAACAGCGGTAGCA |
| r_4383_6 | CGGGGCACTACCAATGCTATCAACGCAGTGCTGGTCTGAGGTCTGACAGT |
| r_3282_1 | CAGTTTTCTTTCCCATAGCCCGATGAAGCGTATAAAATCTAGCTCCCACA |
| r_3282_2 | ACGCCTAAGGCAGTGATTGATCCGGAAGATTACTCGGGAAGCAATTGTCG |
| r_3282_3 | AGCATAAAATTAACACGAATGGGATTGCCA |
| r_3282_4 | CGACGAGCCGTCGAAATTTTGTGCCATCAACCAGGGCAACAGCACCGTTA |
| r_3282_5 | AGTCATGACTAGTACGTTTTACTCTGAGCCGATGTTAGAGATCATGAACT |
| r_3282_6 | AAGTCTTATGTTAGCATCGCAGCCTGTATCACCAGAAAGCATCTACTGGG |
| r_2172_1 | GTACTGGAGCACCAGCCGGCTGAAGCCAGTTACCTTCGGGCAATGTACGG |
| r_2172_2 | CGTGCCGTTCTGCGCTCTCTGTTAACCTCACTGAAGCTCAGTATTTGGTA |
| r_2172_3 | TTCACTGCACTAGAAGAACAGCAATCGTTG |
| r_2172_4 | GAATAATACTACGGCTACAACCGTACTTTAGAATTCAGCTGGTGGCCTAA |
| r_2172_5 | CCTGGAGCTTTCTTGCACGGCTGTGTGCAGTTCTTGAAGTCTTCTAGCTT |
| r_2172_6 | TGTACCGTCTCCAAGCTGAGGTTGACGCGACCCTACTGTTAGGTCGTTCG |

**Table S8.** Photocleavable jack staples (PC: photocleavable spacer).

| Name | Sequence (5'→ 3') |
|---|---|
| 13j22.1_PC | GGGACATTCT [PC] CAGAATCCTGAGAAGTGTTTT [PC] CAGTAATAAAA |
| 13j22.2_PC | AGGCCACCGA [PC] CAGATTCACCAGTCACACGAC [PC] TATAATCAGTG |
| 13j22.3_PC | TTTACATTGG [PC] GTAAAAGAGTCTGTCCATCAC [PC] GAAATGGATTA |
| 13j22.4_PC | GTTGTAGCAA [PC] CATTTTGACGCTCAATCGTCT [PC] GCAAATTAACC |
| 62j31.1_PC | CGGAATCATA [PC] TTTCCCTTAGAATCCTTGAAA [PC] AGAATAAACAC |
| 62j31.2_PC | GCTTAGATTA [PC] TGATAAATAAGGCGTTAAATA [PC] ACATAGCGATA |
| 62j31.3_PC | ACCGACCGTG [PC] AGACGCTGAGAAGAGTCAATA [PC] TGGTTTGAAAT |
| 62j31.4_PC | AAAATCATAG [PC] CATCTTCTGACCTAAATTTAA [PC] GTGAATTTATC |
| 11j33.1_PC | CATTCAACTA [PC] TAATCTTGACAAGAACCGGAT [PC] TAGGAATACCA |
| 11j33.2_PC | AAATCAACGT [PC] AAAGATTCATCAGTTGAGATT [PC] ATTCATTACCC |
| 11j33.3_PC | TTACAGGTAG [PC] AACAAAGCTGCTCATTCAGTG [PC] GAACAACATTA |

| | |
|---|---|
| 11j33.4_PC | CCCTGACGAG [PC] CGTTAATAAAACGAACTAACG [PC] AATAAGGCTTG |
| 73j23.1_PC | TAAACAACTT [PC] TGATATAAGTATAGCCCGGAA [PC] TGGGATTTTGC |
| 73j23.2_PC | CCGTACTCAG [PC] TTAGTAAATGAATTTTCTGTA [PC] TAGGTGTATCA |
| 73j23.3_PC | TTTCCAGACG [PC] GAGGTTTAGTACCGCCACCCT [PC] GTTTTGTCGTC |
| 73j23.4_PC | CCCTCAGAAC [PC] TAGTTAGCGTAACGATCTAAA [PC] CAGAACCGCCA |
| 81j42.1_PC | AAGCCCCAAA [PC] AAGGATAAAAATTTTTAGAAC [PC] GATAATCAGAA |
| 81j42.2_PC | TTAAATGCAA [PC] TCAATCATATGTACCCCGGTT [PC] CCTCATATATT |
| 81j42.3_PC | AACTAGCATG [PC] TGCCTGAGTAATGTGTAGGTA [PC] GGTAATCGTAA |
| 81j42.4_PC | GGGTGAGAAA [PC] CAAACAAGAGAATCGATGAAC [PC] AAGATTCAAAA |
| 12j41.1_PC | ACAGGAGGTT [PC] AGCAAGGCCGGAAACGTCACC [PC] CGCCAGCATTG |
| 12j41.2_PC | CGATAGCAGC [PC] CAGAACCACCACCAGAGCCGC [PC] AATGAAACCAT |
| 12j41.3_PC | GAGCCGCCAC [PC] ACCGTAATCAGTAGCGACAGA [PC] CACCACCCTCA |
| 12j41.4_PC | CTTTAGCGTC [PC] CAGAACCGCCACCCTCAGAGC [PC] ATCAAGTTTGC |
| 63j21.1_PC | ACCCTACTGT [PC] GCAATGTACGGGCACCAGCCG [PC] AGGTTGACGCG |
| 63j21.2_PC | ACTGAAGCTC [PC] TCTTCTAGCTTTTTCTTGCAC [PC] CTGTTAACCTC |
| 63j21.3_PC | AGAATTCAGC [PC] AGCAATCGTTGAACCGTACTT [PC] CTGTAATTATC |
| 63j21.4_PC | ATCATTCCTT [PC] GGGGTTAATCGGTGTATGGGT [PC] TATGGGCTGCG |
| 61j53.1_PC | GTATAAAATC [PC] CATCTACTGGGTGTTAGCATC [PC] CCCGATGAAGC |
| 61j53.2_PC | GAATAGACCT [PC] AGCAATTGTCGGGCAGTGATT [PC] TTACTCTGAGC |
| 61j53.3_PC | TTAACACGAA [PC] GATCTGCCTCAAGGATAAAGG [PC] CGTCGAAATTA |
| 61j53.4_PC | CTTATGCTCA [PC] CGATGTTAGACAGCACCGTTA [PC] CTGGCTGACAC |
| 72j51.1_PC | TTACCTTCGG [PC] TAGGTCGTTCGCTCCAAGCTG [PC] GCTGAAGCCAG |
| 72j51.2_PC | CGAACCCCCC [PC] AGTATTTGGTATCTGCGCTCT [PC] GGCTGTGTGCA |
| 72j51.3_PC | CTAGAAGAAC [PC] GTTCAGCCCGACCGCTGCGCC [PC] CTACGGCTACA |
| 72j51.4_PC | CTATCGTCTT [PC] GTTCTTGAAGTGGTGGCCTAA [PC] TTATCCGGTAA |
| 82j32.1_PC | CACCAGAAAG [PC] TAGCTCCCACATTTCCCATAG [PC] GCAGCCTGTAT |
| 82j32.2_PC | TTACTCGGGA [PC] GATCATGAACTCTAGTACGTT [PC] GATCCGGAAGA |
| 82j32.3_PC | ACCAGGGCAA [PC] TGGGATTGCCATTGTGCCATC [PC] ACACGGTGGTG |
| 82j32.4_PC | TTCAACCGTT [PC] CTGCGCATATTGTCAACTTCG [PC] AGGCGGGAGGA |
| 71j43.1_PC | GTCTATTAAT [PC] GGTCTGACAGTTACCAATGCT [PC] GCCTCCATCCA |
| 71j43.2_PC | GCACCTATCT [PC] AGTGGTCCTGCAACTTTATCC [PC] TAATCAGTGAG |
| 71j43.3_PC | CGAGCGCAGA [PC] CAGCGATCTGTCTATTTCGTT [PC] GCCGGAAGGGC |
| 71j43.4_PC | GCCTGACTCC [PC] TTATCAGCAATAAACCAGCCA [PC] CATCCATAGTT |

| | |
|---|---|
| 83j52.1_PC | GCTGGTCTGA [PC] CCGCGGTGCAACTCACATAGC [PC] ATCAACGCAGT |
| 83j52.2_PC | CCAAAGGCCC [PC] CAGCGGTAGCACATACAAACG [PC] TGTGGACGATC |
| 83j52.3_PC | GGATTCCAGG [PC] TTGATGATCCAGTACCCGGTA [PC] GGAAGGCCGAT |
| 83j52.4_PC | TATCAGTCTT [PC] ACCGAACCCCAGACAGGGGCT [PC] GGGGAACGCTA |

**Table S9.** Oligonucleotides for functionalizing DNA origami (cy 5 ●, cy 5.5 ●, thiol ●, PC, cholesterol, and **spacers**).

| Name | Sequence (5'→ 3') |
|---|---|
| Oligo labeled with cy 5 ● | [cy 5] GCGGTTCAGA |
| Oligo labeled with cy 5.5 ● | [cy 5.5] TCAGCAGC |
| Thiolated Oligo ● | [SH] TGGATCAACCGGGTCTCA |
| Oligo labeled with PC spacers | GCTGCTGAT**TTT**GAGACCCGGTTGATCCA**TTT** [PC] AGGGCCATCG |
| Oligo labeled with cholesterol moieties | TCTGCTCATGCAGAG [chol] |

### S15. Supplementary References

1. Engelhardt, F. A.; Praetorius, F.; Wachauf, C. H.; Brüggenthies, G.; Kohler, F.; Kick, B.; Kadletz, K. L.; Pham, P. N.; Behler, K. L.; Gerling, T.; Dietz, H., Custom-size, functional, and durable DNA origami with design-specific scaffolds. *ACS Nano* **2019,** *13*, 5015-5027.
2. Liu, B.; Liu, J., Freezing directed construction of bio/nano interfaces: reagentless conjugation, denser spherical nucleic acids, and better nanoflares. *Journal of the American Chemical Society* **2017,** *139*, 9471-9474.
3. Elani, Y.; Trantidou, T.; Wylie, D.; Dekker, L.; Polizzi, K.; Law, R. V.; Ces, O., Constructing vesicle-based artificial cells with embedded living cells as organelle-like modules. *Scientific Reports* **2018,** *8*, 4564.
4. Edmondson, A. C., A Fuller explanation: the synergetic geometry of R. Buckminster Fuller. *Boston: Birkhauser Verlag* **1987**.
5. Verheyen, H. F., The complete set of Jitterbug transformers and the analysis of their motion. *Computers Mathematics with Applications* **1989,** *17*, 203-250.
6. Gray, R. W., The Jitterbug motion. http://www.rwllY.Projects.comlrbfnotes/toc.html: **2002**.
7. Zhang, F.; Jiang, S.; Wu, S.; Li, Y.; Mao, C.; Liu, Y.; Yan, H., Complex wireframe DNA origami nanostructures with multi-arm junction vertices. *Nature Nanotechnology* **2015,** *10*, 779-784.
8. Dirks, R. M.; Lin, M.; Winfree, E.; Pierce, N. A., Paradigms for computational nucleic acid design. *Nucleic Acids Research* **2004,** *32*, 1392-1403.
9. Sample, M.; Liu, H.; Diep, T.; Matthies, M.; Sulc, P., Hairygami: Analysis of DNA nanostructures' conformational change driven by functionalizable overhangs. *ACS Nano* **2024,** *18*, 30004-30016.
10. Rovigatti, L.; Šulc, P.; Reguly, I. Z.; Romano, F., A comparison between parallelization approaches in molecular dynamics simulations on GPUs. *Journal of Computational Chemistry* **2015,** *36*, 1-8.
11. DeLuca, M.; Duke, D.; Ye, T.; Poirier, M.; Ke, Y.; Castro, C.; Arya, G., Mechanism of DNA origami folding elucidated by mesoscopic simulations. *Nature Communications* **2024,** *15*, 3015.
12. Bellot, G.; McClintock, M. A.; Lin, C.; Shih, W. M., Recovery of intact DNA nanostructures after agarose gel–based separation. *Nature Methods* **2011,** *8*, 192-194.
13. Heras, J.; Domínguez, C.; Mata, E.; Pascual, V.; Lozano, C.; Torres, C.; Zarazaga, M., GelJ–a tool for analyzing DNA fingerprint gel images. *BMC Bioinformatics* **2015,** *16*, 270.
14. Sun, Y.; Rombola, C.; Jyothikumar, V.; Periasamy, A., Förster resonance energy transfer microscopy and spectroscopy for localizing protein–protein interactions in living cells. *Cytometry Part A* **2013,** *83*, 780-793.
15. Mátyus, L., Fluorescence resonance energy transfer measurements on cell surfaces. A spectroscopic tool for determining protein interactions. *Journal of Photochemistry and Photobiology. B, Biology* **1992,** *12*, 323-337.
16. Jares-Erijman, E. A.; Jovin, T. M., FRET imaging. *Nature Biotechnology* **2003,** *21*, 1387-1395.
17. Hoppe, A.; Christensen, K.; Swanson, J. A., Fluorescence resonance energy transfer-based stoichiometry in living cells. *Biophysical Journal* **2002,** *83*, 3652-3664.
18. Berney, C.; Danuser, G., FRET or no FRET: a quantitative comparison. *Biophysical Journal* **2003,** *84*, 3992-4010.
19. Giordano, L.; Jovin, T. M.; Irie, M.; Jares-Erijman, E. A., Diheteroarylethenes as thermally stable photoswitchable acceptors in photochromic fluorescence resonance energy transfer (pcFRET). *Journal of the American Chemical Society* **2002,** *124*, 7481-7489.
20. Zhang, X.; Servos, M. R.; Liu, J., Instantaneous and quantitative functionalization of gold nanoparticles with thiolated DNA using a pH-assisted and surfactant-free route. *Journal of the American Chemical Society* **2012,** *134*, 7266-7269.
21. Hurst, S. J.; Lytton-Jean, A. K.; Mirkin, C. A., Maximizing DNA loading on a range of gold nanoparticle sizes. *Analytical Chemistry* **2006,** *78*, 8313-8318.
